# Metal Coordination Dynamics Governs Selective Epoxidation in Hyoscyamine 6β-Hydroxylase: Integrated Experimental and Computational Insights

**DOI:** 10.64898/2026.06.01.729211

**Authors:** Jinyan Zhang, Lian Wu, Siqi Wu, Xiao Liu, Lina Dong, Eliott S. Wenger, Ridao Chen, Carsten Krebs, Alexey Silakov, Amie K. Boal, J. Martin Bollinger, Jiahai Zhou, Binju Wang

## Abstract

Iron(II)- and 2-oxoglutarate-dependent (Fe(II)/2OG) enzymes catalyze a wide range of C–H bond activation and functionalization reactions and play essential roles in biosynthesis and metabolic regulation. Despite extensive mechanistic studies, the principles governing selectivity between canonical hydroxylation and alternative transformations remain incompletely understood. Here, we investigate the catalytic mechanism of hyoscyamine 6β-hydroxylase (H6H), a Fe(II)/2OG-dependent oxygenase that sequentially catalyzes the 6β-hydroxylation of hyoscyamine followed by 6,7-exo-epoxidation of 6β-hydroxyhyoscyamine to generate scopolamine. Combined molecular dynamics and QM/MM calculations reveal that an inline Fe(IV)-oxo intermediate initiates hydrogen atom abstraction from the substrate C7 position. The resulting Fe(III)–OH species subsequently deprotonates the substrate hydroxyl group in a process coupled to substrate coordination to the iron center and an in-line-to-off-line rearrangement of the Fe(III)–OH moiety. This coordination dynamics machinery is further supported by the observed chlorination reactivity on the same substrate. Importantly, this coordination switch favors epoxide formation over hydroxyl rebound, thereby directing the reaction toward selective epoxidation. Further computational analysis of the L290F variant demonstrates that steric constraints imposed by L290 are essential for suppressing hydroxylation, revealing a bidirectional regulatory mechanism governing epoxidation/hydroxylation selectivity. Whereas iron coordination dynamics promote epoxidation reactivity, precise substrate positioning and protein-derived steric effects suppress the competing hydroxylation pathway. These findings are consistent with available experimental observations and establish metal coordination dynamics as a key determinant of selective C–H functionalization in Fe(II)/2OG enzymes.

## INTRODUCTION

Iron(II)- and 2-oxoglutarate-dependent (Fe(II)/2OG) enzymes catalyze a diverse range of C−H bond activation and functionalization reactions.^1-9^ The well-established mechanism involves the 2OG-assisted activation of molecular oxygen by the Fe(II) cofactor, generating a highly reactive Fe(IV)-oxo intermediate that initiates subsequent transformations.^10-26^ The most well-studied case of C(*sp*^*3*^) hydroxylation begins with hydrogen atom transfer (HAT) from the substrate to the Fe(IV)-oxo species, resulting in the formation of an intermediate harboring a substrate-centered radical and an Fe(III)−OH center.^27-34^ Subsequent radical coupling between the OH ligand and the substrate carbon (OH-rebound) then yields the hydroxylated product.^35-40^ Remarkably, Fe(II)/2OG enzymes have also evolved to catalyze numerous alternative transformations—including desaturation,^41-47^ halogenation,^22, 48-72^ ring formation,^73-87^ epimerization^88-91^ and endoperoxidation^92-94^—that effectively bypass the thermodynamically favored hydroxylation pathway. Despite significant advances in understanding Fe(II)/2OG catalytic chemistry, the precise mechanistic determinants that divert reaction trajectories away from hydroxylation toward these alternative outcomes remain incompletely elucidated.

To address this fundamental question, we have undertaken a detailed mechanistic investigation of hyoscyamine 6β-hydroxylase (H6H),^78, 95-106^ an Fe(II)/2OG-dependent oxygenase that catalyzes 6β-hydroxylation of hyoscyamine (Hyo) to form 6β-hydroxyhyoscyamine (6-OH Hyo) followed by a 6,7-exo-epoxidation to yield the anticholinergic drug scopolamine (Sco) (Figure 1a). Structural and computational analysis of the hydroxylation step revealed that both Hyo carbons that the enzyme can target for HAT (C6 and C7) are positioned near the iron center but C6 adopts a more upright orientation that is more favorable, explaining the known regioselectivity in the hydroxylation step (Figure 1b & 1c).^103, 105^ The second step, involving 6,7-exo-epoxidation, is mechanistically more complex, requiring C7−H activation, deprotonation of the C6−OH group, and C7−O(C6) bond formation—each posing a significant energetic challenge.

**Figure 1.**
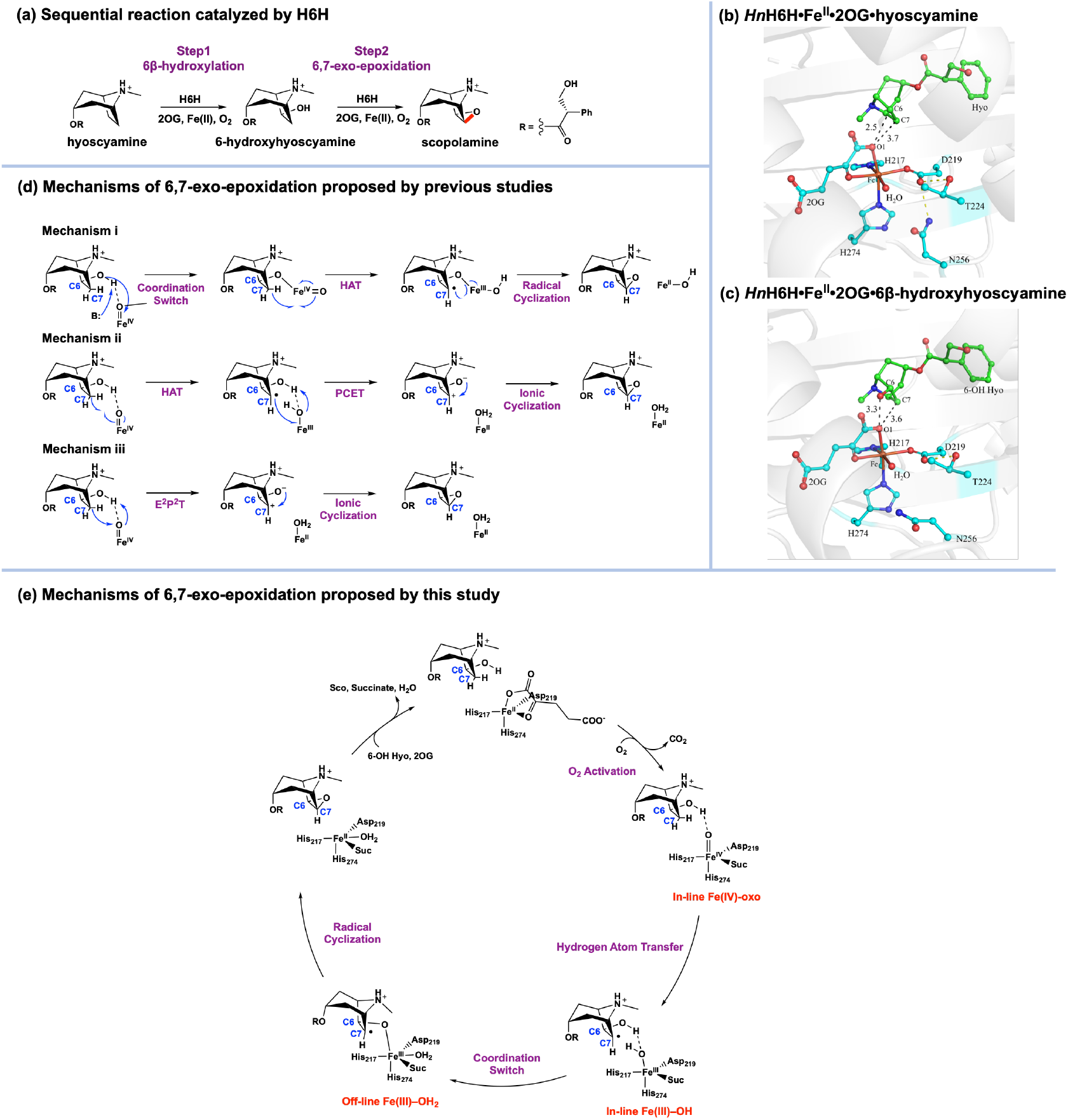
(a) Sequential reaction catalyzed by H6H. (b) Crystal structure of *Hn*H6H•Fe^II^•2OG•hyoscyamine (PDB ID: 8HV0). (c) Crystal structure of *Hn*H6H•Fe^II^•2OG•6β-hydroxyhyoscyamine (PDB ID: 8HRW). (d) Possible mechanisms of 6,7-exo-epoxidation catalyzed by H6H proposed by previous studies. (e) Mechanisms of 6,7-exo-epoxidation catalyzed by H6H proposed by this study.

The mechanistic determinants governing selectivity between hydroxylation and epoxidation in the second step remain unresolved. Ushimaru et al. proposed a mechanism in which the C6-OH group of the substrate directly coordinates to the Fe(IV)-oxo intermediate (Figure 1d, Mechanism i), thereby redirecting reactivity toward C7–H cleavage and subsequent C7–O bond formation.^101^ In the absence of such coordination, the oxo ligand would preferentially abstract a hydrogen from the C6 position. This hypothesis is consistent with observations in other radical-mediated C–X bond-forming reactions, such as in halogenases^22, 52, 53, 57, 59, 67, 71^ and the C–N bond-forming enzyme TqaL.^87^ Analogous mechanisms for C–O bond formation involving Fe–O-linked intermediates have also been proposed for HppE^107^ and LolO.^83^ Recent high-resolution crystallography and spectroscopic data suggested that the significant repositioning of the substrate’s tropane core needed to bring about (C6)–O coordination might not be feasible in H6H.^105^ Moreover, a significant increase in the 7β-hydroxylation/epoxidation product ratio in D_2_O suggested that deprotonation of the C6−OH group could be a key step in determining the fate of the reaction. Based on these findings, two plausible mechanisms were proposed for H6H-catalyzed epoxidation, both involving uncoordinated substrate: (1) a stepwise mechanism involving HAT followed by proton-coupled electron transfer (PCET) from the radical and subsequent polar cyclization of a high-energy zwitterionic intermediate (Figure 1d, Mechanism ii), and (2) a concerted pathway directly to the high-energy zwitterion (Figure 1d, Mechanism iii). Notably, consideration of the seemingly unlikely concerted pathway was driven by the observation of synergistic substrate and solvent deuterium kinetic isotope effects on decay of the Fe(IV)-oxo intermediate, which seemed inconsistent with any stepwise pathway for cleavage of the C7–H and (C6)O–H bonds.

Given the diversity of proposed mechanisms and possibility that substrate coordination during catalysis might have eluded detection by the crystallographic and spectroscopic approaches reported in prior work, we undertook an analysis by a complementary combination of molecular dynamics (MD) and QM/MM calculations informed by high-resolution crystallographic and spectroscopic data from earlier work.^105^ Indeed, our recent studies have emphasized the importance of mid-reaction iron coordination dynamics in modulating reactivity and selectivity in non-heme enzymes.^108^ For instance, our prior computational work on the carrier-protein-independent halogenase, BesD, implied that isomerization of the Fe(III)−OH intermediate formed by HAT is essential to prevent hydroxylation and ensure selective halogenation.^71^ In a more recent study, this mechanistic paradigm has been supported by crystallographic, spectroscopic, and additional DFT analysis.^109^ Building on these insights, we explored here the molecular mechanism of C−O bond formation in the second step of H6H catalysis (Figure 1e). Our study reveals a bidirectional regulatory mechanism that governs the partition between epoxidation and hydroxylation in this reaction. While the in-line-to-off-line shift of the Fe(III)–OH species is essential to drive the epoxidation outcome, the steric constraints imposed by L290 are critical for suppressing the hydroxylation by OH-rebound that might otherwise preempt epoxidation. Subsequent rational engineering of the H6H active site significantly enhanced its epoxidation efficiency, offering important implications for future biocatalytic studies and practical applications.

## RESULTS

### Coordination dynamics allows in-line-off-line Fe(IV)-oxo interconversion yet only in-line isomer drives substrate activation

Given that the mechanism of 2OG-assisted activation of molecular oxygen by the Fe(II) cofactor to generate the Fe(IV)-oxo reactive intermediate has been extensively studied, we begin our investigation directly from the Fe(IV)-oxo species. Based on the crystal structure of *Hn*H6H•Fe^II^•2OG•6-OH Hyo (Figure S1), we constructed the in-line Fe(IV)-oxo species, in which the oxo ligand is oriented toward the substrate. MD simulations (Figure 2a) show that the substrate maintains a stable conformation closely resembling that observed in the crystal structure (Figure 1c). In the in-line Fe(IV)-oxo configuration, the hydroxyl group of 6-OH Hyo forms a persistent and strong hydrogen bond with the oxo ligand, which is also evident in recent vanadyl crystal structures.^105^ Furthermore, the distance between the H(C7) of the substrate and the oxo moiety (∼2.6 Å) in QM/MM optimized structure supports the feasibility of HAT from this geometry (Figure 2b). To assess potential conformational flexibility of the Fe(IV)-oxo species,^57, 71, 108, 110-112^ we also examined the coordination isomerization of the Fe(IV)-oxo species from the in-line configuration to the off-line one. However, our QM/MM calculations indicate that the off-line Fe(IV)-oxo species is 7.1 kcal mol^−1^ higher in energy than its in-line counterpart (Figure S4), primarily due to the loss in the off-line configuration of the hydrogen bond between the oxo group and the substrate’s hydroxyl moiety. Further MD simulation of the off-line Fe(IV)-oxo species shows a significantly larger distance (∼5 Å) between C7–H and the oxo moiety, suggesting that this geometry is unlikely to be competent for the initiating HAT step (Figure S5). Notably, HAT from the C7 position mediated by the off-line Fe(IV)-oxo would first regenerate the in-line configuration, indicating that the off-line species is thermodynamically and kinetically poorly suited for substrate C7–H activation (Figure S6). These findings are consistent with our previous work and recent studies by Chang’s group.^22, 71, 113^ Starting from the off-line Fe(IV)-oxo, we also investigated substrate coordination to Fe, which is coupled to proton transfer to succinate, the only available base to accept the substrate’s alcohol proton. However, QM/MM scans indicate that this process is energetically unfavorable (Figure S7a). Moreover, the resulting six-coordination geometry is unfavorable for the subsequent HAT from C7 (Figure S7b). These findings disfavor Mechanism i (Figure 1d). In summary, our data suggest that the in-line Fe(IV)-oxo intermediate is the sole competent species for HAT from C7 of the substrate. Building upon these findings, we now turn to an examination of the reactivity of this in-line Fe(IV)-oxo species. Consistent with established literature precedent, all QM/MM computational analyses in these studies focus on the quintet ground state of the Fe(IV)-oxo intermediate.^11-16, 18, 19, 25, 27, 114^

**Figure 2.**
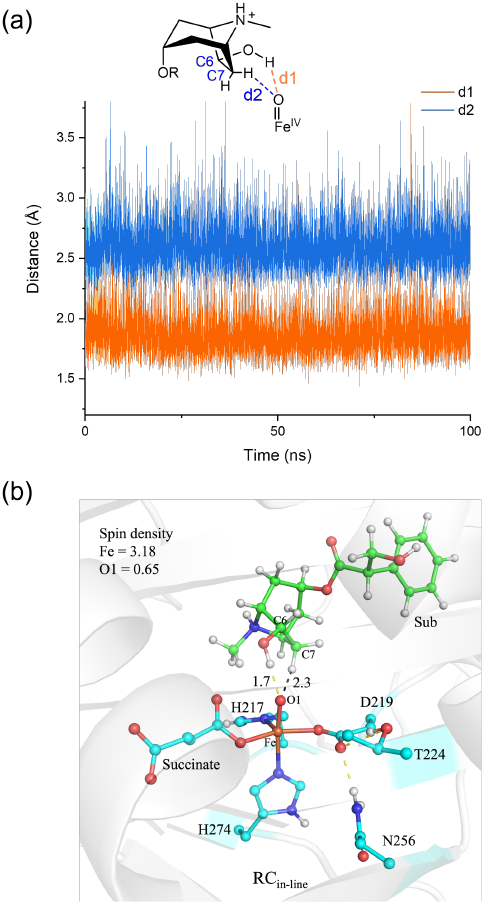
(a) Time evolution of the d1 and d2 distances from in-line conformation in 100 ns MD simulation. The distance between the ferryl O atom and hydroxyl H atom is denoted as d1, while the distance between the ferryl O atom and H atom on C7 is denoted as d2. (b) QM/MM-optimized structure of the Fe(IV)-oxo species. Key distances are given in Å.

### Coordination dynamics enables H6H-catalyzed selective 6,7-exo-epoxidation

Based on the MD-equilibrated structure of the in-line Fe(IV)-oxo species in the complex with 6-OH Hyo, we investigated the mechanism of 6,7-exo-epoxidation. Starting from the initial reactant complex (RC^in-line^), we first considered HAT from the hydroxyl group of 6-OH Hyo to the Fe(IV)-oxo species (Figure S8). However, our QM/MM calculations reveal that the resulting O-centered radical intermediate is 15.2 kcal mol^−1^ higher in energy than RC^in-line^. Furthermore, the subsequent C7–O6 bond formation, which is concerted with a hydrogen shift from C7 to the Fe(III)–OH moiety to yield the epoxide product, must overcome a high activation barrier of approximately 25.2 kcal mol^−1^ (Figure S9). These findings indicate that this reaction pathway is energetically unfavorable and can be excluded. We also tested the feasibility of the P^2^E^2^T mechanism (Figure 1d), but a QM/MM scan shows that the H atom from the OH group cannot be transferred in concert with the HAT from C7 (Figure S10). Thus, Mechanism iii is also not supported by our analysis (Figure 1d).

We next explored HAT from C7 to the Fe(IV)-oxo center (Figure 3a), which proceeds via a barrier of 21.9 kcal mol^−1^ (RC_in-line_ → IC1), leading to the formation of a C7-centered radical intermediate, IC1. From IC1, we examined several potential reaction pathways. First, we investigated the direct C7–O6 coupling accompanied by PCET from substrate to Fe(III)–OH (Figure S11); however, this pathway presents a barrier of ∼18.3 kcal mol^−1^ relative to IC1, thereby implying that mechanism ii is kinetically disfavored (Figure 1d). In addition, we evaluated the possibility of proton transfer from the substrate ammonium to E118,^106^ but this process was found to be unfeasible thermodynamically (Figure S12). Figure 3a illustrates the two alternative pathways that are more favorable energetically. The route depicted in red involves OH-rebound to form a dihydroxylation product (IC1 → PC_OH_), with an activation barrier of 15.6 kcal mol^−1^ relative to IC1. The other pathway involves proton transfer from the substrate hydroxyl group to the Fe(III)–OH moiety, which is coupled to the coordination of substrate O6 to Fe(III) and isomerization of the Fe(III)–OH moiety from an in-line to an off-line position (IC1 → IC2, Figure 3b). This step is associated with a barrier of 13.3 kcal mol^−1^ (IC1 → TS2). Starting from IC2, the subsequent C7–O6 coupling should be facile, with a calculated barrier of only 6.9 kcal mol^−1^ and an exothermicity of 14.6 kcal mol^−1^ (IC2 → PC).

**Figure 3.**
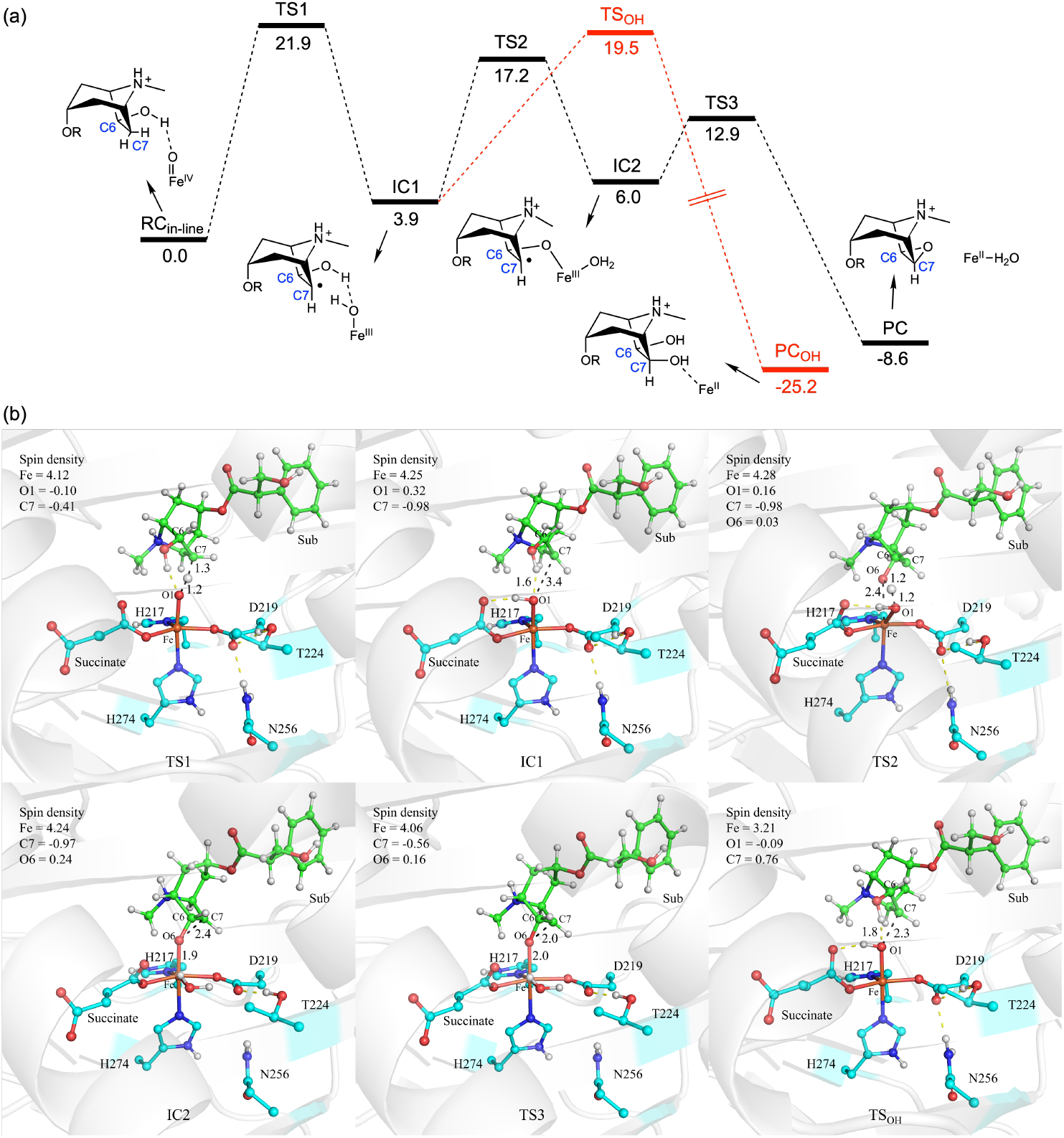
(a) QM(UB3LYP-D3/B2)/MM calculated energy profile (in kcal mol^-1^) for H6H-catalyzed 6,7-exo-epoxidation and 7β-hydroxylation. The dispersion corrections and ZPEs are included in the relative energies. (b) QM(UB3LYP-D3/B1)/MM optimized structures of key species involved in the reaction. Key distances are given in Å.

Thus, our QM/MM calculations show that epoxide formation via TS2 is preferred over 7β-hydroxylation via TS_OH_. The observed selectivity favoring epoxidation can be rationalized by the flexible coordination structure of the iron center. The substrate hydroxyl group maintains a robust hydrogen bond with the hydroxyl oxygen of the Fe(III)–OH complex, which effectively suppresses the OH-rebound pathway (TS_OH_). Moreover, the coordination of the substrate O6 atom to the iron center, which is coupled to movement of the original oxygen ligand from an in-line to an off-line position, enhances the acidity of the substrate C6-OH group and facilitates the proton transfer process. This combination of steps substantially stabilizes TS2 relative to TS_OH_ (Figure 3a). Interestingly, at the H6H active site, succinate exists in a “free” state, devoid of any stable hydrogen-bonding interactions with amino acid residues within the substrate binding pocket. As a consequence, in the IC1 state, succinate is able to engage in a hydrogen bond with the nascent Fe(III)–OH species (Figure 3b & Figure S13). Such a strong H-bond can greatly enhance the basicity of O1 (charge −0.75), thereby facilitating the subsequent proton transfer from the substrate OH group to Fe(III)–OH. In contrast, succinate in BesD is anchored by the second-sphere residue Asn219 through a hydrogen bond, precluding its interaction with Fe(III)–OH (Figure S13). As a result, the oxygen atom of Fe(III)–OH in BesD bears a markedly weaker negative charge of only −0.64. Notably, the calculated kinetic energy difference between the epoxidation route (TS2) and the hydroxylation route (TS_OH_) is 2.3 kcal mol^−1^, which corresponds to a epoxidation-to-hydroxylation product ratio of approximately 96:4, in excellent agreement with the experimental results reported by Wenger, et al.^105^

### Expansion of H6H reaction to chlorination further support the coordination dynamics mechinary

Since the in-line-to-off-line rearrangement of the Fe(III)– OH moiety is involved in epoxide formation by H6H, we asked whether H6H is also capable of performing a chlorination reaction on the native substrate of H6H. If so, this would not only further support the coordination dynamics machinery of H6H, but also expand its catalytic repertoire. Specifically, our previous studies showed that a post-HAT coordination switch is vital for selective chlorination in the carrier-protein-independent halogenase BesD, a finding that has since received experimental support.^71, 115^ More recently, our investigation of the carrier-protein-dependent halogenase SyrB2 also suggested that a post-HAT coordination switch of the Cl– Fe(III)–OH complex acts as a key step in enforcing selective halogenation.^116^ Motivated by these scenarios, we generated H6H variants lacking the canonical carboxylate ligand of the facial triad. ^117, 118^ Remarkably, the D219A and D219G substitutions produced chlorinated products from hyoscyamine substrates (Figure 4). As demonstrated in the halogenases BesD and SyrB2, the post-HAT coordination switch is critical for achieving selective halogenation. Without rearrangement of the Fe(III)–OH intermediate, chlorine rebound would be sterically inaccessible (Figure S14). Thus, the observed chlorination reactivity on hyoscyamine in H6H provides strong evidence that H6H possesses a coordination dynamics machinery for both epoxidation and chlorination.

**Figure 4.**
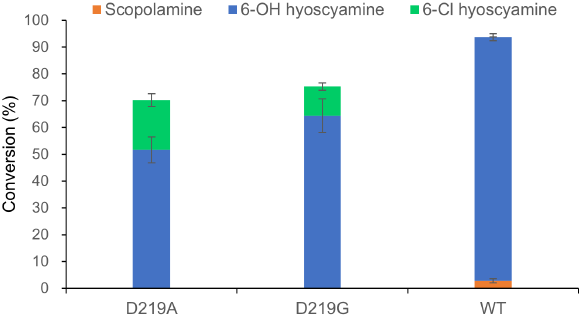
The percentage conversion of hyoscyamine to hydroxylation and chlorination products catalyzed by the mutants of *Hn*H6H.

### Steric effects in L290F impede the conformational change of Fe(III)–OH and facilitates OH-rebound

Recently, it was found that the introduction of phenylalanine (F) in place of L289 of *Ab*H6H (the corresponding residue at the equivalent position of *Hn*H6H is L290) shifts the outcome of the second step from primarily epoxidation to primarily hydroxylation.^105^ It was anticipated that the L289F substitution might introduce steric perturbations that alter the substrate-cofactor spatial arrangement. Statistical analyses of Fe(II)/2OG non-heme enzymes indicate that a larger Fe(IV)-oxo angle relative to the target hydrogen of the substrate favors hydroxylation, whereas a smaller angle promotes alternative reaction pathways.^53, 71, 119, 120^ To further validate our mechanistic proposal from the previous section, we conducted QM/MM calculations on the L290F *Hn*H6H variant (Figure 5). MD simulations and QM/MM geometry optimizations reveal that the replacement of the curved L290 by the flat F290 tilts the Fe=O bond towards the substrate, increasing the Fe–O–C7 angle to 127.8° in L290F, compared to 115.3° in the wild-type (WT) (Figure 6a). Notably, this angular shift exerts minor effects on the HAT from the substrate C7-H to the in-line Fe(IV)-oxo in L290F (21.6 kcal mol^-1^, TS1^L290F^). Starting from the Fe(III)–OH species of IC1^L290F^, both isomerization and OH-rebound pathways remain accessible. However, the OH-rebound pathway via TS_OH_^L290F^ (13.7 kcal mol^-1^) is kinetically favored over the isomerization pathway through TS2^L290F^ (17.2 kcal mol^-1^). For the Fe(III)–OH intermediate, we found the O–C7 distances and Fe–O–C7 angles to be very similar between the WT (3.4 Å and 105.2°) and L290F variant (3.6 Å and 105.8°), suggesting that the binding configuration of the substrate might not be the key factor in the enhanced C7 hydroxylation by the L290F variant (Figure 6b). However, our computational analysis suggests that steric hindrance by L290 specifically raises the OH-rebound barrier, and its replacement by F290 lowers the barrier for the OH-rebound process (Figure S15). In line with our previous analysis of the BesD reaction,^71^ our calculations on H6H imply that the OH-rebound process is dominated in both the WT enzyme and the L290 variant by migration of the substrate radical toward the Fe(III)–OH species rather than movement of the OH ligand toward the radical. This fact is evidenced from the minor Fe–O distance change in comparing IC1^L290F^ to TS_OH_^L290F^. Thus, our results collectively support that the steric hindrance caused by L290 is a major factor contributing to the elevated energy barrier for OH-rebound in the WT enzyme.

**Figure 5.**
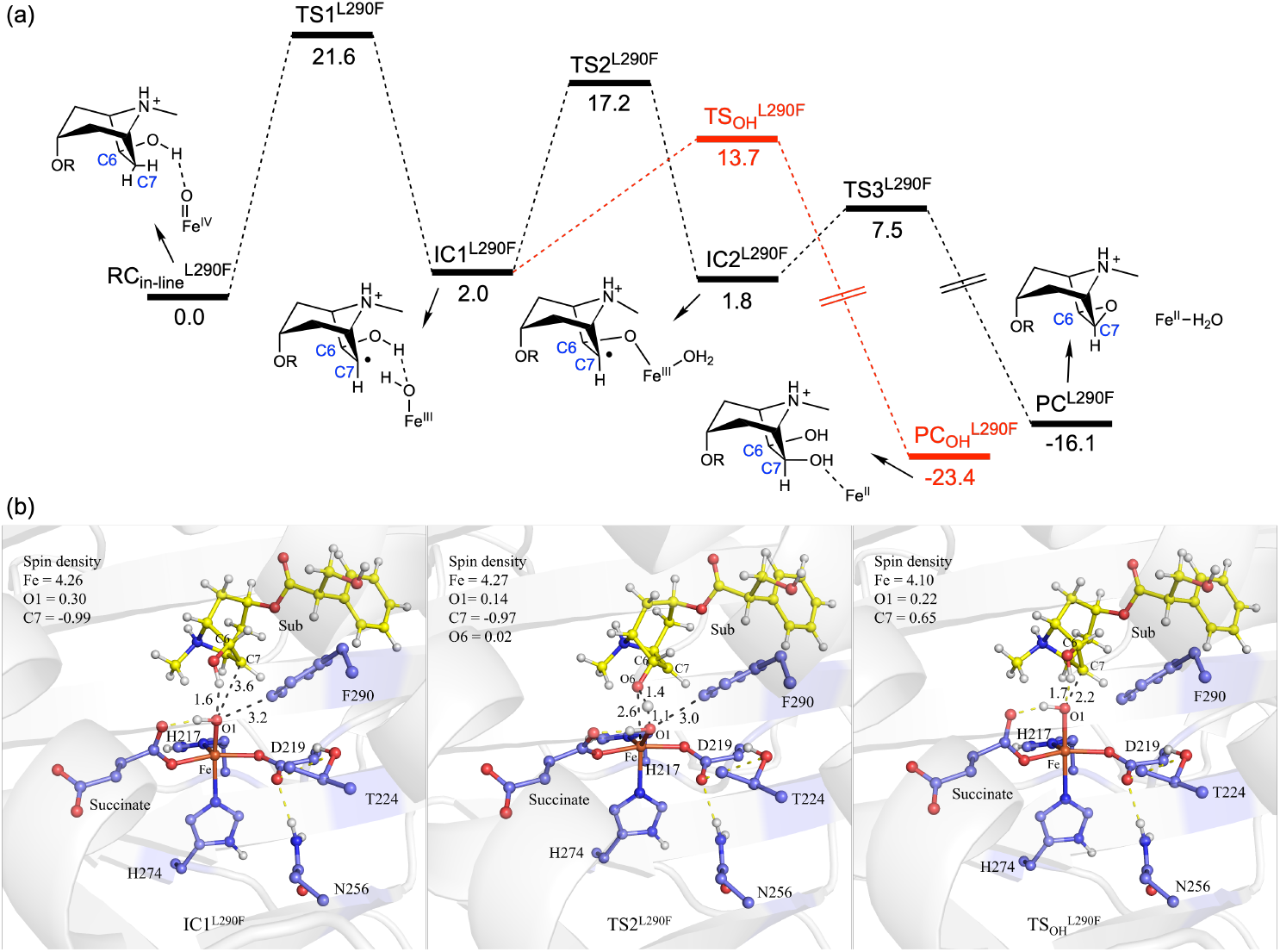
(a) QM/MM calculated energy profile (in kcal mol^-1^) for H6H (L290F)-catalyzed 6,7-exo-epoxidation and 7β-hydroxylation. The dispersion corrections and ZPEs are included in the relative energies. (b) QM(UB3LYP-D3/B1)/MM optimized structures of key species involved in the reaction. Key distances are given in Å.

**Figure 6.**
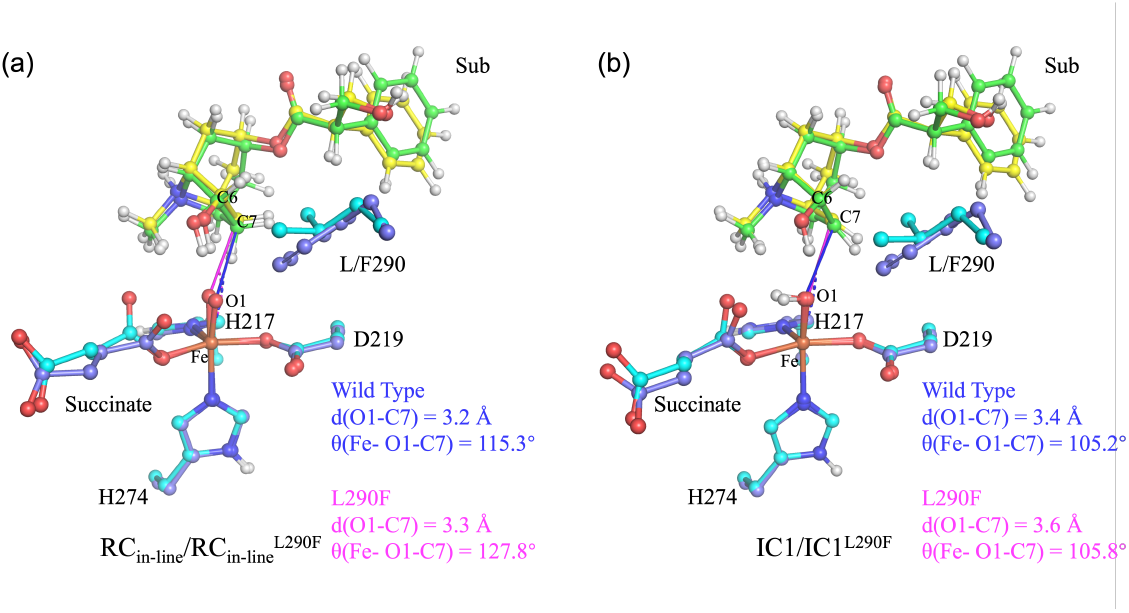
Structural superposition of (a) RC_in-line_/RC_in-line_^L290F^ and (b) IC1/IC1^L290F^. Carbon atoms in wild-type (active site) and wild-type (substrate) are shown in cyan and green, while carbon atoms in L290F (active site) and L290F (substrate) are shown in purple and yellow, respectively.

### Steric effects in rational engineering of H6H enhance 6,7-exo-epoxidation efficiency

The second 6,7-exo-epoxidation step catalyzed by H6H proceeds with significantly lower efficiency than the initial 6β-hydroxylation step (Figure 4 & Figure 7a).^78, 105^ Given the pivotal role of C–O cyclization reactions in biosynthetic pathways, the rational engineering of H6H to enhance its catalytic efficiency represents a promising strategy for synthetic biology applications. To systematically investigate structure-activity relationships in H6H, we designed a series of mutagenesis experiments targeting residues in both the first and second coordination spheres of the active site (Figure 7a). Our rationale was based on the well-established principle that metal-coordinating residues and second-sphere residues involved in non-covalent interactions often exert substantial influence on enzymatic activity and selectivity.^46, 63, 72, 119, 121-129^ For the Fe-coordinating residue D219, combined evidence from X-ray crystallography (Figure 1b & 1c) and MD simulations (Figure 2) reveals a conserved hydrogen-bonding network with second-sphere residues T224 and N256. This structural conservation underscores the catalytic essentiality of these residues. Interestingly, the D219E substitution led to a reduced overall conversion of Hyo to 6-OH Hyo and Sco, while simultaneously displaying enhanced epoxidation activity, thereby promoting Sco production. In contrast, the D219N variant exhibited complete loss of both 6β-hydroxylation activity (first step) and 6,7-exo-epoxidation (second step, using 6β-hydroxyhyoscyamine as the substrate). Notably, among second-sphere residues, the T224A substitution significantly improved 6,7-exo-epoxidation activity, achieving a 13.9-fold increase in epoxidation conversion from Hyo compared to the wild-type H6H, while preserving 6-OH Hyo accumulation at 60% of the wild-type level. Further enzyme kinetic studies using 6-OH Hyo as substrate demonstrated that the catalytic efficiency (*K*_cat_/*K*_M_) of the T224A variant in mediating 6,7-exo-epoxidation was approximately 12-fold higher than that of the wild-type enzyme (Table S3). In contrast, the N256A variant retained only 35% of 6-OH Hyo accumulation relative to the wild-type enzyme and showed no detectable epoxidation activity. More drastic effects were observed with N256S and N256D substitutions, which nearly abolished hydroxylation activity (>95% reduction) and completely eliminated epoxidation activity, highlighting the critical role of N256 in maintaining the iron coordination environment.

**Figure 7.**
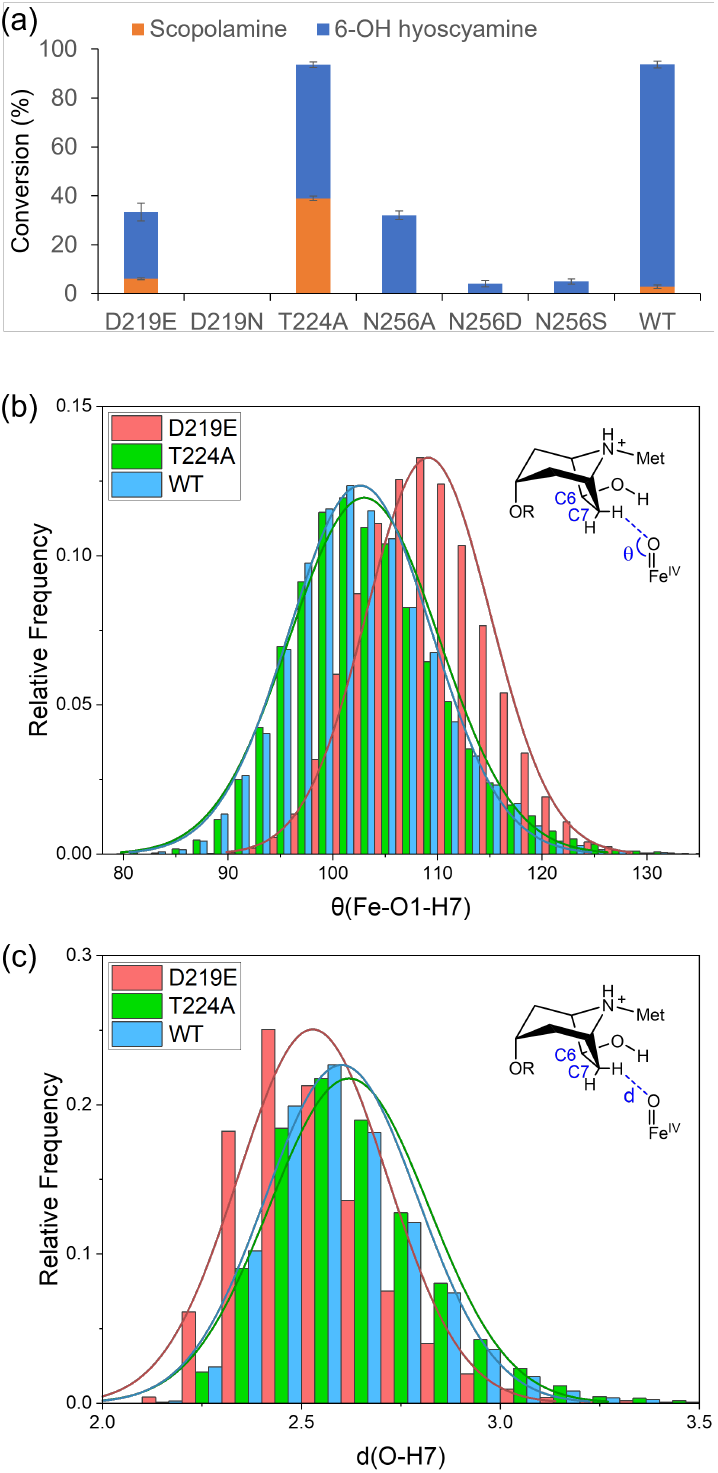
(a) Catalytic activity of wild-type (WT) and mutants of *Hn*H6H toward hyoscyamine. The total conversion was calculated as the sum of the individual conversion percentages of 6-hydroxyhyoscyamine (6-OH Hyo) and scopolamine (Sco) in the reaction. Data represent the mean ± S.D. from three independent replicates. (b) Time evolution of the angles Fe–O–H in WT, D219E and T224A. (c) Time evolution of the distances between the ferryl O atom and H(C7) atom in WT, D219E and T224A.

To elucidate the mechanistic basis for enhanced 6,7-exo-epoxidation in the D219E and T224A variants, we conducted MD simulations. The D219E variant exhibited a significantly enlarged Fe–O1–H7 angle compared to WT (Figure 7b), suggesting a shift toward a σ-pathway-dominated HAT mechanism, accompanied by a reduced HAT distance (Figure 7c). These structural changes provide a plausible explanation for the improved 6,7-exo-epoxidation kinetics while simultaneously rationalizing the reduced 6β-hydroxylation activity (Table S3). In contrast, the QM/MM calculations on the T224A variant showed that it should maintain an active site geometry nearly identical to that in the WT enzyme (Figure S16). Although the computed HAT energy barriers for the T224A and WT enzymes are similar, the variant exhibits a substantially higher imaginary frequency at the transition state for the HAT step (Figure S16), indicating an increased transmission coefficient and faster reaction (Table S3). In conclusion, our integrated experimental and computational approach identifies D219E and T224A as strategically valuable H6H variants for enhancing catalytic efficiency in scopolamine biosynthesis. These findings not only advance our mechanistic understanding of H6H but also provide a foundation for optimizing enzymatic C–O cyclization in synthetic biology applications.

## DISCUSSION

There is accumulating evidence that coordination dynamics play vital roles in controlling mechanistic pathways and outcomes in non-heme-iron enzymes, particularly in Fe(II)/2OG dependent enzymes.^108^ Previous spectroscopic and computational studies suggested that the coordination isomerization of the Fe(IV)-oxo species in the Fe(II)/2OG family could be facilitated by the unsaturated coordination environment of the iron center.^53, 110, 111^ For the TqaL-catalyzed C–H amination,^84, 85^ we found that the pre-HAT coordination switch of Fe(IV)-oxo species from in-line to off-line enables the coordination of substrate amino group (Figure 8a), thereby suppressing the kinetics of the unwanted hydroxylation reaction.^87^ In H6H, we found that substrate coordination is thermodynamically inaccessible (Figure S7). In addition, the off-line Fe(IV)-oxo species is incapable of abstracting a hydrogen atom from C7 of the substrate (Figure S5 & S6). Instead, all structural and computational evidence supports that only the in-line Fe(IV)-oxo species is responsible for HAT from the substrate C7–H site. Subsequently, the resulting Fe(III)–OH species acts as a base to deprotonate the substrate’s OH group. This deprotonation is coupled with the coordination of the substrate oxygen to the iron center and a configurational switch of the Fe(III)–OH moiety from the in-line to the off-line position (Figure 7b, H6H). Our analysis implies that this mechanistic sequence is critical to the H6H-catalyzed selective C–O coupling. Such a post-HAT coordination switch was first demonstrated in our earlier study of the carrier-protein-independent halogenase BesD (Figure 8b, BesD) and then further proved by our recent investigation of the carrier-protein-dependent halogenase SyrB2 (Figure 8b, SyrB2).^116^ This coordination dynamics machinery is further supported by the observed chlorination reactivity on the same substrate.

**Figure 8.**
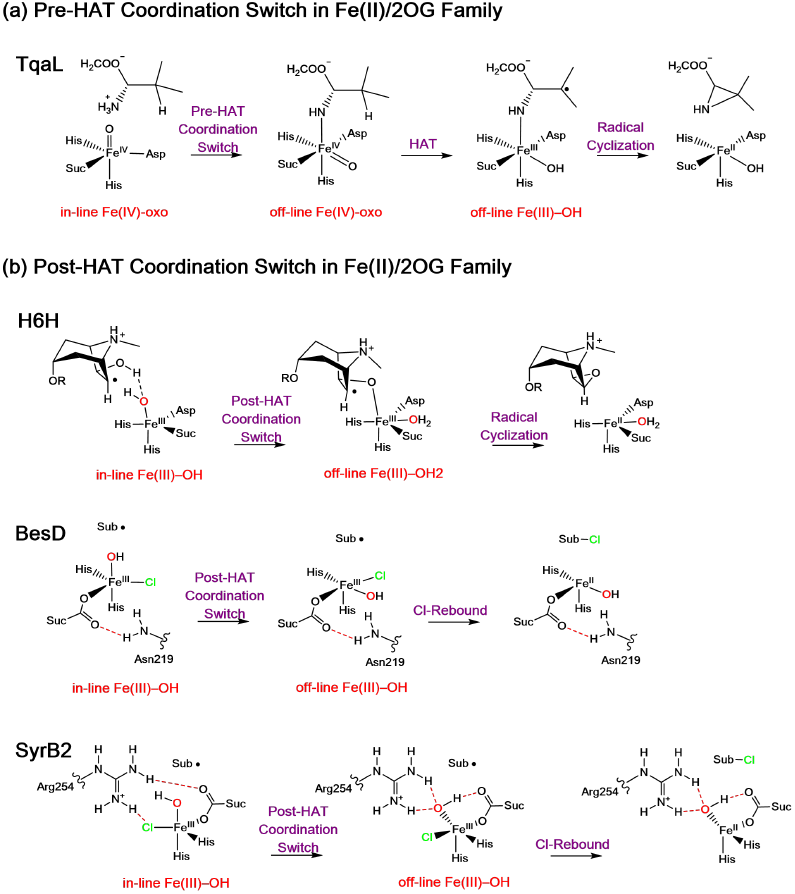
(a) Pre-HAT and (b) post-HAT coordination switch in Fe(II)/2OG dependent enzymes.

Steric effects further influence the selectivity and activity of the reaction. Our QM/MM calculations revealed, for example, that the replacement in H6H of the curved L290 by the flat F290 can reduce the steric interactions between the substrate and F290 for the OH-rebound step. This is mainly because OH-rebound mostly involves the migration of the substrate radical toward the Fe(III)–OH moiety, while the presence of the curved L290 impedes such movement. Consequently, this substitution enhances hydroxylation relative to epoxidation. In additional enzyme engineering efforts, we found that the D219E and T224A substitutions significantly enhance the efficiency of 6,7-exo-epoxidation. This rate enhancement is attributed to the steric effects introduced by these substitutions, which alter substrate positioning. Collectively, these results strongly demonstrate that steric effects arising from second-sphere amino acid residues are critically important for modulating both the selectivity and reactivity of H6H catalysis.

In summary, while the dynamics of iron coordination are primarily responsible for the epoxidation reactivity, the competing hydroxylation pathway is concurrently suppressed by precise substrate positioning and protein-derived steric effects. These insights not only advance our understanding of selectivity control in H6H but also provide a framework for designing enzymes to optimize specific transformations, such as C–O bond formation. More broadly, this work underscores the key roles of metal coordination dynamics and steric effects for modulating reactivity and selectivity in enzymatic catalysis.

## ASSOCIATED CONTENT

Supporting Information

Tables S1-S3, Figures S1-S23, materials and methods, and cartesian coordinates of QM region from QM/MM calculations (PDF). This material is available free of charge via the Internet at http://pubs.acs.org.

## AUTHOR INFORMATION

### Author Contributions

^#^J.Z., L.W. and S.W. contributed equally.

### Notes

The authors declare no competing financial interest.

## ACKNOWLEDGMENT

We thank Prof. Hung-wen Liu and Dr. Mark W. Ruszczycky for their valuable suggestions on our work. This work has been supported by Scientific Research Innovation Capability Support Project for Young Faculty (ZYGXONJSKYCXNLZCXM-B6), Jiangsu Basic Research Center for Synthetic Biology (BK20233003 to J.Z.), National Natural Science Foundation of China (22121001), Fundamental Research Funds for the Central Universities (20720240124), and the National Key Research, Development Program of China (No. 2023YFA0914100).

## Table of Contents

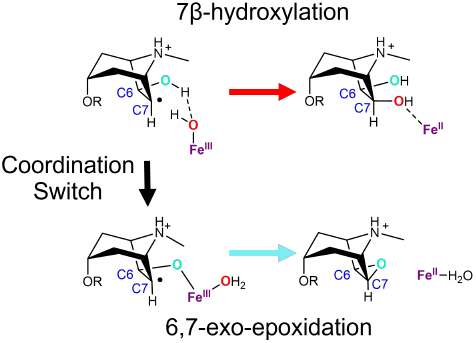

