## Supporting information for "Metal Coordination Dynamics Governs Selective Epoxidation in Hyoscyamine 6β-Hydroxylase: Integrated Experimental and Computational Insights"

#### Metal Coordination Dynamics is a Key Determent for the Selective Epoxidation in Hyoscyamine 6β-Hydroxylase

### Materials and Methods

#### Section S1. Crystallization and Structural Determination

##### Protein expression and purification for crystallization

The plasmid was kindly provided by Prof. Youli Xiao The proteins *Hn*H6H (from Hyoscyamus niger) was expressed in *E. coli* BL21(DE3) in LB medium containing 50 μg $mL^{-1}$ kanamycin, with C-teriminal TEV site and 8×His. Protein expression was induced by 0.4 mM IPTG when $OD_{600}$ reached 1.0, followed by incubation at 16 °C for 18 h. Cells were harvested by centrifugation at 8,000 rpm for 3 min at 4 °C, resuspended in pre-cooled buffer A containing 25 mM Tris (pH 8.0), 500 mM NaCl, 5 mM β-mercaptoethanol, and 1 mM phenylmethylsulfonyl fluoride (PMSF), and lysed by French press with a high-pressure homogenizer (500–600 bar). Cell debris was removed by centrifugation at 18,000 rpm for 30 min at 4 °C. The supernatant was loaded onto a pre-equilibrated Ni-NTA affinity column (GE Healthcare, USA), washed with 100 mL buffer A, and then washed with a buffer containing 25 mM Tris (pH 8.0), 500 mM NaCl, 30 mM imidazole, and 5 mM β-mercaptoethanol. The His-tagged protein was finally eluted with a buffer B containing 25 mM Tris (pH 8.0), 500 mM NaCl, 300 mM imidazole, and 5 mM β-mercaptoethanol. The purified protein was concentrated with an Amicon Ultra-30K (Millipore), and the concentrated protein was then loaded to a pre-balanced Hi Load Superdex G75 column (GE Healthcare) in a buffer containing 25 mM Tris (pH 8.0), 150 mM NaCl, and 1 mM dithiothreitol (DTT). The fractions were then collected and concentrated for crystallization and activity assay.

##### Protein crystallization

Crystals were obtained using the sitting drop vapour diffusion method at 20 °C. In general, 1 μL protein sample (10 mg $mL^{-1}$) was mixed with 1 μL reservoir solution and equilibrated against 50 μL reservoir solution. The native *Hn*H6H crystals were grown in 0.1 M HEPES (pH 7.0), 1 M ammonium sulfate and 1 M potassium chloride, and square-like crystals appeared after one week at 20 °C. For the *Hn*H6H-substrate complex crystals, purified protein (10 mg $mL^{-1}$) was firstly diluted by buffer containing 25 mM Tris (pH 8.0), 150 mM NaCl and 1 mM DTT and mixed with 5 mM 2OG and 2 mM substrate on ice and set screening immediately. The complex crystals of *Hn*H6H•$Fe^{II}$•2OG• hyoscyamine was grown in 0.1 M Sodium acetate, 25% pEG4000 and 8% isopropanol. The crystals of *Hn*H6H•$Fe^{II}$•2OG•6β-hydroxyhyoscyamine were grown in 0.1 M MES pH6.5, 0.2 M

Magnesium chloride, 25%pEG4000. All crystals were flash-frozen in liquid nitrogen using the reservoir solution containing 20% (v/v) glycerol as cryo-protectant.

**Data collection and structure determination**

All X-ray diffraction data were collected at the Shanghai Synchrotron Radiation Facility (SSRF). For apo *Hn*H6H, the complexes *Hn*H6H•$Fe^{II}$•2OG•hyoscyamine, and *Hn*H6H•$Fe^{II}$•2OG•6β-hydroxyhyoscyamine data were collected at the wavelength of ~ 0.9792 Å in beamline BL18U1 and 0.97853 Å in beamline 19U1, respectively. Data reduction and integration were achieved with HKL3000[1] and the XDS[2] software package. The statistics for data collection are listed in Table S1. The phase of apo *Hn*H6H was solved by molecular replacement in the Phenix-Phaser program using 6TTO as searching model.[3] All other complex structures were determined by molecular replacement using Phaser with apo *Hn*H6H crystal structure (PDB ID: 9V7R) as the search model. Iterative cycles of model building and refinement were performed in Coot[4] and Phenix,[3] respectively. Refinement statistics for each final model are recorded in Table S1. Structure figures were drawn using PyMol 2.4.1 (Schrodinger, LLC).[5] The structures were validated by MOLPROBITY[6]. For the flexibility of N-/C-terminal, we could not build the N-terminal 1-29 residues and C-terminal 344-361 residues in almost all *Hn*H6H structures. We checked the real space correlation coefficient (RSCC) of all ligands by validation server of PDB database, and all ligands were well fit the density, RSCC>0.9.[7]

#### Section S2. Computational Details

##### System setup

The structure of wild-type H6H complexed with cofactor succinate and substrate 6β-hydroxyhyoscyamine was prepared based on latest crystal structure of *Hn*H6H•$Fe^{II}$•2OG•6β-hydroxyhyoscyamine (PDB ID: 9V6V), while for L290F, T224A and D219E, mutations were introduced manually. The protonation state of titratable residues was assigned on the basis of pKa values from PROPKA program[8] along with the local hydrogen-bonded networks. Histidine 43,44,45,134,137,217,274 were protonated at δ position, while histidine 66 was protonated at ε position. All glutamic acid and aspartic acid residues were deprotonated. The Amber ff14SB force field[9] was employed for the normal protein residues. For wild-type H6H, the nonheme iron center was model as both in-line and off-line Fe(IV)-oxo, while for L290F, T224A and D219E, the the nonheme iron center was model as in-line Fe(IV)-oxo. The parameters of the Fe(IV)-oxo active site were obtained from the "MCPB.py" tool[10, 11] of AmberTools18.[12] The general AMBER force field (gaff)[13] was used to generate the parameters for the substrate, with the partial atomic charges obtained from the RESP method[14] at the B3LYP/def2SVP level of theory.[15-18] The parmchk2 utility from AmberTools18 was used to generate the missing parameters of the substrate. Sodium irons were added to the protein surface to neutralize the total charge of the system. The resulting system was solvated in a rectangular box of TIP3P waters extending up of a minimum distance of 15 Å from the protein surface.

##### Classical MD simulations

All MD simulations were performed with the GPU version of Amber 18 package.[12] At first the system was fully minimized using combined steepest descent and conjugate gradient methods. Then the system was gently heated from 0 to 300 K under the NVT ensemble for 50 ps with a weak restraint of 15 kcal/mol/Å applied on protein. To achieve a uniform density after the heating dynamics, 1 ns of density equilibration was performed under the NPT ensemble, where the target temperature of 300 K was maintained by using the Langevin thermostat with a collision frequency of 2 ps, while the target pressure of 1.0 atm was controlled by using the Berendsen barostat with a pressure relaxation time of 1 ps. Afterwards, we removed all restraints on the protein and further equilibrated the system for 4 ns under the NPT ensemble. Finally, a productive MD simulation under the NPT ensemble was conducted for 100 ns. The periodic boundary conditions were used for all MD simulations, while the time-step is 2 fs. The covalent bonds containing hydrogen atoms were constrained using SHAKE[19] to enable an integration step of 2 fs. Nonbonded interactions were treated with Particle Mesh Ewald (PME)[20] with the cutoff set at 10 Å. As shown in Figure S2, the MD simulations are well converged in 100 ns.

**Simulated annealing**

To reasonably simulate the plausible conformations of the T224A, we performed simulated annealing (SA) on T224A. The SA protocol was carried out as follows: heating from 0 K to 400 K over 400 ps, maintaining at 400 K for 400 ps, and then cooling from 400 to 0 K over 800ps. To optimize sampling efficiency while preserving the protein's secondary structure, we selected 400 K for the sampling process, applying a weak restraint of 25 kcal/mol/Å to the entire protein complex during the heating phase. Additionally, the cooling rate was reduced to ensure a more gradual temperature drop, minimizing the risk of instabilities from quenching. This completes one full SA cycle, from which only the final snapshot of the cooling phase was retained. A total of 100 SA cycles were performed, all under the NVT ensemble, where the target temperature was controlled using the Langevin thermostat with a collision frequency of 2 $ps^{-1}$. The covalent bonds containing hydrogen atoms were constrained using SHAKE to enable an integration step of 2 fs. Nonbonded interactions were treated with Particle Mesh Ewald (PME) with the cutoff set at 10 A. All simulations were conducted under periodic boundary conditions (PBC). After SA, snaps were further minimized using combined steepest descent and conjugate gradient methods.

**QM/MM methodology**

For wild-type H6H, we select the representative snapshots based on the time evolution of the root mean square deviation (RMSD) and the analysis of the distance between target H and Fe(IV)-oxo species and the corresponding hydrogen-bond network during the MD trajectory (Figure 2a, main text). For the L290F mutant, unfortunately, the substrate could not stably bind during the MD simulations. Therefore, we selected reasonable snapshots from the early stage of the MD trajectory (within 30 ns). For the T224A, we selected snapshots from 100 SA cycles where the Fe(IV)-oxo species was positioned at a favorable distance from the target H of substrate for HAT to occur. All QM/MM calculations were performed with ChemShell software,[21, 22] combining Turbomole[23] for the QM region and DL_POLY[24] for the MM region with the Amber force field. The electronic embedding[25] scheme was used to account for the polarizing effect of the enzyme environment on the QM region. Hydrogen link atoms with the charge-shift model were applied to treat the QM/MM boundary. The QM/MM system contains the whole protein and solvation waters within 8 Å of protein. During QM/MM geometry optimizations, the QM region was treated with hybrid UB3LYP density functional.[15-17] For geometry optimization and frequency calculations, the double-ζ basis set def2-SVP[18] (labeled as B1) was used. Transition states were located with relaxed potential energy surface scans followed by full TS optimizations using the dimer optimizer implemented in the DL-FIND code. The energies of all species were further corrected with a larger basis set def2-TZVP[18] (labeled as B2). Dispersion corrections computed with Grimme's D3BJ method[26-28] were included in QM regions in all QM/MM calculations. The reported final energies in Figure 3, Figure 5, Figure S4 and Figure S8 include the electronic energies at the B3LYP/B2 level and zero-point energies (ZPE) at the B3LYP/B1 level, as well as dispersion corrections. Atoms included in the QM regions are further demonstrated in Figure S3.

**Section S3. Mutagenesis Experiments**

**General**

The chemicals were purchased from InnoChem Science & Technology Co., Ltd. (Beijing, China) and MedChemExpress (Shanghai, China) without further purification. The gene encoding a H6H protein (Genbank no: M62719) was chemically synthesized by Generay Biotech Co., Ltd. (Shanghai, China) with a codon-optimization for *E. coli* expression. Primer synthesis and DNA sequencing were conducted at Tsingke Biotech Co., Ltd. (Beijing, China).

The HPLC analyses were performed on an Agilent 1260 Infinity II HPLC system (Agilent Technologies, Germany) equipped with a SUPERIOREX ODS column (250 mm × 4.6 mm I.D., 5 μm, Osaka Soda, Japan) at a flow rate of 1 mL/min and maintained at a temperature of 35 °C. The mobile phase consisted of a gradient elution system comprising solvents A (0.08% trifluoroacetic acid (TFA) aqueous solution) and B (acetonitrile containing 0.08% TFA). The gradient program was as follows: 0–22 min, linear change from A:B (90:10, *v*/*v*) to A:B (70:30, *v*/*v*); 22–23 min, linear change to A:B (0:100, *v*/*v*); 23–28 min, isocratic elution with A:B (0:100, *v*/*v*); 28–28.1 min, linear change to A:B (90:10, *v*/*v*); 28.1–35 min, isocratic elution with A:B (90:10, *v*/*v*). The HPLC-ESI-HRMS analysis was conducted using a Shimadzu LC20A HPLC System (Shimadzu, Kyoto, Japan) coupled with an LCMS-9030 qToF mass spectrometer (Shimadzu, Kyoto, Japan) equipped with an electrospray ionization (ESI) source. Data acquisition, analysis and reporting were performed using Shimadzu LabSolutions software and CBM-20 A system controller.

**Gene cloning**

The codon-optimized *H6H* gene was synthesized and inserted into pET-24b (+) vector between the *Nde* I and *BamH* I sites to generate an expression construct for efficient production of CHis6-tagged H6H. The resulting plasmid was transformed into *E. coli* BL21 (DE3) for protein expression. The mutated *H6H* DNA sequence was generated using the Fast Mutagenesis System (TransGen, Beijing, China), following the manufacturer's protocol with the wild-type *H6H* expression plasmid as a template. The resulting construct was amplified in *E. coli* DH5a, confirmed by sequencing, and utilized for the transformation of BL21 (DE3) competent cells.

**Protein expression and purification**

The cells harboring the recombinant plasmid were cultured in LB medium supplemented with kanamycin (30 mg/L) at 16 °C under constant shaking at 260 r.p.m. until the $OD_{600}$ reached 0.6. Subsequently, protein overexpression was induced by adding isopropyl β-D-1-thiogalactopyranoside (IPTG) to a final concentration of 0.1 mM. The resultant culture was further incubated at 16 °C for 18 h to facilitate H6H protein expression.

The cells were pelleted by centrifugation and subsequently disruptured using a sonifier. Debris and membranes were removed by centrifugation at 10,000 g for 45 min. The resulting supernatant was then subjected to purification using a Ni Sepharose™ 6 fast flow resin column (GE Healthcare, Piscataway, USA) following the manufacturer's instructions. A linear gradient of 20–500 mM imidazole (in 50 mM Tris-HCl, 300 mM NaCl, and 10% (*v*/*v*) glycerol, pH 7.4) was employed for separation. After SDS–PAGE analysis, fractions containing the desired protein were collected and concentrated by ultrafiltration with 10 kDa ultrafiltration tubes (Millipore, Billerica, USA). To facilitate buffer exchange, the protein fraction was subjected to a PD-10 column (GE Healthcare, Piscataway, USA) and eluted with 50 mM Tris-HCl, 10% (*v*/*v*) glycerol at pH 7.4. The resulting protein solution was concentrated and stored at −80 °C prior to being used in enzymatic assays.

#### In vitro enzyme activity assays

Hyoscyamine (1 mM) was incubated with H6H (24 μM), 2OG (8 mM), $FeSO_4$ (2 mM), sodium ascorbate (8 mM) in Tris-HCl buffer (50 mM, pH 7.4) at 30 °C for 16 h (100 μL total volume). Three parallel assays were routinely carried out for each reaction. The reaction was terminated by adding of two reaction volumes of acetonitrile and the resulting mixture was centrifuged at 12,000 g for 30 min to remove precipitated proteins. The supernatant was subjected to analysis using HPLC. The enzymatic products and unconsumed substrate were monitored at 220 nm, and the conversion rates were determined based on standard curves for the substrate and products, respectively.

#### Enzyme kinetics

To determine the kinetic parameters, increasing concentrations (10–750 μM) of 6β-hydroxyhyoscyamine were incubated with H6H (24 μM), 2OG (8 mM), $FeSO_4$ (2 mM), and sodium ascorbate (8 mM) in Tris-HCl buffer (50 mM, pH 7.4) at 30 °C for 1 h. The enzymatic products were quantified by HPLC-ESI-HRMS analysis. Kinetic parameters were calculated by nonlinear regression analysis using GraphPad Prism 10 software.

**Optimized H6H sequence**

>*H6H_opt*

ATGGCAACCTTTGTGAGTAATTGGAGTACCAAATCAGTGAGCGAATCTTTTATTGCACCGTT
GCAGAAACGCGCAGAAAAAGATGTGCCGGTGGGTAATGATGTGCCGATTATTGATCTCCAGC
AGCATCATCATCTGCTGGTTCAGCAGATTACCAAAGCATGTCAGGATTTTGGCCTGTTTCAG
GTTATTAATCATGGCTTTCCGGAAGAACTGATGCTGGAAACTATGGAAGTGTGTAAAGAATT
TTTTGCACTGCCGGCAGAAGAAAAAGAAAAATTTAAACCGAAAGGCGAAGCAGCAAAATTTG
AACTGCCGCTGGAACAGAAAGCAAAACTGTATGTTGAAGGCGAACAGCTGTCTAATGAAGAA
TTTCTGTATTGGAAAGATACCCTGGCACATGGTTGTCATCCGCTGGATCAGGATCTGGTTAA
TAGTTGGCCGGAAAAACCGGCAAAATATCGCGAAGTGGTGGCAAAATATAGCGTGGAAGTTC
GTAAACTGACCATGAGGATGCTGGATTATATTTGTGAAGGTCTGGGTCTGAAACTGGGCTAT
TTTGATAATGAACTGTCACAGATTCAGATGATGCTGACCAATTATTATCCGCCGTGTCCCGA
TCCGTCTAGTACCCTGGGCTCAGGCGGCCATTATGATGGTAATCTGATTACCCTGCTACAGCA
GGATCTGCCGGGCTTGCAGCAGCTGATTGTTAAAGATGCAACCTGGATTGCAGTTCAGCCGA
TTCCGACCGCATTTGTTGTTAATCTGGGTCTGACCCTGAAAGTTATTACCAATGAAAAATTT
GAAGGTAGCATTCATCGCGTGGTGACCGATCCGACCCGTGATCGCGTTTCTATTGCAACCCTG
ATTGGCCCGGATTATTCTTGTACCATTGAACCGGCAAAAGAACTGCTGAATCAGGATAATCC
GCCGCTGTATAAACCGTATAGCTATAGCGAATTTGCAGATATTTATCTGAGCGATAAAAGCG
ATTATGATAGCGGCGTTAAACCGTATAAAATTAATGTGTAA

**Isolation and characterization of 6β-chloro-hyoscyamine**

The 6β-chloro-hyoscyamine was isolated by carrying out a large-scale enzymatic reaction with the purified D219A protein and hyoscyamine under the conditions described above. The isolation was conducted using HPLC equipped with a CAPCELL PAK C18 MGIII column (250 mm × 10 mm I.D., 5 μm, Osaka Soda, Japan) maintained at a temperature of 35 °C. The mobile phase consisted of a gradient elution system with solvents A (0.08% TFA aqueous solution) and B (acetonitrile containing 0.08% TFA), flowing at a rate of 3 mL/min with the following gradient: 0–18 min (10% B–38% B), 18–19 min (38% B–100% B), 19–21 min (100% B–10% B), 21–30 min (10% B–10% B). The fractions containing the product were lyophilized to dryness and analyzed by HR-ESI-MS and NMR.

The molecular formula of isolated product was assigned as $C_{17}H_{22}ClNO_3$ by HR-ESI-MS (*m/z* 324.1362 $[M+H]^+$, calcd. 324.1361) together with the diagnostic ratio of isotopic ions 3:1 at *m/z* 324 and 326, which suggested that it may be a chlorinated product of hyoscyamine. $^1H$ NMR analysis in $CD_3OD$ revealed all the main proton signals for the structure. However, most carbon-13 signals in tropane moiety could not be detected in $^{13}C$ NMR spectrum with the same solvent. Finally, NMR spectra with complete carbon signals were acquired in $CD_3OD$ containing 0.5 M DCl. The C6β-chlorination was confirmed via a combination of 1D and 2D NMR analysis (Figure S17–S23).

HR-ESI-MS (positive) *m/z*: 324.1362 $[M+H]^+$ (calcd. 324.1361 for $C_{17}H_{22}ClNO_3$); $^1H$ NMR ($CD_3OD$ with 0.5 M DCl, 500 MHz) $\delta$ 7.31–7.41 (5H, m, H-4'–9'), 5.04 (1H, m, H-3), 4.81 (1H, dd, *J*= 8.5, 4.0 Hz, H-6), 4.16 (1H, m, overlaped, H-5), 4.15 (1H, dd, *J*= 10.5, 8.5 Hz, H-3'a), 4.07 (1H, m, H-1), 3.90 (1H, dd, *J*= 8.5, 6.0 Hz, H-2'), 3.83 (1H, dd, *J*= 10.5, 6.0 Hz, H-3'b), 3.07 (3H, s, NMe), 2.59 (1H, dt, *J*= 16.5, 4.0 Hz, H-4b), 2.49 (1H, dt, *J*= 16.5, 4.0 Hz, H-2b), 2.43 (1H, m, H-7b), 2.35 (2H, m, H-4a, 7a), 1.90 (1H, d, *J*= 16.5 Hz, H-2a); $^{13}C$ NMR ($CD_3OD$ with 0.5 M DCl, 125 MHz) $\delta$ 172.58 (C-1'), 137.14 (C-4'), 130.13 (×2, C-6', 8'), 129.32 (×2, C-5', 9'), 129.06 (C-7'), 72.56 (C-5), 65.69 (C-3), 64.84 (C-1), 64,32 (C-3'), 56.68 (C-6), 55.46 (C-2'), 44.51 (NMe), 38.40 (C-7), 35.41 (C-4), 34.74 (C-2).

**Table S1.** Data collection and refinement statistics for *Hn*H6H crystal structures

| | *Hn*H6H•$Fe^{II}$ | *Hn*H6H•$Fe^{II}$•2OG•hyoscyamine | *Hn*H6H•$Fe^{II}$•2OG•6β-hydroxyhyoscyamine |
|---|---|---|---|
| PDB code | 9V7R | 9V7M | 9V6V |
| Data collection | | | |
| Space group | $P_1$ | $P2_1$ | $P2_1$ |
| Cell dimensions | | | |
| a, b, c (Å) | 43.13, 43.16, 91.46 | 43.21, 80.07, 47.80 | 43.37, 80.04, 48.23 |
| α, β, γ (∘) | 80.77, 87.93, 83.35 | 90.00, 98.12, 90.00 | 90.00, 97.45, 90.00 |
| Resolution (Å) | 50.00-2.45 (2.55-2.45) | 19.57-1.70 (1.73-1.70) | 19.38- 1.70 (1.73-1.70) |
| Rmerge | 0.076 (0.344) | 0.105 (0.754) | 0.048 (0.211) |
| Rpim | 0.047 (0.222) | 0.048 (0.316) | 0.020 (0.088) |
| CC1/2 [a] | 0.996 (0.851) | 0.996 (0.882) | 0.999 (0.977) |
| No. of unique reflections | 23244 | 35429 (1905) | 35949 (1957) |
| Ι /σ Ι | 13.0 (3.5) | 13.3 (3.1) | 22.7 (7.0) |
| Completeness (%) [a] | 97.7 (97.7) | 99.9 (99.8) | 99.9 (100.0) |
| Redundancy | 3.5 (3.3) | 6.6 (6.6) | 6.4 (6.6) |
| Refinement | | | |
| Resolution (Å) | 19.82-2.45 (2.56-2.45) | 19.57-1.70 (1.72-1.70) | 19.38-1.70 (1.75-1.70) |
| No. reflections | 23237 (1176) | 69568 (3557) | 35925 (1824) |
| Rwork/Rfree | 0.1765 /0.2351 | 0.1667 /0.1989 | 0.1635 /0.1905 |
| No. atoms | | | |
| Protein | 4781 | 2547 | 2567 |
| Ligand | 18 | 39 | 40 |
| Water | 157 | 289 | 303 |
| Wilson B factors | 29.35 | 17.80 | 16.36 |
| Average B factors ($Å^2$) | 34.10 | 21.95 | 21.88 |
| Macromolecules | 34.21 | 21.18 | 20.82 |
| Ligands | 39.54 | 16.54 | 16.47 |
| solvent | 30.57 | 29.41 | 31.52 |
| R.m.s. deviations | | | |
| bond lengths (Å) | 0.006 | 0.007 | 0.007 |
| bond angles (°) | 0.863 | 0.866 | 0.870 |
| Ramachandran Outliers (%) | 0.17 | 0.64 | 0.32 |
| Ramachandran Favored (%) | 95.85 | 97.12 | 98.08 |

[a] Statistics for the highest-resolution shell are shown in parentheses.

**Table S2.** The main primers used in this study

| Primers | Sequences (5' to 3') |
|---|---|
| Hn-D219A-F | GGGCTCAGGCGGCCATTATGCTGGTAATCTGATTAC |
| Hn-D219A-R | AGCATAATGGCCGCCTGAGCCCAGGGTACTAGACGGATC |
| Hn-D219E-F | GGGCTCAGGCGGCCATTATGAAGGTAATCTGATTAC |
| Hn-D219E-R | TTCATAATGGCCGCCTGAGCCCAGGGTACTAGACGGATC |
| Hn-D219G-F | GGGCTCAGGCGGCCATTATGGCGGTAATCTGATTAC |
| Hn-D219G-R | GCCATAATGGCCGCCTGAGCCCAGGGTACTAGACGGATC |
| Hn-D219N-F | GGGCTCAGGCGGCCATTATAATGGTAATCTGATTAC |
| Hn-D219N-R | ATTATAATGGCCGCCTGAGCCCAGGGTACTAGACGGATC |
| Hn-T224A-F | CATTATGATGGTAATCTGATTGCGCTGCTACAGCAGGATC |
| Hn-T224A-R | CGCAATCAGATTACCATCATAATGGCCGCCTGAGCCC |
| Hn-N256A-F | CCGACCGCATTTGTTGTTGCGCTGGGTCTGACCCTG |
| Hn-N256A-R | CGCAACAACAAATGCGGTCGGAATCGGCTGAACTGC |
| Hn-N256D-F | CCGACCGCATTTGTTGTTGATCTGGGTCTGACCCTG |
| Hn-N256D-R | ATCAACAACAAATGCGGTCGGAATCGGCTGAACTGC |
| Hn-N256S-F | CCGACCGCATTTGTTGTTAGCCTGGGTCTGACCCTG |
| Hn-N256S-R | GCTAACAACAAATGCGGTCGGAATCGGCTGAACTGC |

**Table S3.** The steady-state enzyme kinetics values for the wild-type and mutants of H6H catalyzing the epoxidation of 6β-hydroxyhyoscyamine

| H6H | $K_M$ (μM) | $K_{cat}$ ($s^{-1}$) | $K_{cat}/K_M$ ($s^{-1}mM^{-1}$) |
|---|---|---|---|
| WT | 22.97±3.55 | 0.678*$10^{-3}$ | 0.030 |
| T224A | 10.22±1.76 | 3.641*$10^{-3}$ | 0.356 |
| N256A | 6.85±1.49 | 0.006*$10^{-3}$ | 0.001 |

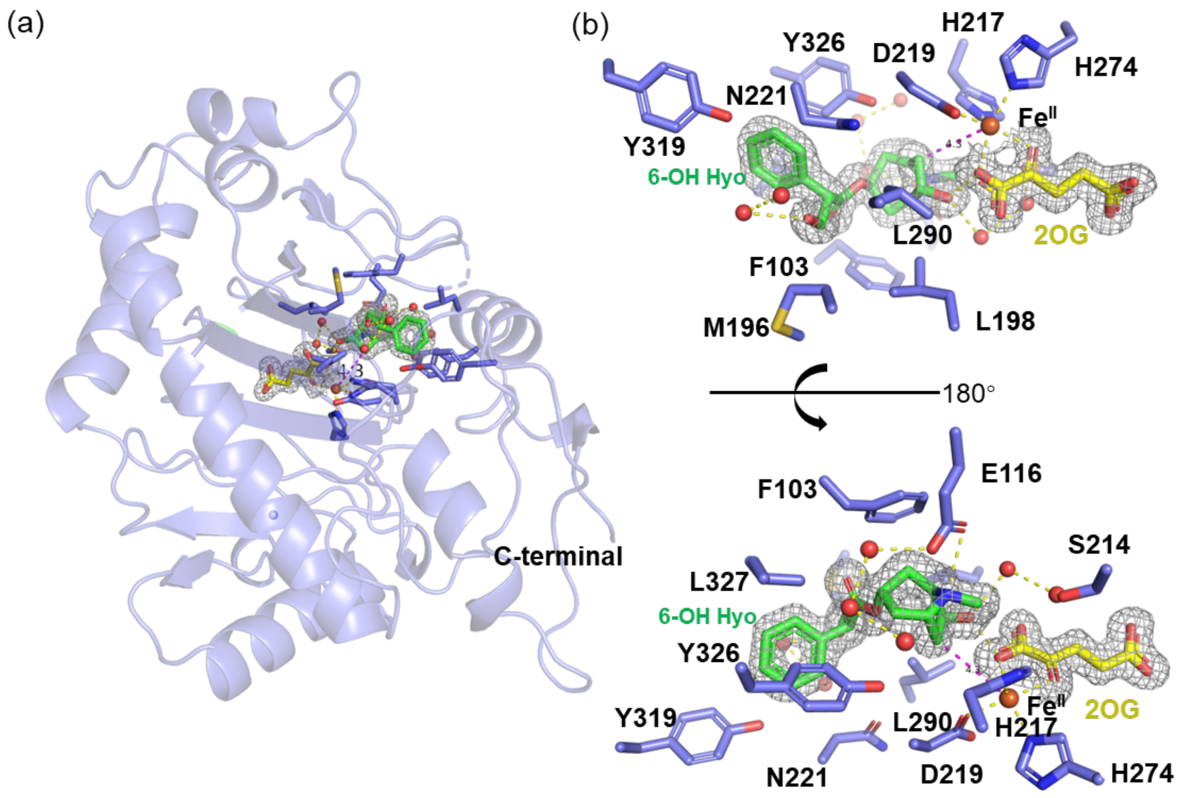

**Figure S1.** The complex structure of *Hn*H6H complex with 6-hydroxyhyoscyamine (6-OH Hyo), 2OG and iron. (a) The overall structure of *Hn*H6H•$Fe^{II}$•2OG•6-OH Hyo, was shown as cartoon in slate. The residues in active site pocket and 2OG (yellow), 6-hydroxyhyoscyamine (green) were shown as sticks. (b) The details of active site pocket, 2OG and 6-OH Hyo were modeled into its omit map (gray), contoured at 3.0 σ. The residues of active pocket could be clearer by above and below two angles. The distance between C7 of 6-hydroxyhyoscyamine and iron center was 4.5 Å, labeled by dash line in magenta. The hydrogen bond was labeled by yellow dash line.

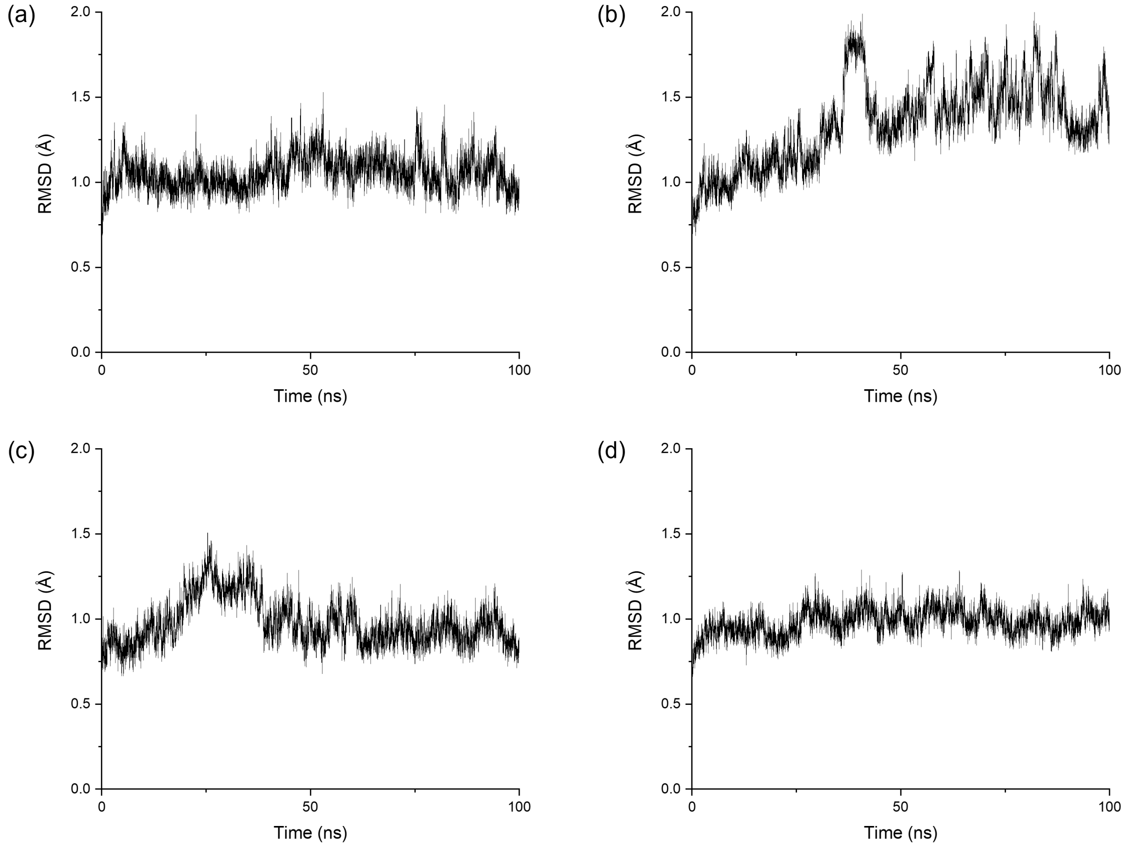

**Figure S2.** Time evolution of the root mean square deviations (RMSD) for the protein backbone and substrate of (a) wtH6H (in-line Fe(IV)-oxo), (b) wtH6H (off-line Fe(IV)-oxo), (c) T224A (in-line Fe(IV)-oxo) and (d) D219E (in-line Fe(IV)-oxo).

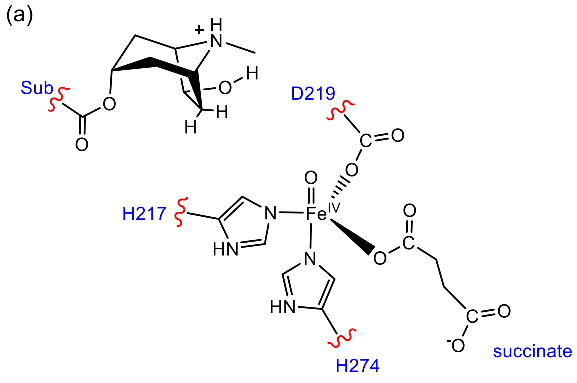

**Figure S3.** QM region for QM/MM calculations of (a) wild-type H6H and T224A, (b) L290F.

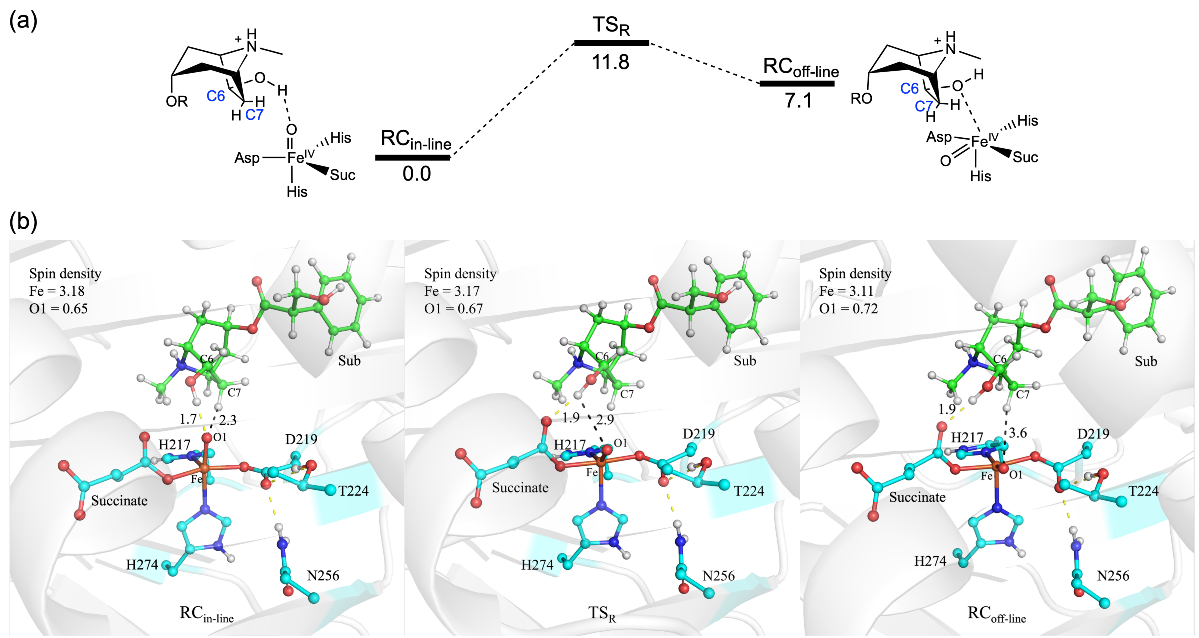

**Figure S4.** (a) QM(UB3LYP/B2)/MM calculated energy profile (in kcal $mol^{-1}$) for isomerization of Fe(IV)-oxo species in H6H. The dispersion corrections and ZPEs are included in the relative energies. (b) QM(UB3LYP/B1)/MM optimized structures of key species involved in the reaction. The key distances are given in Å.

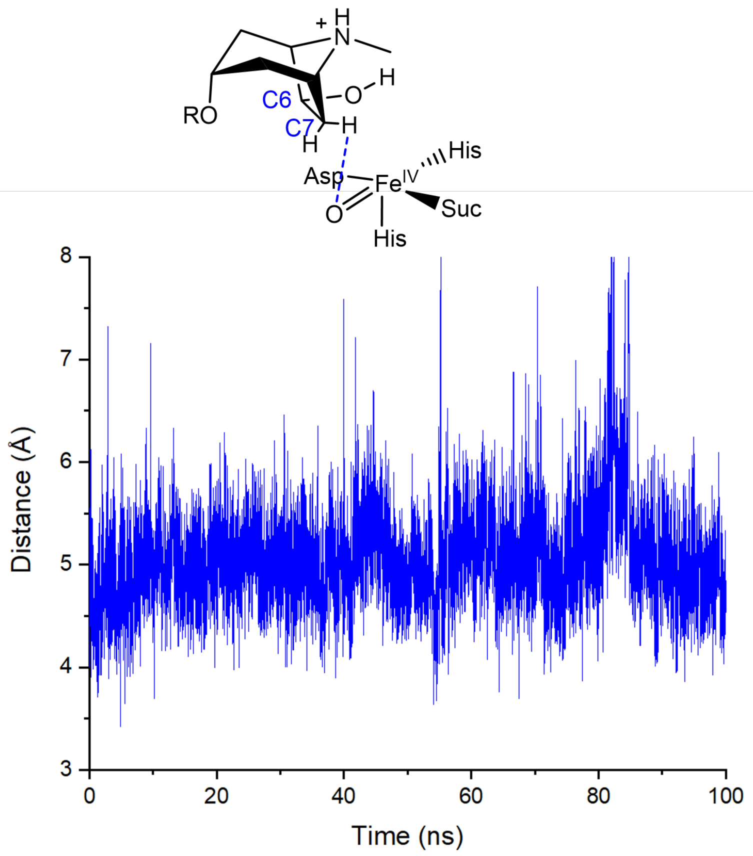

**Figure S5.** Time evolution of distance of ferryl O atom and H(C7) atom in off-line conformation in 100 ns MD simulation.

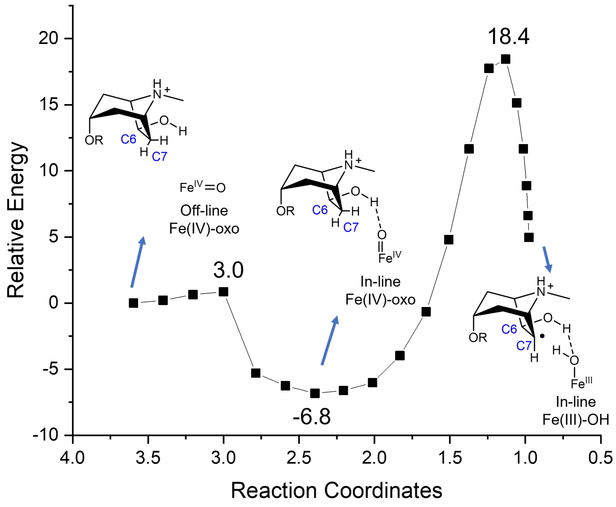

**Figure S6.** QM(UB3LYP/B1)/MM scanned relative energies (in kcal mol$^{-1}$) for HAT mediated by off-line Fe(IV)-oxo.

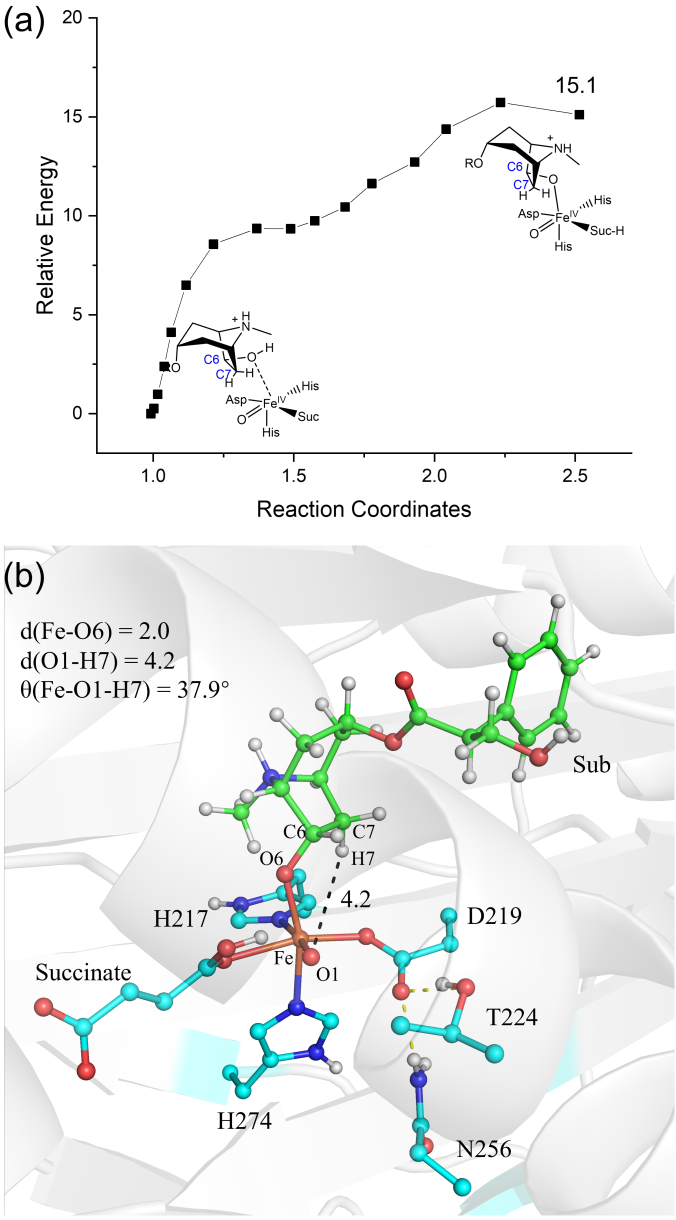

**Figure S7.** (a) QM(UB3LYP/B1)/MM scanned relative energies (in kcal mol$^{-1}$) for substrate coordination in off-line conformation, where the proton from substrate's OH group was shifted to succinate. (b) Structure of coordination geometry obtained by QM(UB3LYP/B1)/MM scan. The key distances are given in Å.

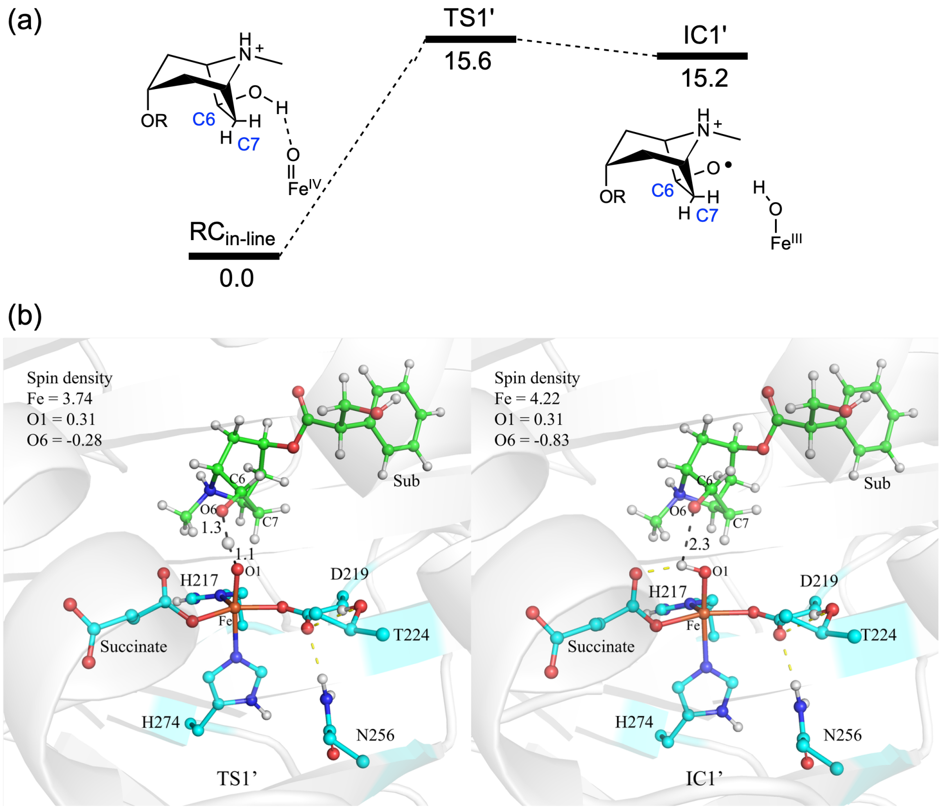

**Figure S8.** (a) QM(UB3LYP/B2)/MM calculated energy profile (in kcal mol$^{-1}$) for HAT from hydroxyl group of 6-OH Hyo. The dispersion corrections and ZPEs are included in the relative energies. (b) QM(UB3LYP/B1)/MM optimized structures of key species involved in the reaction. The key distances are given in Å.

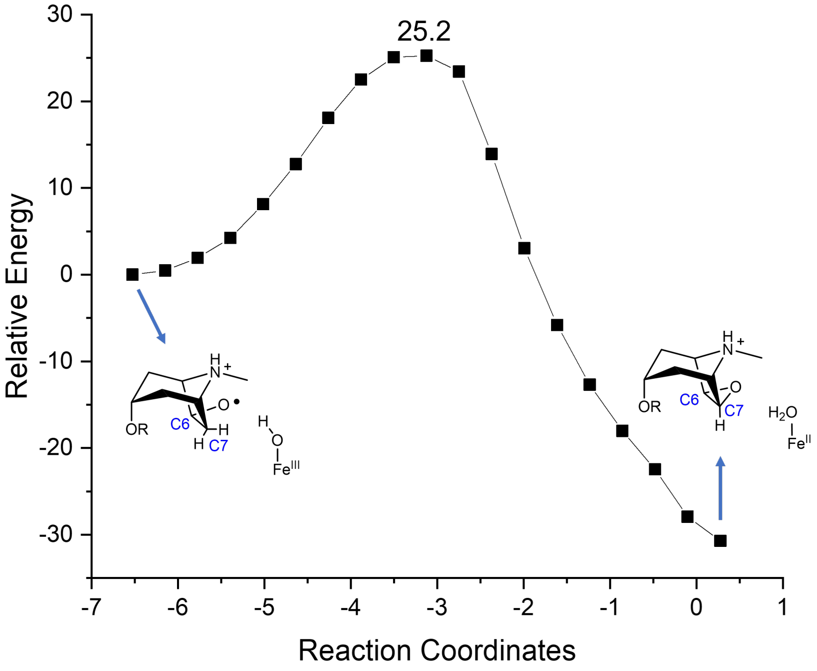

**Figure S9.** QM(UB3LYP/B1)/MM scanned relative energies (in kcal mol$^{-1}$) for C–O coupling from IC1'.

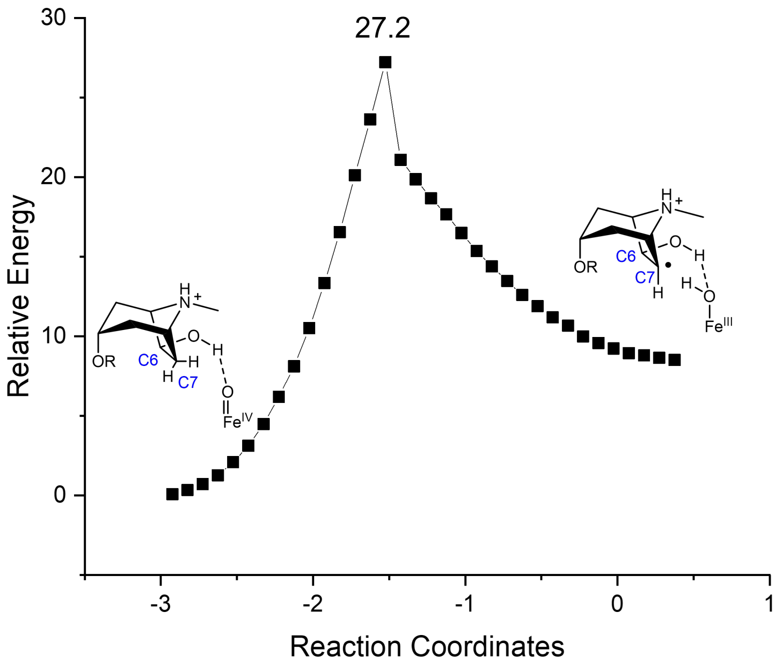

**Figure S10.** QM(UB3LYP/B1)/MM scanned relative energies (in kcal $mol^{-1}$) for $P^2E^2T$ mechanism.

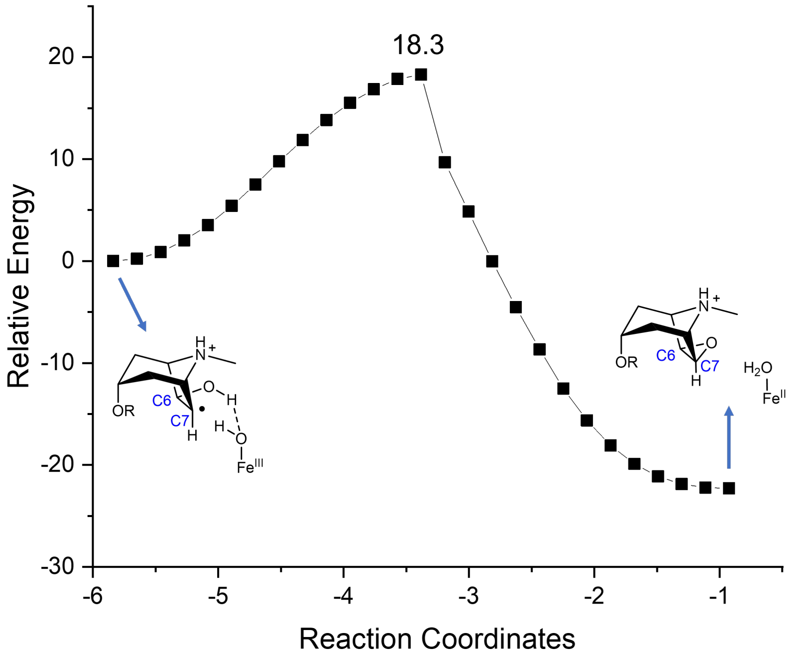

**Figure S11.** QM(UB3LYP/B1)/MM scanned relative energies (in kcal mol$^{-1}$) for direct C7-O6 coupling coupled with HAT from O6 to Fe(III)-OH.

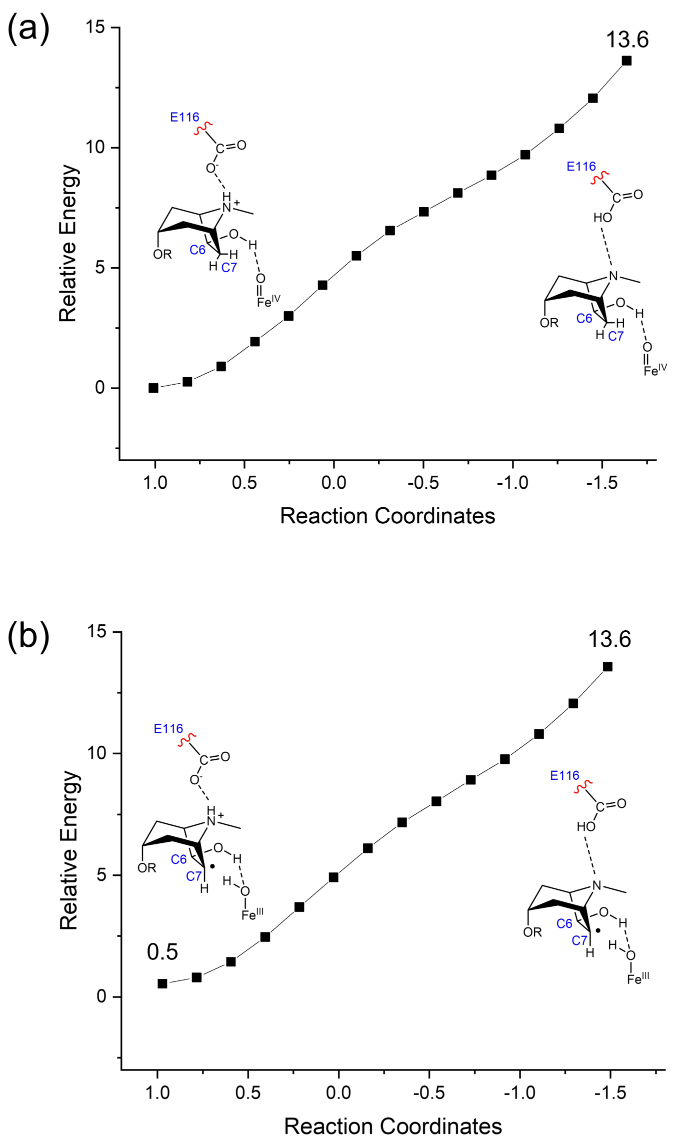

**Figure S12.** QM(UB3LYP/B1)/MM scanned relative energies (in kcal mol$^{-1}$) for proton transfer from substate to E116 in (a) Fe(IV)-oxo and (b) Fe(III)–OH states.

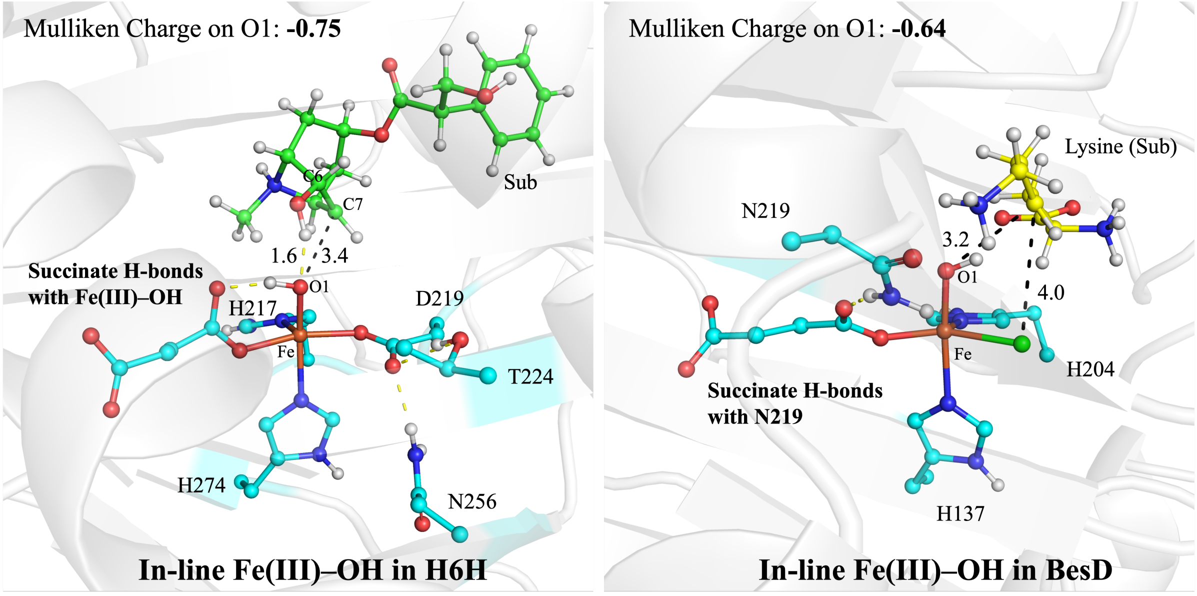

**Figure S13.** Structure and charge distribution comparison of Fe(III)–OH species in H6H and BesD.

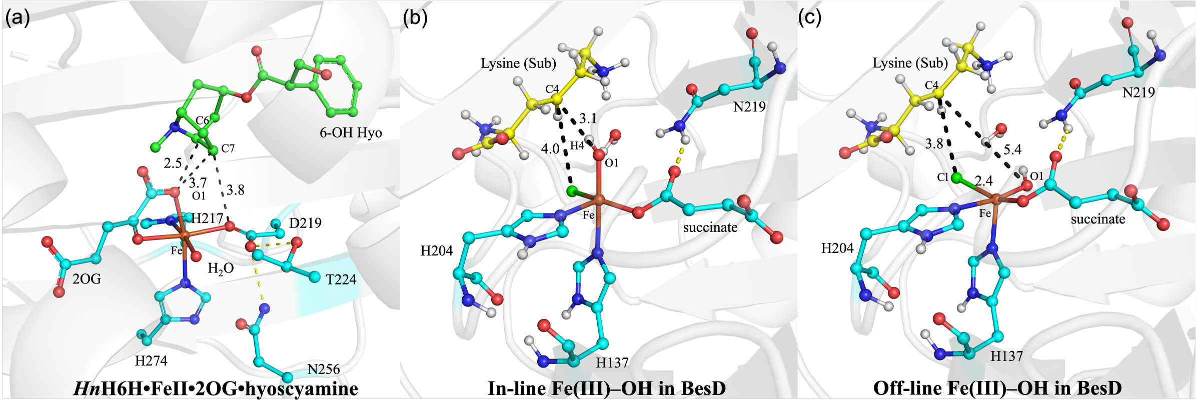

**Figure S14.** (a) Crystal structure of *Hn*H6H•Fe$^{II}$•2OG•hyoscyamine (PDB ID: 8HV0). (b) QM(UB3LYP/B1)/MM optimized structures of in-line Fe(III)–OH species in BesD. (c) QM(UB3LYP/B1)/MM optimized structures of off-line Fe(III)–OH species in BesD. (Reproduced from our previous work, main text reference 71)

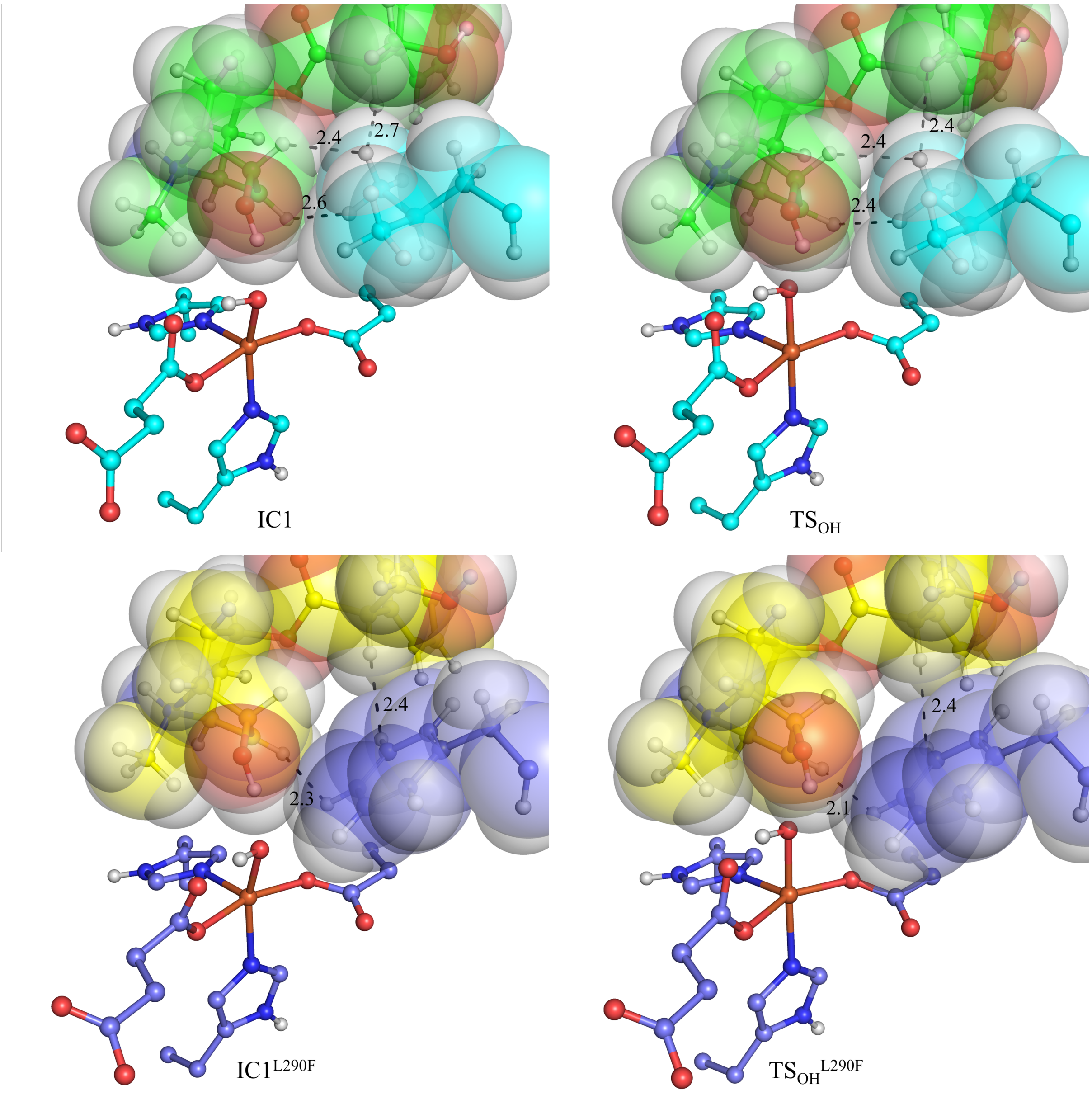

**Figure S15.** QM(UB3LYP/B1)/MM optimized structures of key species involved in OH-rebound. Spheres are shown to emphasis steric effect. The key distances are given in Å. Carbon atoms in wild-type (active site) and wild-type (substrate) are shown in cyan and green, while carbon atoms in L290F (active site) and L290F (substrate) are shown in purple and yellow, respectively.

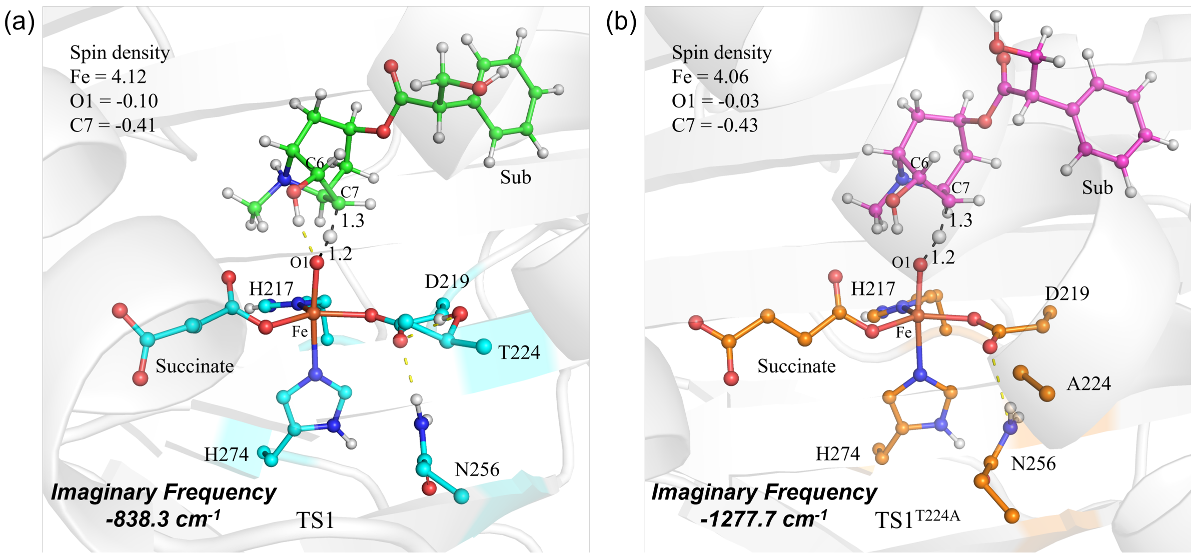

**Figure S16.** QM(UB3LYP/B1)/MM optimized structures of key species involved in HAT catalyzed by (a) wild-type H6H and (b) T224A.

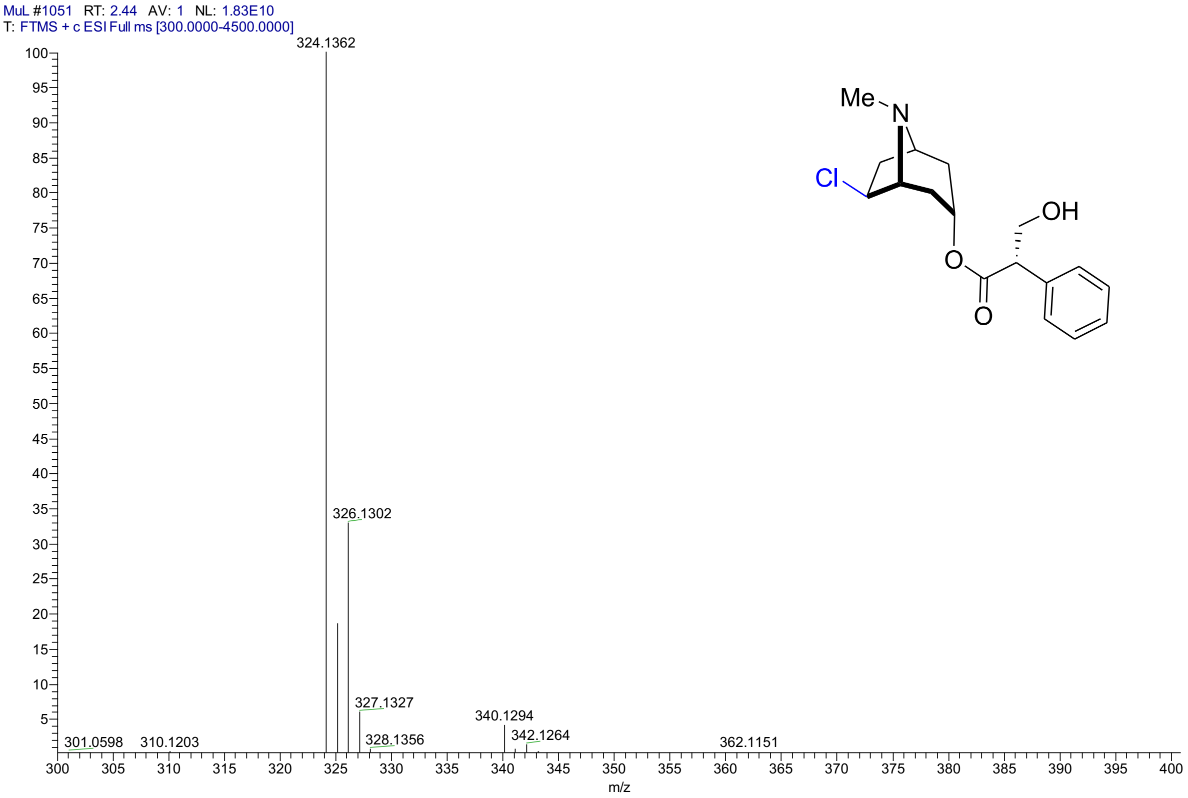

**Figure S17.** HR-ESI-MS spectrum (positive) of 6β-chloro-hyoscyamine

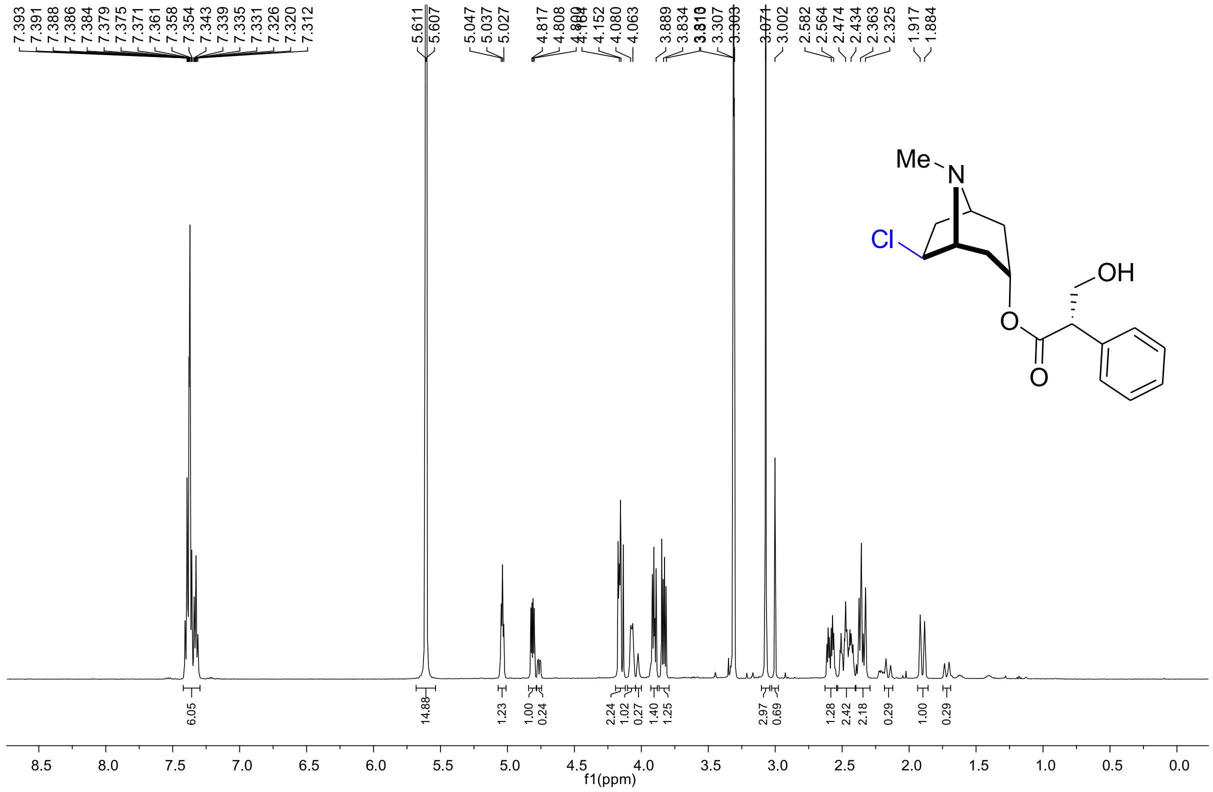

**Figure S18.** [1]H NMR spectrum of 6β-chloro-hyoscyamine (500 MHz in $CD_3OD$ with 0.5 M DCl)

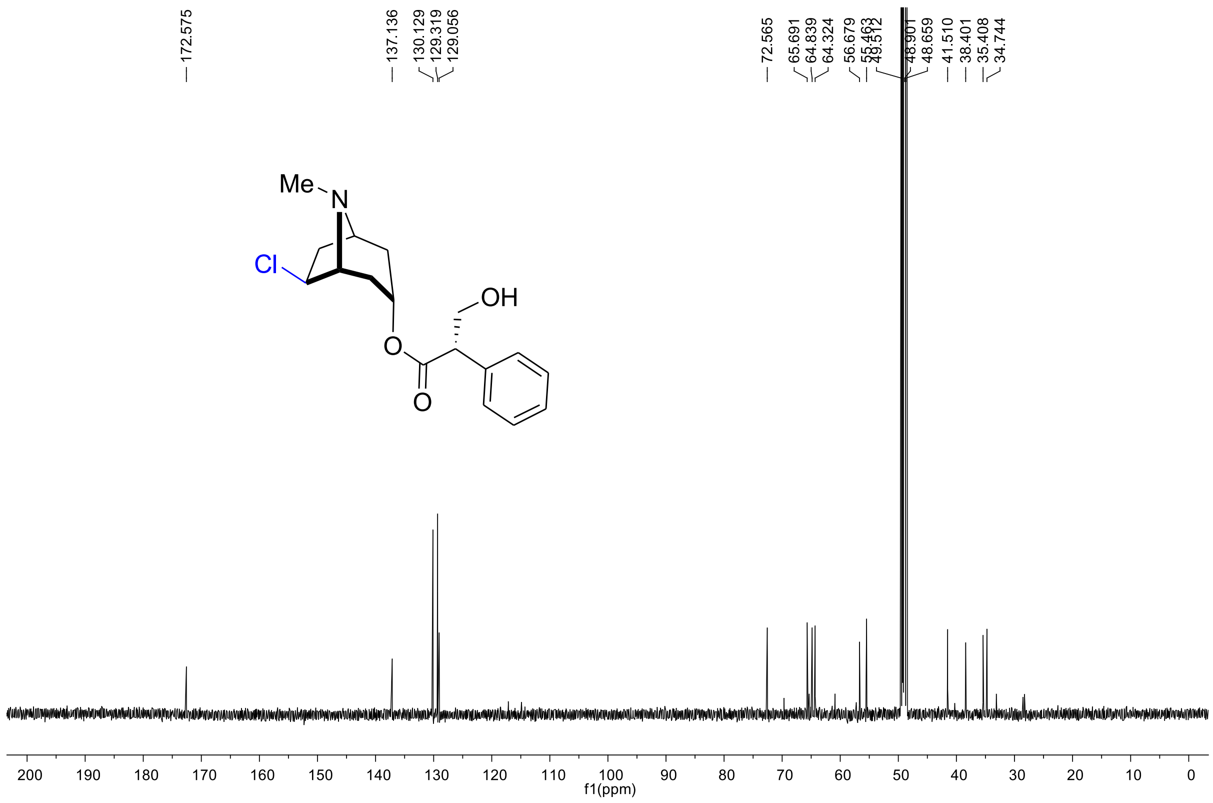

**Figure S19.** $^{13}C$ NMR spectrum of 6β-chloro-hyoscyamine (125 MHz in $CD_3OD$ with 0.5 M DCl)

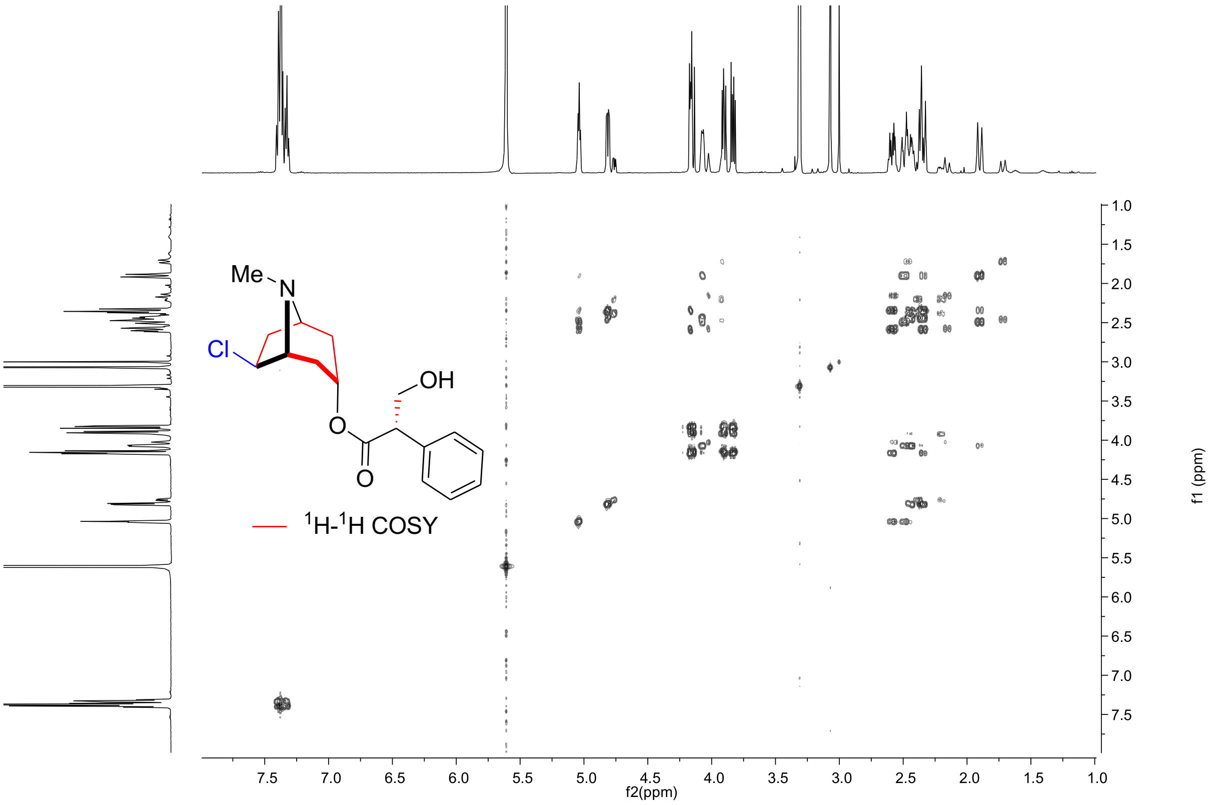

**Figure S20.** $^{1}H$-$^{1}H$ COSY spectrum of 6β-chloro-hyoscyamine (500 MHz in $CD_3OD$ with 0.5 M DCl)

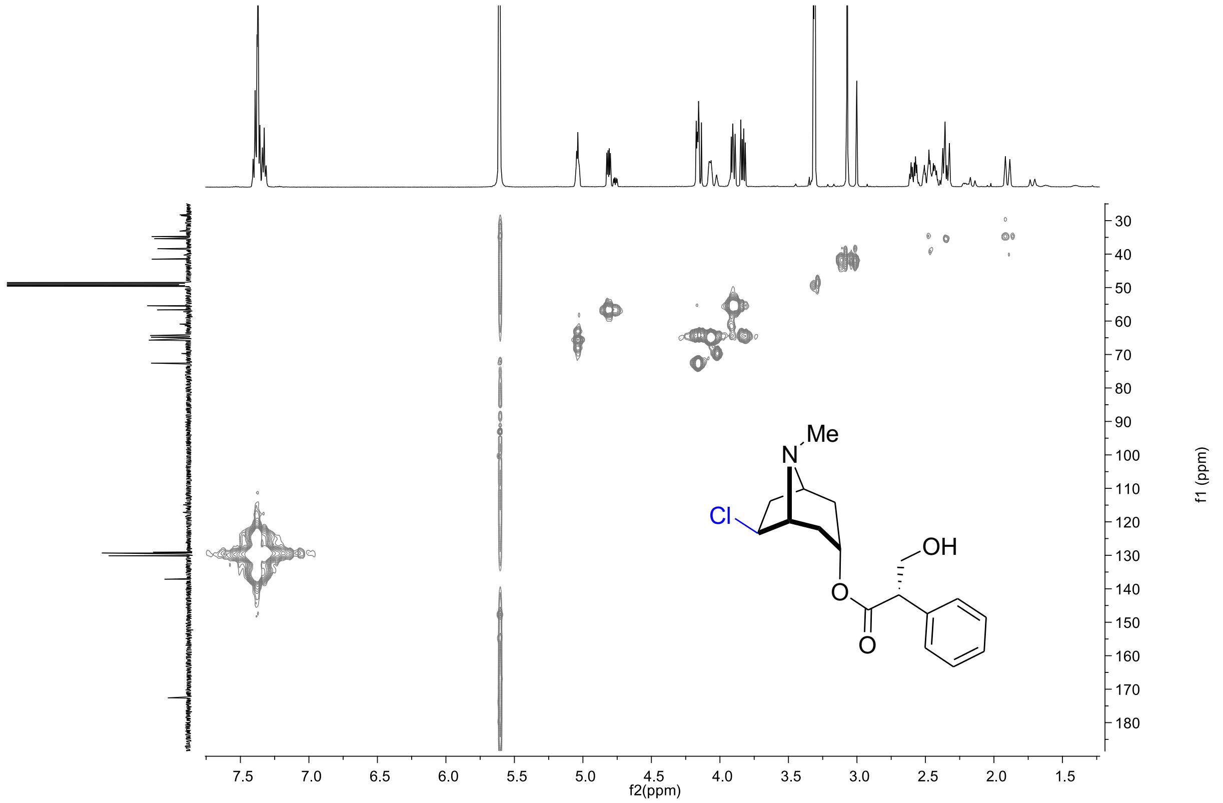

**Figure S21.** HSQC spectrum of 6β-chloro-hyoscyamine (500 MHz in $CD_3OD$ with 0.5 M DCl)

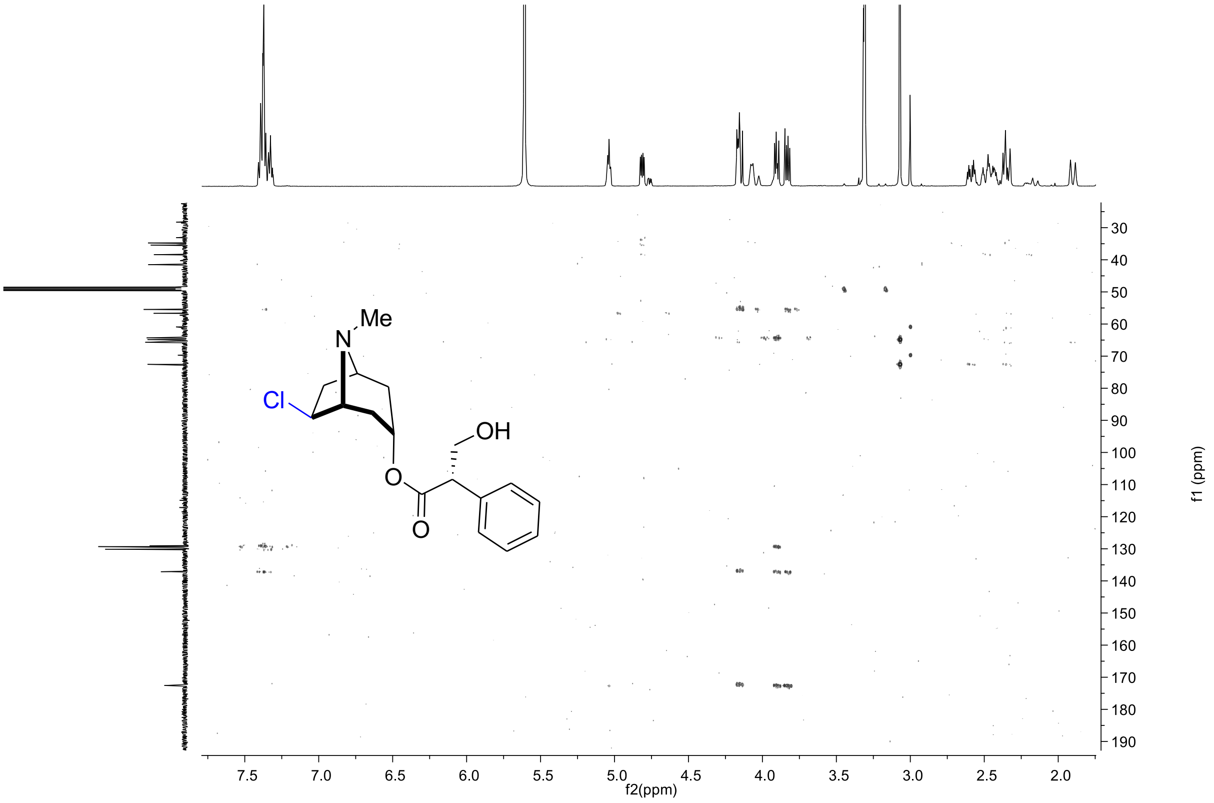

**Figure S22.** HMBC spectrum of 6β-chloro-hyoscyamine (500 MHz in $CD_3OD$ with 0.5 M DCl)

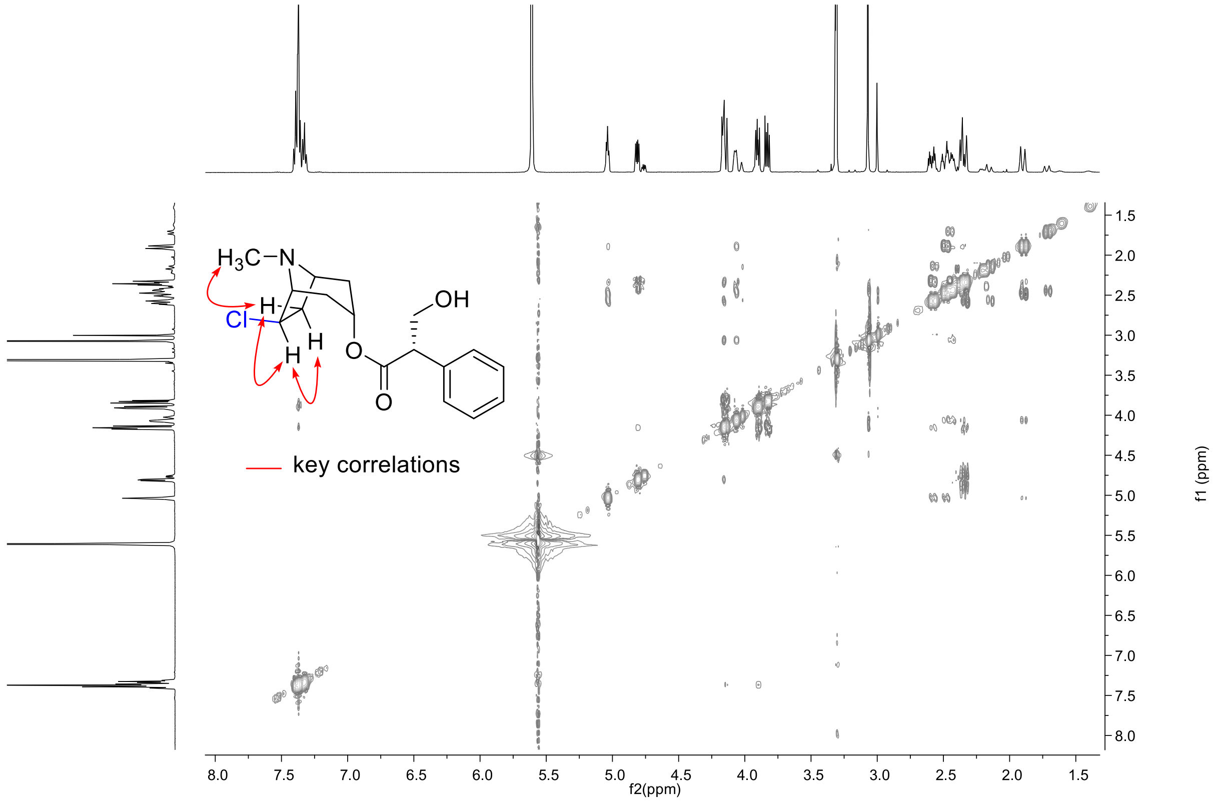

**Figure S23.** ROESY spectrum of 6β-chloro-hyoscyamine (500 MHz in $CD_3OD$ with 0.5 M DCl)

### Cartesian Coordinates of QM Region from QM/MM Calculations

**$RC_{ax}$**

| | | | |
|---|---|---|---|
| C | 14.2508391 | 8.7204736 | 11.9749303 |
| N | 13.9553407 | 7.6927422 | 11.1044139 |
| H | 14.5432154 | 6.8736622 | 10.8938001 |
| C | 12.7187460 | 7.8897873 | 10.6114665 |
| H | 12.2535203 | 7.2384325 | 9.8758369 |
| N | 12.1960849 | 9.0018984 | 11.1232717 |
| C | 13.1404275 | 9.5335610 | 11.9774583 |
| H | 12.9352984 | 10.4371873 | 12.5404936 |
| C | 10.1509360 | 12.8049889 | 14.0174150 |
| H | 9.5983070 | 12.8268619 | 14.9688111 |
| H | 9.8677031 | 13.7195853 | 13.4746140 |
| C | 9.6301450 | 11.6274540 | 13.1834464 |
| O | 10.5019572 | 10.8649049 | 12.6091423 |
| O | 8.4107179 | 11.4819885 | 13.0904373 |
| C | 10.0032390 | 6.2577153 | 13.9339343 |
| N | 10.1947820 | 7.3127357 | 14.8060477 |
| H | 10.3098975 | 7.2863844 | 15.8239378 |
| C | 10.1844176 | 8.4499724 | 14.0956872 |
| H | 10.3510648 | 9.4310155 | 14.5237173 |
| N | 9.9803345 | 8.1865434 | 12.8095973 |
| C | 9.8696566 | 6.8184222 | 12.6896139 |
| H | 9.7102917 | 6.3558740 | 11.7210691 |
| Fe | 10.1280965 | 9.5143572 | 11.2335080 |
| O | 9.9320135 | 10.6525299 | 10.0891078 |
| O | 10.4091932 | 8.4478542 | 8.2634269 |
| C | 9.5787354 | 7.8532881 | 8.9388005 |
| C | 8.7637924 | 6.7033224 | 8.3477083 |
| C | 7.3964118 | 7.1086177 | 7.7647582 |
| C | 6.7358156 | 5.9189387 | 7.0214817 |
| O | 6.8769663 | 5.8997375 | 5.7539723 |
| O | 6.1482373 | 5.0546953 | 7.6911977 |
| O | 9.3548781 | 8.0918788 | 10.2109406 |
| H | 8.6111354 | 5.9252972 | 9.1094010 |
| H | 9.3822838 | 6.2874778 | 7.5386849 |
| H | 7.5342714 | 7.9391122 | 7.0550889 |
| H | 6.7365930 | 7.4378406 | 8.5820061 |
| C | 11.8663513 | 12.6254230 | 8.0641931 |
| C | 12.5277410 | 12.4311984 | 9.4623268 |
| C | 13.9848286 | 12.0344012 | 9.1773787 |
| C | 14.9412602 | 13.2256116 | 9.0510486 |
| C | 14.5121691 | 14.2438625 | 7.9910633 |
| C | 13.8838202 | 13.5856577 | 6.7545809 |
| C | 13.0157137 | 12.3657108 | 7.0684468 |
| C | 13.4422942 | 16.3778058 | 8.0977433 |
| C | 13.3482793 | 9.9780908 | 7.8459230 |
| N | 13.8852818 | 11.3725868 | 7.8166182 |
| O | 13.5682459 | 15.1635834 | 8.6340288 |
| O | 13.9900057 | 16.7804041 | 7.1081164 |
| O | 10.8217891 | 11.7380716 | 7.7801091 |
| H | 11.5295534 | 13.6706870 | 7.9466709 |
| H | 11.9984493 | 11.6467653 | 10.0150597 |
| H | 12.4912578 | 13.3467041 | 10.0639918 |
| H | 15.9470378 | 12.8605959 | 8.7975748 |
| H | 15.0228824 | 13.7237523 | 10.0270869 |
| H | 14.7043324 | 13.2769523 | 6.0910811 |
| H | 13.3066060 | 14.3270118 | 6.1868728 |
| H | 10.4333051 | 11.3695621 | 8.6007372 |
| H | 13.3531239 | 9.6001396 | 6.8170659 |
| H | 14.0185940 | 9.3690801 | 8.4657139 |
| H | 12.3272074 | 9.9601707 | 8.2346682 |
| H | 15.3687786 | 14.8534322 | 7.6749824 |
| H | 12.6315881 | 11.8943009 | 6.1535975 |
| H | 14.3812107 | 11.3005341 | 9.8921329 |
| H | 14.8271832 | 11.3336904 | 7.3689056 |
| H | 10.0913581 | 5.2105417 | 14.2232952 |
| H | 15.1778145 | 8.7910787 | 12.5439634 |
| H | 11.2152168 | 12.8285808 | 14.2515452 |
| H | 12.7661640 | 16.9878665 | 8.6966933 |

**$TS_R$**

| | | | |
|---|---|---|---|
| C | 14.2006444 | 8.7116838 | 12.0362891 |
| N | 13.8737540 | 7.6395302 | 11.2320689 |
| H | 14.4512122 | 6.8078771 | 11.0446933 |
| C | 12.6225957 | 7.8117919 | 10.7763867 |
| H | 12.1160630 | 7.1071039 | 10.1239290 |
| N | 12.1156907 | 8.9540910 | 11.2411687 |
| C | 13.0920286 | 9.5261713 | 12.0326002 |
| H | 12.9054251 | 10.4600572 | 12.5495939 |
| C | 10.3510139 | 12.9481073 | 13.7267111 |
| H | 9.7150887 | 13.2356004 | 14.5744963 |
| H | 10.2453370 | 13.7345821 | 12.9615714 |

| | | | |
|---|---|---|---|
| C | 9.7887363 | 11.6568927 | 13.1272601 |
| O | 10.5949075 | 10.9526448 | 12.3862471 |
| O | 8.6137374 | 11.3775229 | 13.3489583 |
| C | 9.9134037 | 6.2156434 | 13.9347207 |
| N | 9.9783407 | 7.2707667 | 14.8249399 |
| H | 10.0793278 | 7.2398691 | 15.8439473 |
| C | 9.8739541 | 8.4133713 | 14.1229527 |
| H | 9.8866431 | 9.4006277 | 14.5714292 |
| N | 9.7447059 | 8.1504588 | 12.8306425 |
| C | 9.7679246 | 6.7811752 | 12.6925332 |
| H | 9.6956605 | 6.3199072 | 11.7119637 |
| Fe | 10.0475379 | 9.5037778 | 11.2127315 |
| O | 8.8791117 | 10.3717096 | 10.5114309 |
| O | 10.1113597 | 8.9155217 | 8.0012346 |
| C | 9.6661138 | 8.0203474 | 8.7312236 |
| C | 8.9451249 | 6.8139261 | 8.1359440 |
| C | 7.4960110 | 7.1324636 | 7.7104518 |
| C | 6.8237880 | 5.9402934 | 6.9820191 |
| O | 6.9476834 | 5.9204339 | 5.7121073 |
| O | 6.2332870 | 5.0830530 | 7.6569297 |
| O | 9.7851507 | 8.0076446 | 10.0196859 |
| H | 8.9406494 | 5.9906965 | 8.8640751 |
| H | 9.5203013 | 6.4946365 | 7.2529707 |
| H | 7.5060134 | 8.0000104 | 7.0324691 |
| H | 6.9078481 | 7.3842948 | 8.6061048 |
| C | 11.7076109 | 12.3947082 | 8.1494935 |
| C | 12.3910896 | 12.1443421 | 9.5271086 |
| C | 13.8642946 | 11.8459784 | 9.2203803 |
| C | 14.7571263 | 13.0825300 | 9.0692497 |
| C | 14.2913420 | 14.0589973 | 7.9843119 |
| C | 13.6621094 | 13.3575433 | 6.7689239 |
| C | 12.8434575 | 12.1117442 | 7.1221942 |
| C | 13.2297910 | 16.1961650 | 8.0675889 |
| C | 13.3291419 | 9.7586642 | 7.9011873 |
| N | 13.7832515 | 11.1754296 | 7.8593019 |
| O | 13.3486593 | 14.9862514 | 8.6190630 |
| O | 13.7514533 | 16.5681254 | 7.0523324 |
| O | 10.5091917 | 11.7229588 | 7.9302814 |
| H | 11.4569300 | 13.4590647 | 8.0578142 |
| H | 11.9118913 | 11.2926123 | 10.0230648 |
| H | 12.2940171 | 13.0016802 | 10.2046128 |
| H | 15.7798726 | 12.7569793 | 8.8289710 |
| H | 14.8120441 | 13.6062386 | 10.0334935 |
| H | 14.4803261 | 13.0701385 | 6.0930746 |
| H | 13.0480702 | 14.0688417 | 6.2034366 |
| H | 10.5287141 | 10.7626912 | 8.1123104 |
| H | 13.3097460 | 9.3770861 | 6.8737492 |
| H | 14.0443754 | 9.1792972 | 8.4990788 |
| H | 12.3300780 | 9.6822894 | 8.3382325 |
| H | 15.1338744 | 14.6709321 | 7.6371028 |
| H | 12.4601944 | 11.6165490 | 6.2204551 |
| H | 14.3155124 | 11.1386695 | 9.9296952 |
| H | 14.7208608 | 11.1899514 | 7.3982576 |
| H | 10.0424544 | 5.1705855 | 14.2162648 |
| H | 15.1383872 | 8.7884453 | 12.5865736 |
| H | 11.3830514 | 12.9241045 | 14.0765534 |
| H | 12.5913694 | 16.8348403 | 8.6779812 |

**$RC_{eq}$**

| | | | |
|---|---|---|---|
| C | 14.2010337 | 8.7172710 | 12.0451187 |
| N | 13.8746540 | 7.6458695 | 11.2395311 |
| H | 14.4555087 | 6.8179378 | 11.0451257 |
| C | 12.6187445 | 7.8097650 | 10.7957201 |
| H | 12.1100528 | 7.1057278 | 10.1448533 |
| N | 12.1067895 | 8.9469016 | 11.2705548 |
| C | 13.0864884 | 9.5239316 | 12.0545416 |
| H | 12.8981788 | 10.4579198 | 12.5704263 |
| C | 10.3665289 | 12.9643543 | 13.7121776 |
| H | 9.7277774 | 13.2483317 | 14.5589975 |
| H | 10.2710411 | 13.7544177 | 12.9501951 |
| C | 9.8062971 | 11.6755018 | 13.1066291 |
| O | 10.5967473 | 11.0100504 | 12.3097496 |
| O | 8.6501800 | 11.3680667 | 13.3794129 |
| C | 9.9046155 | 6.2189954 | 13.9309980 |
| N | 9.9633071 | 7.2748260 | 14.8201622 |
| H | 10.0634415 | 7.2454776 | 15.8393164 |
| C | 9.8563294 | 8.4173677 | 14.1168486 |
| H | 9.8634007 | 9.4058475 | 14.5633056 |
| N | 9.7287765 | 8.1516278 | 12.8256207 |
| C | 9.7610598 | 6.7828556 | 12.6872470 |
| H | 9.6935708 | 6.3203691 | 11.7067504 |
| Fe | 10.0689531 | 9.4851979 | 11.2189821 |
| O | 8.6274404 | 10.0925510 | 10.8269449 |
| O | 10.0695539 | 8.9509494 | 7.9469497 |
| C | 9.6822709 | 8.0340483 | 8.6886290 |
| C | 8.9569587 | 6.8207657 | 8.1133176 |
| C | 7.5025222 | 7.1342333 | 7.7023800 |
| C | 6.8259672 | 5.9420774 | 6.9790263 |

O 6.9491183 5.9192369 5.7087243
O 6.2330335 5.0874815 7.6551796
O 9.8677280 8.0133090 9.9629133
H 8.9629476 6.0058961 8.8508150
H 9.5213930 6.4914262 7.2272010
H 7.5047844 8.0015734 7.0241358
H 6.9233061 7.3860240 8.6039217
C 11.7080526 12.3900502 8.1630246
C 12.4015767 12.1418341 9.5358975
C 13.8720413 11.8440424 9.2209634
C 14.7615873 13.0820900 9.0664239
C 14.2896591 14.0571867 7.9828364
C 13.6553454 13.3542779 6.7708828
C 12.8374123 12.1081549 7.1280530
C 13.2300449 16.1949282 8.0684910
C 13.3294798 9.7559386 7.9041541
N 13.7827972 11.1729904 7.8601087
O 13.3485769 14.9845674 8.6197054
O 13.7516643 16.5665801 7.0531676
O 10.5051025 11.7178540 7.9643095
H 11.4545402 13.4536577 8.0721649
H 11.9264881 11.2899709 10.0355431
H 12.3065996 12.9988156 10.2140054
H 15.7845406 12.7594386 8.8238068
H 14.8171539 13.6057961 10.0306572
H 14.4713869 13.0650535 6.0929978
H 13.0408667 14.0655086 6.2059079
H 10.5415064 10.7443136 8.0634717
H 13.3090426 9.3731498 6.8771158
H 14.0451035 9.1770973 8.5019700
H 12.3306971 9.6800560 8.3423033
H 15.1303156 14.6691697 7.6311802
H 12.4523470 11.6134626 6.2266270
H 14.3273393 11.1366914 9.9275705
H 14.7181038 11.1881505 7.3941191
H 10.0361004 5.1743301 14.2128728
H 15.1410317 8.7921020 12.5918098
H 11.3959925 12.9320113 14.0688648
H 12.5917227 16.8339180 8.6786581

**TS1'**

C 14.2607751 8.7142247 11.9714399
N 13.9708337 7.6875467 11.0970494
H 14.5598498 6.8686614 10.8881162
C 12.7389497 7.8827710 10.5957354
H 12.2783941 7.2385140 9.8506629
N 12.2133848 8.9942733 11.1080060
C 13.1509249 9.5265704 11.9702413
H 12.9412856 10.4308188 12.5307588
C 10.1500781 12.8131706 14.0083342
H 9.5938797 12.8273699 14.9576915
H 9.8706840 13.7315268 13.4701289
C 9.6335865 11.6436560 13.1620373
O 10.5090745 10.9039526 12.5646240
O 8.4154302 11.4834294 13.0779757
C 10.0106091 6.2902240 13.9098724
N 10.2084161 7.3474692 14.7779919
H 10.3295290 7.3228133 15.7957358
C 10.1908702 8.4826764 14.0631982
H 10.3605575 9.4650251 14.4867467
N 9.9756926 8.2165477 12.7801138
C 9.8658970 6.8484311 12.6652020
H 9.6992265 6.3825164 11.6993044
Fe 10.1777332 9.5485559 11.1613918
O 10.0466584 10.7246207 9.9465810
O 10.4856777 8.3450848 8.2687376
C 9.5896511 7.8219861 8.9206600
C 8.7504793 6.6888869 8.3326009
C 7.3845239 7.1083061 7.7601206
C 6.7220774 5.9192902 7.0185748
O 6.8691126 5.8962387 5.7520037
O 6.1335563 5.0572010 7.6898746
O 9.3154036 8.1272578 10.1661159
H 8.5934025 5.9169600 9.1000576
H 9.3592427 6.2627319 7.5222128
H 7.5237082 7.9383708 7.0501874
H 6.7303019 7.4398024 8.5805383
C 11.8529178 12.6091505 8.1558820
C 12.5887560 12.4482205 9.5325341
C 14.0320469 12.0494531 9.1966273
C 14.9757536 13.2445990 9.0296752
C 14.5033798 14.2486081 7.9743448
C 13.8316986 13.5766978 6.7680424
C 12.9862013 12.3503253 7.1188112
C 13.4334545 16.3799582 8.0941889
C 13.3655628 9.9775402 7.8941469
N 13.8900498 11.3756733 7.8445195
O 13.5793049 15.1722936 8.6388332

| | | | |
|---|---|---|---|
| O | 13.9606338 | 16.7786446 | 7.0917459 |
| O | 10.8310012 | 11.7154188 | 7.9358391 |
| H | 11.5142729 | 13.6562303 | 8.0331997 |
| H | 12.0838412 | 11.6791132 | 10.1236619 |
| H | 12.5709440 | 13.3809557 | 10.1080789 |
| H | 15.9769469 | 12.8872321 | 8.7510133 |
| H | 15.0801482 | 13.7527631 | 9.9985857 |
| H | 14.6291994 | 13.2674198 | 6.0763004 |
| H | 13.2297124 | 14.3104402 | 6.2172955 |
| H | 10.3795316 | 11.1013569 | 8.9748317 |
| H | 13.3712336 | 9.5870479 | 6.8697151 |
| H | 14.0413653 | 9.3799221 | 8.5190370 |
| H | 12.3446076 | 9.9631942 | 8.2832089 |
| H | 15.3455570 | 14.8567779 | 7.6193537 |
| H | 12.5824014 | 11.8687000 | 6.2181896 |
| H | 14.4572554 | 11.3213869 | 9.9009612 |
| H | 14.8185080 | 11.3428735 | 7.3685539 |
| H | 10.0916682 | 5.2434783 | 14.2028230 |
| H | 15.1853374 | 8.7866982 | 12.5441520 |
| H | 11.2141767 | 12.8313309 | 14.2437733 |
| H | 12.7598332 | 16.9884029 | 8.6975929 |

**IC1'**

| | | | |
|---|---|---|---|
| C | 14.3684278 | 8.7000790 | 12.0216627 |
| N | 14.0535285 | 7.6335821 | 11.2069950 |
| H | 14.6286720 | 6.7992133 | 11.0255027 |
| C | 12.8186474 | 7.8160203 | 10.7141147 |
| H | 12.3227851 | 7.1175242 | 10.0442431 |
| N | 12.3167356 | 8.9637968 | 11.1669194 |
| C | 13.2711674 | 9.5290091 | 11.9884586 |
| H | 13.0740903 | 10.4606321 | 12.5085985 |
| C | 10.2268209 | 12.8623924 | 14.0127922 |
| H | 9.6613671 | 12.9104146 | 14.9552073 |
| H | 9.9669819 | 13.7700446 | 13.4470708 |
| C | 9.7126467 | 11.6716749 | 13.1941759 |
| O | 10.5778035 | 10.9510199 | 12.5619905 |
| O | 8.4933139 | 11.4882804 | 13.1638699 |
| C | 9.9714106 | 6.2550149 | 13.9181986 |
| N | 10.1346863 | 7.3125462 | 14.7933752 |
| H | 10.2517211 | 7.2858476 | 15.8106788 |
| C | 10.0873363 | 8.4512419 | 14.0816100 |
| H | 10.2203439 | 9.4356032 | 14.5157730 |
| N | 9.8899486 | 8.1892384 | 12.7958455 |
| C | 9.8181066 | 6.8179270 | 12.6757648 |
| H | 9.6759964 | 6.3574685 | 11.7027190 |
| Fe | 10.3137554 | 9.5384324 | 11.1663675 |
| O | 10.0555606 | 10.7511508 | 9.8264797 |
| O | 9.7285323 | 8.8290825 | 7.9750312 |
| C | 9.4136667 | 7.9327472 | 8.7748506 |
| C | 8.7286169 | 6.6805276 | 8.2425029 |
| C | 7.2994977 | 6.9826197 | 7.7497784 |
| C | 6.6702940 | 5.7877415 | 7.0027341 |
| O | 6.8265849 | 5.7788355 | 5.7358230 |
| O | 6.0836222 | 4.9142345 | 7.6606464 |
| O | 9.6080673 | 7.9952147 | 10.0512698 |
| H | 8.7113774 | 5.8929489 | 9.0082731 |
| H | 9.3295641 | 6.3290012 | 7.3887268 |
| H | 7.3422449 | 7.8443693 | 7.0664956 |
| H | 6.6692830 | 7.2395790 | 8.6148621 |
| C | 11.8897788 | 12.8654271 | 7.9803068 |
| C | 12.4945791 | 12.6209011 | 9.4184255 |
| C | 13.9369329 | 12.1529804 | 9.1690165 |
| C | 14.9556412 | 13.2931549 | 9.0545229 |
| C | 14.5911925 | 14.3261429 | 7.9853790 |
| C | 13.9809062 | 13.6860042 | 6.7299663 |
| C | 13.0563867 | 12.5024121 | 7.0243606 |
| C | 13.5364890 | 16.4773928 | 8.0681780 |
| C | 13.2197778 | 10.1173780 | 7.8520408 |
| N | 13.8410528 | 11.4776449 | 7.8149384 |
| O | 13.6537539 | 15.2630516 | 8.6084160 |
| O | 14.1024657 | 16.8791381 | 7.0894281 |
| O | 10.8020177 | 12.0692883 | 7.7894278 |
| H | 11.6337406 | 13.9357408 | 7.8689030 |
| H | 11.8745705 | 11.8680236 | 9.9186411 |
| H | 12.4725178 | 13.5441425 | 10.0061028 |
| H | 15.9455485 | 12.8757981 | 8.8193304 |
| H | 15.0461027 | 13.7905535 | 10.0298922 |
| H | 14.8085450 | 13.3420065 | 6.0937183 |
| H | 13.4483877 | 14.4444622 | 6.1405837 |
| H | 9.8878831 | 10.3091971 | 8.9589418 |
| H | 13.1577822 | 9.7487383 | 6.8217682 |
| H | 13.8733539 | 9.4624725 | 8.4418317 |
| H | 12.2201211 | 10.1680254 | 8.2921016 |
| H | 15.4735844 | 14.9122988 | 7.6960202 |
| H | 12.6865152 | 12.0358893 | 6.1005452 |
| H | 14.2734768 | 11.4092552 | 9.9042752 |
| H | 14.7916812 | 11.3789815 | 7.3923678 |
| H | 10.0759087 | 5.2090307 | 14.2064072 |

H 15.2880675 8.7684648 12.6027421
H 11.2884604 12.8708859 14.2596049
H 12.8446646 17.0800278 8.6566312

**TS1**

C 14.3504483 8.6826725 12.0175156
N 14.0885097 7.6688331 11.1211375
H 14.6932935 6.8666845 10.8966378
C 12.8523207 7.8390010 10.6232598
H 12.4052443 7.2024549 9.8627425
N 12.2986800 8.9220959 11.1630443
C 13.2189658 9.4633879 12.0362386
H 12.9814403 10.3491073 12.6158505
C 10.1749990 12.8233474 14.0614451
H 9.6294059 12.8260303 15.0171671
H 9.8913073 13.7522089 13.5435473
C 9.6483517 11.6659637 13.1990391
O 10.5059860 10.9419460 12.5652930
O 8.4255311 11.5179418 13.1358061
C 9.9890256 6.2689430 13.8878745
N 10.1672252 7.3226924 14.7638604
H 10.2735615 7.2921674 15.7824131
C 10.1545301 8.4620561 14.0517147
H 10.3072772 9.4432003 14.4860859
N 9.9640378 8.2047108 12.7645420
C 9.8605900 6.8369623 12.6437073
H 9.7054593 6.3683743 11.6758613
Fe 10.3128817 9.5587292 11.1129894
O 10.5218807 10.8142055 9.8607448
O 10.6281464 8.1986830 8.3074436
C 9.6107161 7.8330577 8.8950278
C 8.7511806 6.7175619 8.2986083
C 7.3432189 7.1000615 7.8095537
C 6.6951480 5.9189053 7.0409962
O 6.8688889 5.9035759 5.7767136
O 6.0866588 5.0532770 7.6902481
O 9.2422330 8.2981576 10.0602773
H 8.6575290 5.9125499 9.0458663
H 9.3286313 6.3250710 7.4503213
H 7.4128523 7.9648838 7.1305515
H 6.7153716 7.3671194 8.6718736
C 11.7383477 12.6764319 7.8321762
C 12.2852217 12.4360847 9.2578611
C 13.7204484 11.9464164 9.1132993
C 14.7429278 13.0967679 9.1097913
C 14.4465143 14.1622409 8.0499601
C 13.8915169 13.5775146 6.7408583
C 12.9596391 12.3799770 6.9358037
C 13.4551693 16.3363930 8.1471625
C 13.1464073 9.9558526 7.6815822
N 13.7245324 11.3329045 7.7282182
O 13.4998652 15.1028161 8.6558298
O 14.0674144 16.7348084 7.1950624
O 10.6542536 11.8708271 7.4723791
H 11.4725752 13.7422424 7.7228050
H 11.5066672 11.5647700 9.7556353
H 12.1655130 13.2567972 9.9745580
H 15.7432941 12.6780756 8.9242720
H 14.7705317 13.5639024 10.1033532
H 14.7469981 13.2723662 6.1220781
H 13.3796270 14.3626885 6.1689782
H 10.3480543 11.3743355 8.2662730
H 13.1108597 9.6384265 6.6338805
H 13.8137786 9.2924416 8.2461095
H 12.1435633 9.9359364 8.1139522
H 15.3538801 14.7382548 7.8256910
H 12.6387450 11.9432627 5.9806872
H 14.0079986 11.1794783 9.8471854
H 14.6998312 11.2787459 7.3570311
H 10.0784645 5.2235504 14.1832085
H 15.2674091 8.7606914 12.6016072
H 11.2410125 12.8377441 14.2883261
H 12.7810419 16.9621204 8.7320425

**IC1**

C 14.3681290 8.7141400 11.9972184
N 14.0543964 7.6501772 11.1783121
H 14.6261317 6.8118825 11.0025048
C 12.8285443 7.8425228 10.6692232
H 12.3337336 7.1471598 9.9955567
N 12.3300498 8.9958088 11.1153794
C 13.2793077 9.5527203 11.9499953
H 13.0862796 10.4868308 12.4668803
C 10.2414247 12.8744885 13.9706401
H 9.6619609 12.9261084 14.9039396
H 9.9957528 13.7825144 13.3992791
C 9.7365452 11.6879782 13.1431197
O 10.6053076 11.0134413 12.4602102

O 8.5245823 11.4649262 13.1409020
C 9.9897492 6.3040387 13.9051309
N 10.1634702 7.3668477 14.7720615
H 10.2826025 7.3446553 15.7896392
C 10.1239627 8.5007136 14.0522718
H 10.2605298 9.4867258 14.4816869
N 9.9219058 8.2307615 12.7679375
C 9.8389842 6.8579740 12.6591457
H 9.6927285 6.3909102 11.6903963
Fe 10.3298603 9.5549795 11.1212724
O 9.9323583 10.6872655 9.7258023
O 9.6122214 8.7862612 7.9602757
C 9.3825672 7.8721570 8.7744802
C 8.7071626 6.6100511 8.2642615
C 7.2715073 6.9062787 7.7867124
C 6.6450049 5.7201910 7.0261459
O 6.7945445 5.7315683 5.7587458
O 6.0661424 4.8355155 7.6759114
O 9.6537241 7.9459868 10.0332309
H 8.7102915 5.8254463 9.0327920
H 9.3003051 6.2635052 7.4030638
H 7.3044773 7.7776668 7.1160467
H 6.6459360 7.1467730 8.6596892
C 11.9579412 12.9679039 8.0910679
C 12.6170049 12.6656602 9.4100029
C 14.0036287 12.1591424 9.1855695
C 15.0607261 13.2835445 9.0961133
C 14.7012821 14.3562508 8.0637414
C 14.0451187 13.7793665 6.7991308
C 13.0783469 12.6275198 7.0771929
C 13.6547151 16.5028535 8.1843236
C 13.2115967 10.2046436 7.7959199
N 13.8684647 11.5455000 7.7973527
O 13.8007433 15.2965866 8.7320468
O 14.2022628 16.9087589 7.1955987
O 10.8056545 12.2201736 7.8279375
H 11.7307437 14.0491343 8.0053780
H 9.7406600 10.1134036 8.9308669
H 12.1271104 12.7219989 10.3816182
H 16.0359210 12.8454274 8.8382623
H 15.1667588 13.7488550 10.0848568
H 14.8481092 13.4235344 6.1380824
H 13.5350949 14.5767017 6.2426525
H 10.4599974 11.8124770 8.6632954
H 13.1160908 9.8778790 6.7542532
H 13.8538122 9.5065832 8.3476385
H 12.2241411 10.2764863 8.2600406
H 15.5954484 14.9259413 7.7769995
H 12.6545085 12.2079854 6.1548870
H 14.3279877 11.3835054 9.8946245
H 14.8067226 11.4449154 7.3499553
H 10.0804792 5.2577498 14.1968750
H 15.2838722 8.7779799 12.5849309
H 11.2995653 12.8782050 14.2321584
H 12.9526526 17.0979297 8.7683269

**TS2**

C 14.3407268 8.6985110 12.0124660
N 14.0022850 7.6277072 11.2126971
H 14.5611023 6.7813581 11.0380532
C 12.7671397 7.8282431 10.7250580
H 12.2543855 7.1469932 10.0506399
N 12.2918467 8.9888632 11.1698882
C 13.2571040 9.5443465 11.9781537
H 13.0861248 10.4887488 12.4813455
C 10.3267337 12.9989070 13.8560490
H 9.7084990 13.1206845 14.7561137
H 10.1538187 13.8822234 13.2225930
C 9.8195519 11.7902581 13.0665659
O 10.6658340 11.1977389 12.2807812
O 8.6365215 11.4750398 13.1870639
C 9.9876840 6.4688047 13.7238228
N 10.1212544 7.5289510 14.6022458
H 10.2255481 7.4932671 15.6217845
C 10.0715415 8.6684504 13.8881172
H 10.1559924 9.6517053 14.3345534
N 9.9038031 8.4078004 12.5968787
C 9.8520611 7.0327992 12.4776578
H 9.7364978 6.5674480 11.5035434
Fe 10.3654667 9.7690315 10.9277582
O 9.1242762 10.9377637 9.8869230
O 8.9066589 9.1101761 7.9697446
C 9.1462210 8.1249247 8.6854930
C 8.6216586 6.7631508 8.2379157
C 7.1347431 6.8598495 7.8487060
C 6.5951736 5.6473634 7.0686554
O 6.7707917 5.6756224 5.8037749
O 6.0309937 4.7374497 7.6962557

O 9.7920557 8.1848119 9.8050990
H 8.7930189 6.0054883 9.0155149
H 9.2112833 6.4673600 7.3531205
H 7.0167262 7.7514487 7.2166073
H 6.5373386 6.9893008 8.7628857
C 12.1608420 12.1827958 9.0035927
C 13.1800567 12.3722249 10.0883017
C 14.5593535 12.1625097 9.5640749
C 15.2384514 13.4673020 9.0977395
C 14.4085464 14.2328264 8.0607462
C 13.5880732 13.3306720 7.1206844
C 13.0719671 12.0308024 7.7481729
C 13.2251001 16.2886953 8.1567640
C 14.0895380 9.8816455 8.5389543
N 14.2956228 11.3365246 8.3117586
O 13.5270611 15.1500421 8.7851553
O 13.5765976 16.6070147 7.0530917
O 11.3449984 11.0637998 9.2143912
H 11.5439504 13.0924973 8.8664095
H 8.7718424 10.3904978 9.1416370
H 12.9370156 12.6173914 11.1179048
H 16.2217506 13.2290576 8.6663665
H 15.4175413 14.1096801 9.9694889
H 14.2343892 13.0619598 6.2706063
H 12.7577432 13.9052740 6.6952153
H 10.1446083 11.2479880 9.3936817
H 13.9771104 9.4127008 7.5543655
H 14.9646280 9.4675300 9.0551742
H 13.1756734 9.7458792 9.1179642
H 15.0659375 14.8598583 7.4461187
H 12.6042766 11.3867875 6.9922907
H 15.2263323 11.6004059 10.2333911
H 15.0938576 11.4368686 7.6482841
H 10.0438450 5.4246669 14.0315388
H 15.2644435 8.7659741 12.5871515
H 11.3717798 12.9724849 14.1646712
H 12.6039982 16.9273756 8.7847664

**IC2**

C 14.5315727 8.6237409 12.1116050
N 14.2047917 7.5374498 11.3314300
H 14.7686728 6.6953811 11.1614145
C 12.9649629 7.7129203 10.8451418
H 12.4623215 7.0181007 10.1765345
N 12.4772843 8.8722240 11.2715032
C 13.4347077 9.4535023 12.0671917
H 13.2521199 10.4036462 12.5560205
C 10.3333748 12.8955000 13.9677905
H 9.7647676 12.6221396 14.8714983
H 10.0020112 13.9071754 13.6996965
C 9.9183733 11.9806683 12.8109114
O 10.7721210 11.1966302 12.2817081
O 8.7253875 12.0832529 12.4486704
C 10.0895050 6.4851442 13.7361238
N 10.2759105 7.5482784 14.5994295
H 10.3775257 7.5180961 15.6184721
C 10.2706859 8.6807751 13.8632975
H 10.4403859 9.6674596 14.2824225
N 10.0811440 8.4096770 12.5809459
C 9.9682199 7.0405160 12.4831528
H 9.8297763 6.5661299 11.5153910
Fe 10.5930927 9.7058583 10.7915005
O 8.5852996 10.2560715 10.6200776
O 8.8510639 9.2133525 8.1634433
C 9.2168570 8.1586776 8.7076516
C 8.6627986 6.8258222 8.1972835
C 7.1773349 6.9359272 7.8190387
C 6.6229215 5.7272997 7.0435319
O 6.8011612 5.7423792 5.7781246
O 6.0445676 4.8275933 7.6740365
O 10.0436401 8.1026024 9.6982619
H 8.8357083 6.0395565 8.9462893
H 9.2544065 6.5465000 7.3072775
H 7.0583137 7.8335009 7.1949388
H 6.5876439 7.0673711 8.7382227
C 12.0513518 11.9434912 9.1576715
C 13.1319869 12.2147245 10.1647348
C 14.4830656 12.0876072 9.5490154
C 15.0315472 13.4318598 9.0265582
C 14.0992865 14.1094335 8.0128312
C 13.2539197 13.1349581 7.1671997
C 12.8866975 11.8133059 7.8462725
C 12.8279351 16.1097870 8.0635523
C 14.1186237 9.7629385 8.5847620
N 14.1978487 11.2245898 8.3274117
O 13.2435867 15.0434151 8.7506238
O 13.0140240 16.3155794 6.8940798
O 11.3003907 10.8094579 9.4520699

| | | | |
|---|---|---|---|
| H | 11.3796112 | 12.8164003 | 9.0406164 |
| H | 8.4692108 | 10.3536332 | 9.6507343 |
| H | 12.9513472 | 12.4656238 | 11.2054594 |
| H | 16.0128494 | 13.2643942 | 8.5587922 |
| H | 15.1933435 | 14.1047197 | 9.8789971 |
| H | 13.8292849 | 12.8979889 | 6.2590096 |
| H | 12.3459557 | 13.6415757 | 6.8260431 |
| H | 8.4668903 | 11.0907573 | 11.1533236 |
| H | 13.9837927 | 9.2673730 | 7.6164710 |
| H | 15.0483306 | 9.4221065 | 9.0569933 |
| H | 13.2562080 | 9.5682658 | 9.2209751 |
| H | 14.6899036 | 14.7238865 | 7.3224943 |
| H | 12.4150198 | 11.1225762 | 7.1356753 |
| H | 15.2324723 | 11.5859248 | 10.1783220 |
| H | 14.9458992 | 11.3746083 | 7.6141601 |
| H | 10.1138134 | 5.4413042 | 14.0489724 |
| H | 15.4490891 | 8.7001812 | 12.6950325 |
| H | 11.3895753 | 12.8932891 | 14.2370541 |
| H | 12.2910360 | 16.8103881 | 8.7030556 |

**TS3**

| | | | |
|---|---|---|---|
| C | 14.5775149 | 8.6216483 | 12.1190858 |
| N | 14.2455120 | 7.5266389 | 11.3538253 |
| H | 14.8076521 | 6.6838582 | 11.1855030 |
| C | 13.0003333 | 7.6955584 | 10.8776291 |
| H | 12.4905310 | 6.9925137 | 10.2232086 |
| N | 12.5143484 | 8.8572413 | 11.2951560 |
| C | 13.4787736 | 9.4492975 | 12.0731842 |
| H | 13.2969416 | 10.4041753 | 12.5553858 |
| C | 10.3310061 | 12.9138369 | 14.0014737 |
| H | 9.7710356 | 12.6052720 | 14.8993990 |
| H | 9.9895091 | 13.9304240 | 13.7679798 |
| C | 9.9292012 | 12.0301855 | 12.8151622 |
| O | 10.7904998 | 11.2499968 | 12.2933572 |
| O | 8.7487737 | 12.1542472 | 12.4205293 |
| C | 10.0811230 | 6.4672215 | 13.7447715 |
| N | 10.2634817 | 7.5291011 | 14.6109538 |
| H | 10.3681102 | 7.4978393 | 15.6292266 |
| C | 10.2455448 | 8.6635980 | 13.8775198 |
| H | 10.4099551 | 9.6515781 | 14.2972619 |
| N | 10.0510555 | 8.3944178 | 12.5959573 |
| C | 9.9496856 | 7.0254628 | 12.4938507 |
| H | 9.8150628 | 6.5560480 | 11.5226758 |
| Fe | 10.6260255 | 9.7003518 | 10.8652839 |
| O | 8.6755014 | 10.3534727 | 10.5372040 |
| O | 8.9672258 | 9.2379148 | 8.1298016 |
| C | 9.2918385 | 8.1694037 | 8.6927311 |
| C | 8.6986167 | 6.8527269 | 8.1754455 |
| C | 7.2122070 | 6.9816502 | 7.8063110 |
| C | 6.6421286 | 5.7735448 | 7.0388182 |
| O | 6.8220579 | 5.7753856 | 5.7728815 |
| O | 6.0533794 | 4.8834351 | 7.6733948 |
| O | 10.0924243 | 8.0873303 | 9.6885255 |
| H | 8.8612663 | 6.0611873 | 8.9212435 |
| H | 9.2774100 | 6.5630872 | 7.2805684 |
| H | 7.1003196 | 7.8786653 | 7.1793677 |
| H | 6.6285595 | 7.1226753 | 8.7283758 |
| C | 11.9159863 | 11.9859660 | 8.8863128 |
| C | 12.8438358 | 12.2212381 | 9.9977760 |
| C | 14.2529508 | 11.9941975 | 9.5468371 |
| C | 14.8995106 | 13.3304800 | 9.1106819 |
| C | 14.0875154 | 14.0636154 | 8.0314330 |
| C | 13.3097987 | 13.1405666 | 7.0675979 |
| C | 12.8428696 | 11.8105062 | 7.6685781 |
| C | 12.8428649 | 16.0903377 | 8.0494426 |
| C | 14.0377856 | 9.6972534 | 8.4518878 |
| N | 14.0860999 | 11.1786686 | 8.2747026 |
| O | 13.1921384 | 14.9922938 | 8.7263340 |
| O | 13.0917669 | 16.3181959 | 6.8962429 |
| O | 11.5738462 | 10.7953449 | 9.5086393 |
| H | 11.0842535 | 12.6760366 | 8.6804685 |
| H | 8.5647226 | 10.3770470 | 9.5613992 |
| H | 12.5359956 | 12.4896923 | 11.0032865 |
| H | 15.9093974 | 13.1210991 | 8.7266670 |
| H | 15.0178344 | 13.9846872 | 9.9840387 |
| H | 13.9679719 | 12.9140624 | 6.2156183 |
| H | 12.4573472 | 13.6865376 | 6.6529972 |
| H | 8.5166168 | 11.1922851 | 11.0424834 |
| H | 13.9490511 | 9.2575066 | 7.4519605 |
| H | 14.9738418 | 9.3719972 | 8.9248221 |
| H | 13.1754095 | 9.4386564 | 9.0613521 |
| H | 14.7568719 | 14.6855177 | 7.4239704 |
| H | 12.4298852 | 11.1407777 | 6.9023125 |
| H | 14.8985724 | 11.4493306 | 10.2501716 |
| H | 14.8962026 | 11.3613258 | 7.6405339 |
| H | 10.1124153 | 5.4227708 | 14.0549498 |
| H | 15.4911247 | 8.6973607 | 12.7087060 |
| H | 11.3893575 | 12.9083024 | 14.2621052 |

H 12.3008811 16.7912734 8.6842731

**PC**

C 14.4641720 8.7190503 12.0381392
N 14.1713567 7.6516081 11.2158138
H 14.7460633 6.8154641 11.0516495
C 12.9508305 7.8424859 10.6839734
H 12.4635250 7.1458588 10.0051235
N 12.4390400 8.9915115 11.1142431
C 13.3675054 9.5483350 11.9684084
H 13.1519187 10.4780137 12.4857774
C 10.2830094 12.9203178 13.9170283
H 9.7142358 12.5271374 14.7768165
H 9.9209037 13.9404941 13.7429742
C 9.9708458 12.0877892 12.6587179
O 10.7554276 11.1463438 12.3317417
O 8.9318355 12.4002442 12.0302693
C 10.0477489 6.3018276 13.9304917
N 10.2196106 7.3696628 14.7928500
H 10.3335830 7.3501609 15.8100759
C 10.1895525 8.5005851 14.0597263
H 10.3351896 9.4967997 14.4690757
N 9.9981224 8.2187961 12.7801270
C 9.9118251 6.8488174 12.6787430
H 9.7788047 6.3811771 11.7060082
Fe 10.3769056 9.4335189 11.0960725
O 8.7943189 10.4569142 10.2481901
O 9.4615304 9.0186594 8.0156905
C 9.4068387 7.9554955 8.6838277
C 8.7186781 6.7364852 8.0716327
C 7.2555358 7.0173831 7.6928323
C 6.6129394 5.8389117 6.9407679
O 6.8368486 5.7904854 5.6832063
O 5.9508015 5.0066379 7.5807497
O 9.8660996 7.8169951 9.8612001
H 8.7784513 5.8927646 8.7729336
H 9.2756729 6.4572692 7.1629608
H 7.2322030 7.9107243 7.0494793
H 6.6786283 7.2198876 8.6079174
C 11.8569802 12.5655783 8.1947538
C 12.4637972 12.3254554 9.5060401
C 13.8772263 11.8263072 9.2514077
C 14.8375420 13.0204860 9.1440057
C 14.4298579 14.0445963 8.0692609
C 13.7615130 13.4334473 6.8196797
C 12.8872338 12.2114814 7.1336703
C 13.4594925 16.2252300 8.1799640
C 13.3186634 9.7858665 7.7831229
N 13.7733902 11.2140706 7.8611426
O 13.5279917 15.0019089 8.7137993
O 14.0355034 16.6021025 7.1967186
O 11.4225059 11.4889815 9.0198814
H 11.1386330 13.3600511 7.9888827
H 8.9837871 10.4110816 9.2853500
H 12.2518837 12.9106914 10.4037003
H 15.8475223 12.6486376 8.9209871
H 14.8987306 13.5300520 10.1153125
H 14.5489742 13.1354134 6.1143079
H 13.1790616 14.2049919 6.2991513
H 8.7990710 11.3655844 10.6541586
H 13.4305120 9.4767975 6.7374048
H 13.9822659 9.1865705 8.4184897
H 12.2813962 9.6964688 8.1128470
H 15.3109284 14.6139484 7.7468424
H 12.4831478 11.7397570 6.2264606
H 14.2318955 11.0625984 9.9555391
H 14.7192462 11.2393364 7.4169676
H 10.1291715 5.2551806 14.2236942
H 15.3691541 8.7900907 12.6414863
H 11.3324892 12.9190063 14.2114052
H 12.7946562 16.8607334 8.7649386

**TS$_{OH}$**

C 14.4482740 8.6626743 12.0484624
N 14.1631883 7.5966911 11.2262356
H 14.7559859 6.7799688 11.0319831
C 12.9159356 7.7432005 10.7473400
H 12.4471374 7.0356111 10.0674128
N 12.3809354 8.8624345 11.2188206
C 13.3221308 9.4535874 12.0321549
H 13.1043889 10.3818892 12.5505266
C 10.2265421 12.8518259 13.9910511
H 9.6554645 12.8416962 14.9307716
H 9.9596280 13.7852221 13.4725071
C 9.7357576 11.7021250 13.0997982
O 10.6155091 11.0971210 12.3825741
O 8.5228034 11.4640885 13.0843831
C 10.0528996 6.3813144 13.8916104

| | | | |
|---|---|---|---|
| N | 10.2617345 | 7.4335685 | 14.7645124 |
| H | 10.3707926 | 7.4006667 | 15.7830316 |
| C | 10.2632739 | 8.5726339 | 14.0517196 |
| H | 10.4340873 | 9.5529447 | 14.4824193 |
| N | 10.0509340 | 8.3134853 | 12.7685138 |
| C | 9.9201933 | 6.9471353 | 12.6490290 |
| H | 9.7499015 | 6.4903512 | 11.6790021 |
| Fe | 10.3990369 | 9.5526213 | 11.1711678 |
| O | 10.7615334 | 10.6672009 | 9.7292202 |
| O | 9.9590518 | 8.7959346 | 7.9942695 |
| C | 9.4409967 | 8.0123295 | 8.8098071 |
| C | 8.7248005 | 6.7685924 | 8.2951911 |
| C | 7.3246883 | 7.0781496 | 7.7274413 |
| C | 6.7051529 | 5.8544774 | 7.0103884 |
| O | 6.8405606 | 5.8261363 | 5.7420873 |
| O | 6.1470540 | 4.9811144 | 7.6925541 |
| O | 9.4650357 | 8.1810482 | 10.0875619 |
| H | 8.6449893 | 6.0169461 | 9.0926117 |
| H | 9.3510411 | 6.3613922 | 7.4857692 |
| H | 7.4074634 | 7.9049724 | 7.0054397 |
| H | 6.6638381 | 7.3848161 | 8.5527441 |
| C | 11.7226839 | 12.7504324 | 7.9302429 |
| C | 12.2585102 | 12.3256756 | 9.2752896 |
| C | 13.6799275 | 11.8810368 | 9.1152804 |
| C | 14.7096388 | 13.0380299 | 9.1707634 |
| C | 14.4498792 | 14.1466208 | 8.1418729 |
| C | 13.8610797 | 13.6369786 | 6.8129566 |
| C | 12.9032562 | 12.4538910 | 6.9749664 |
| C | 13.5535658 | 16.3533134 | 8.2693516 |
| C | 13.1473228 | 9.9820767 | 7.5480832 |
| N | 13.6890870 | 11.3625079 | 7.6887065 |
| O | 13.5503889 | 15.1167218 | 8.7724565 |
| O | 14.1990153 | 16.7371658 | 7.3314605 |
| O | 10.5474854 | 12.0847871 | 7.5339740 |
| H | 11.5435563 | 13.8425221 | 7.9201203 |
| H | 10.5446141 | 10.0845057 | 8.9502861 |
| H | 11.9011312 | 12.7130483 | 10.2256509 |
| H | 15.7137466 | 12.6205431 | 8.9937761 |
| H | 14.7181140 | 13.4729123 | 10.1784625 |
| H | 14.6978519 | 13.3380068 | 6.1662397 |
| H | 13.3627634 | 14.4610045 | 6.2858197 |
| H | 10.1680626 | 11.7195553 | 8.3539272 |
| H | 13.0679666 | 9.7517540 | 6.4810653 |
| H | 13.8536559 | 9.2923052 | 8.0270875 |
| H | 12.1759520 | 9.9123520 | 8.0339862 |
| H | 15.3828979 | 14.6855457 | 7.9291703 |
| H | 12.5459654 | 12.0680716 | 6.0107884 |
| H | 13.9766808 | 11.0761476 | 9.8044436 |
| H | 14.6628842 | 11.3496239 | 7.3101864 |
| H | 10.1076444 | 5.3311157 | 14.1782335 |
| H | 15.3611975 | 8.7443505 | 12.6383496 |
| H | 11.2871258 | 12.8607418 | 14.2423473 |
| H | 12.8771378 | 16.9988878 | 8.8294819 |

**$PC_{OH}$**

| | | | |
|---|---|---|---|
| C | 14.4912138 | 8.6630122 | 12.1006487 |
| N | 14.1916137 | 7.5918541 | 11.2933641 |
| H | 14.7668234 | 6.7585787 | 11.1189950 |
| C | 12.9545851 | 7.7659617 | 10.7915401 |
| H | 12.4767832 | 7.0547305 | 10.1213910 |
| N | 12.4385926 | 8.9075761 | 11.2306637 |
| C | 13.3845162 | 9.4809704 | 12.0533570 |
| H | 13.1882776 | 10.4242860 | 12.5545039 |
| C | 10.2381729 | 12.8940238 | 14.0872189 |
| H | 9.6897009 | 12.8502250 | 15.0392648 |
| H | 9.9699253 | 13.8513145 | 13.6145655 |
| C | 9.7280645 | 11.7857789 | 13.1567295 |
| O | 10.5737425 | 11.2549367 | 12.3433894 |
| O | 8.5228683 | 11.5141612 | 13.2011200 |
| C | 10.0480230 | 6.3221178 | 13.9552261 |
| N | 10.2111144 | 7.3547697 | 14.8565404 |
| H | 10.3114338 | 7.3033329 | 15.8737625 |
| C | 10.1815106 | 8.5132994 | 14.1674907 |
| H | 10.3033656 | 9.4868166 | 14.6328681 |
| N | 9.9976289 | 8.2861977 | 12.8776211 |
| C | 9.9149666 | 6.9204660 | 12.7249103 |
| H | 9.7737750 | 6.4792535 | 11.7418553 |
| Fe | 10.4384275 | 9.5981850 | 11.2436092 |
| O | 10.8863233 | 10.7641345 | 9.4040224 |
| O | 9.9034531 | 8.9186166 | 7.9250922 |
| C | 9.4414557 | 8.0793924 | 8.7494681 |
| C | 8.7466241 | 6.8390795 | 8.2015073 |
| C | 7.3234152 | 7.1293010 | 7.6814957 |
| C | 6.6959273 | 5.9034248 | 6.9751594 |
| O | 6.8368646 | 5.8614855 | 5.7070685 |
| O | 6.1252058 | 5.0423145 | 7.6608815 |
| O | 9.5179450 | 8.2217812 | 10.0068011 |
| H | 8.7049449 | 6.0666493 | 8.9820652 |

| | | | |
|---|---|---|---|
| H | 9.3604490 | 6.4629163 | 7.3676763 |
| H | 7.3693877 | 7.9629243 | 6.9632845 |
| H | 6.6864805 | 7.4192440 | 8.5312470 |
| C | 11.4851926 | 12.3779590 | 7.6590981 |
| C | 11.8038240 | 11.7771814 | 9.0772368 |
| C | 13.2933222 | 11.3752574 | 8.9745509 |
| C | 14.2633514 | 12.5298395 | 9.2695097 |
| C | 14.1623851 | 13.7375668 | 8.3229621 |
| C | 13.7247194 | 13.3742306 | 6.8928528 |
| C | 12.7733627 | 12.1827198 | 6.8351312 |
| C | 13.3739382 | 15.9677593 | 8.4880736 |
| C | 12.9427938 | 9.6742024 | 7.1449105 |
| N | 13.4723130 | 11.0176088 | 7.5177020 |
| O | 13.2380644 | 14.7087950 | 8.9198839 |
| O | 14.1119610 | 16.3350639 | 7.6145202 |
| O | 10.4261355 | 11.7120984 | 7.0266933 |
| H | 11.2802675 | 13.4568153 | 7.7547224 |
| H | 10.6331648 | 10.0866707 | 8.6850373 |
| H | 11.7102910 | 12.5334632 | 9.8676777 |
| H | 15.2872498 | 12.1326417 | 9.1940952 |
| H | 14.1350801 | 12.8671116 | 10.3074856 |
| H | 14.6236052 | 13.1515046 | 6.3019343 |
| H | 13.2651719 | 14.2433696 | 6.4055810 |
| H | 9.7740038 | 11.4578188 | 7.6964448 |
| H | 13.2767039 | 9.4651347 | 6.1248071 |
| H | 13.3602422 | 8.9296975 | 7.8340535 |
| H | 11.8534611 | 9.6664199 | 7.1811109 |
| H | 15.1358147 | 14.2413844 | 8.2737717 |
| H | 12.5287066 | 11.8982129 | 5.8038614 |
| H | 13.5270660 | 10.5071727 | 9.6008066 |
| H | 14.4854010 | 11.0350000 | 7.2572754 |
| H | 10.1239229 | 5.2677331 | 14.2208936 |
| H | 15.4042235 | 8.7378113 | 12.6913140 |
| H | 11.3049675 | 12.8806139 | 14.3104602 |
| H | 12.7237662 | 16.6519042 | 9.0333159 |

**RC$^{L290F}$**

| | | | |
|---|---|---|---|
| C | 48.9926400 | 37.1288531 | 48.3158995 |
| N | 49.0718207 | 35.9607922 | 47.5872696 |
| H | 49.9207385 | 35.4177773 | 47.3923500 |
| C | 47.8417663 | 35.6541354 | 47.1314044 |
| H | 47.6225866 | 34.7929779 | 46.5035678 |
| N | 46.9740023 | 36.5701838 | 47.5435570 |
| C | 47.6699653 | 37.5004371 | 48.2778831 |
| H | 47.1551205 | 38.3440518 | 48.7208312 |
| C | 43.5585838 | 39.6758097 | 50.2162794 |
| H | 42.9872931 | 39.5493769 | 51.1476118 |
| H | 43.0141567 | 40.4128781 | 49.6091112 |
| C | 43.5222557 | 38.3683620 | 49.4136489 |
| O | 44.6274950 | 38.0313309 | 48.8135323 |
| O | 42.4714401 | 37.7413071 | 49.3317357 |
| C | 45.6603428 | 33.6226554 | 50.5665943 |
| N | 45.5281972 | 34.7482887 | 51.3560288 |
| H | 45.6530438 | 34.8260273 | 52.3700801 |
| C | 45.1900326 | 35.7736079 | 50.5572505 |
| H | 45.0423278 | 36.7881485 | 50.9035944 |
| N | 45.0834721 | 35.3662401 | 49.2985573 |
| C | 45.3791877 | 34.0212945 | 49.2839635 |
| H | 45.3630546 | 33.4590022 | 48.3551056 |
| C | 39.4987247 | 42.5322376 | 45.7142446 |
| H | 39.3529682 | 42.6993113 | 44.6319685 |
| H | 39.6984440 | 43.5258146 | 46.1480772 |
| C | 40.7044557 | 41.6546869 | 45.9874011 |
| C | 41.8667686 | 42.2634388 | 46.4835709 |
| H | 41.8705594 | 43.3438734 | 46.6506262 |
| C | 42.9999191 | 41.5103938 | 46.8007369 |
| H | 43.8788534 | 42.0019376 | 47.2275977 |
| C | 42.9938643 | 40.1285711 | 46.6221842 |
| H | 43.8566125 | 39.5260764 | 46.9001378 |
| C | 41.8604433 | 39.5068254 | 46.0880781 |
| H | 41.8749679 | 38.4260588 | 45.9250843 |
| C | 40.7295662 | 40.2637409 | 45.7703767 |
| H | 39.8537698 | 39.7626620 | 45.3499282 |
| Fe | 44.8726597 | 36.5374306 | 47.5966063 |
| O | 44.6909276 | 37.4785635 | 46.2827222 |
| O | 42.9310706 | 35.5868851 | 45.2428297 |
| C | 43.7474525 | 34.7687282 | 45.6357602 |
| C | 43.9719612 | 33.4510323 | 44.9081331 |
| C | 42.6547267 | 32.6997521 | 44.7160437 |
| C | 42.7269196 | 31.5357802 | 43.7181372 |
| O | 43.2686880 | 31.7600729 | 42.6098695 |
| O | 42.1692445 | 30.4558173 | 44.0617857 |
| O | 44.4776661 | 34.9181203 | 46.7234729 |
| H | 44.7165964 | 32.8442264 | 45.4435454 |
| H | 44.3844680 | 33.6850917 | 43.9120485 |
| H | 41.9249787 | 33.4167141 | 44.3046003 |
| H | 42.2639512 | 32.3343770 | 45.6771245 |
| C | 45.5012947 | 40.1118675 | 44.2100169 |

C 46.3613083 40.1820609 45.5083208
C 47.8250916 40.1801504 45.0370018
C 48.3854732 41.5692616 44.7085757
C 47.5366666 42.3451401 43.6997384
C 47.0035485 41.4391914 42.5836546
C 46.5476205 40.0642282 43.0745607
C 45.8277694 44.0171418 43.8986508
C 47.6191368 37.9244493 43.9097770
N 47.7395047 39.4018276 43.7387225
O 46.4325401 42.9606217 44.4407602
O 46.1056800 44.5423826 42.8544824
O 44.6655418 38.9866542 44.1367435
H 44.8878260 41.0201661 44.1039890
H 46.1544728 39.3065327 46.1364260
H 46.1442892 41.0756607 46.1021784
H 49.3978288 41.4573090 44.2934400
H 48.4919018 42.1416698 45.6392118
H 47.8153346 41.2991023 41.8568138
H 46.1895333 41.9364215 42.0387280
H 44.5962745 38.5341620 45.0051186
H 48.5357500 37.5620839 44.3937814
H 46.7405982 37.6838519 44.5149581
H 47.5200228 37.4793849 42.9115887
H 48.1132582 43.1671128 43.2553807
H 46.2085639 39.4337715 42.2416984
H 48.5108270 39.6531169 45.7132441
H 48.5992737 39.5685875 43.1702628
H 46.0154524 32.6621293 50.9398958
H 49.8253383 37.5618092 48.8701824
H 44.5477468 40.0609157 50.4639025
H 45.0301254 44.3587486 44.5582992
H 38.5521672 42.1567059 46.1029245

**TS1$^{L290F}$**

C 49.1188918 37.1636276 48.3575173
N 49.1992565 35.9862328 47.6455536
H 50.0480536 35.4443569 47.4505873
C 47.9730699 35.6690263 47.1927986
H 47.7538703 34.8042333 46.5700408
N 47.1077260 36.5890032 47.5978531
C 47.7973769 37.5328768 48.3163201
H 47.2804208 38.3862204 48.7365155
C 43.5794223 39.6818363 50.2187312
H 43.0085832 39.5394376 51.1477472
H 43.0326231 40.4248922 49.6207682
C 43.5504195 38.3883417 49.3947312
O 44.6513810 38.0985310 48.7568173
O 42.5165361 37.7363950 49.3252803
C 45.6701690 33.6682284 50.5012617
N 45.5304195 34.7891487 51.2961389
H 45.6419895 34.8577948 52.3124371
C 45.2093847 35.8211444 50.4984154
H 45.0581764 36.8327063 50.8511735
N 45.1216168 35.4240392 49.2354378
C 45.4114556 34.0779692 49.2161455
H 45.4108576 33.5218039 48.2829565
C 39.4839883 42.5326562 45.7110047
H 39.3375365 42.7059245 44.6298903
H 39.6909137 43.5220679 46.1507265
C 40.6811494 41.6432152 45.9793539
C 41.8460524 42.2362299 46.4879144
H 41.8594299 43.3151100 46.6642528
C 42.9699881 41.4694383 46.8035072
H 43.8537557 41.9484693 47.2343477
C 42.9507707 40.0897223 46.6115965
H 43.8099342 39.4813876 46.8823451
C 41.8149707 39.4846174 46.0653872
H 41.8196589 38.4055464 45.8903560
C 40.6934112 40.2545051 45.7483130
H 39.8164930 39.7655503 45.3164949
Fe 45.0444139 36.6552147 47.5279324
O 45.0897040 37.6943332 46.2183029
O 43.0164496 35.7255040 45.1971107
C 43.8373282 34.8961705 45.5632117
C 44.0113900 33.5543900 44.8593204
C 42.6892558 32.7982668 44.7118053
C 42.7424637 31.6115543 43.7313505
O 43.2736271 31.8137712 42.6135399
O 42.1847701 30.5382221 44.0954144
O 44.6302803 35.0659443 46.5976324
H 44.7669806 32.9538176 45.3864904
H 44.4025379 33.7621414 43.8489057
H 41.9473626 33.5045659 44.3031114
H 42.3196243 32.4492142 45.6874337
C 45.3892422 40.0402067 44.1438542
C 46.1978103 39.9276067 45.4560662
C 47.6693615 39.9495647 45.0482157
C 48.2576094 41.3561642 44.8327071

| | | | |
|---|---|---|---|
| C | 47.4874843 | 42.2068850 | 43.8183171 |
| C | 46.9739766 | 41.3790840 | 42.6316499 |
| C | 46.4679516 | 39.9936125 | 43.0349078 |
| C | 45.8278236 | 43.9288570 | 43.9787433 |
| C | 47.4876698 | 37.7887897 | 43.7545370 |
| N | 47.6315667 | 39.2704467 | 43.6928678 |
| O | 46.3779416 | 42.8479264 | 44.5308573 |
| O | 46.1520425 | 44.4484131 | 42.9445246 |
| O | 44.4343375 | 39.0124399 | 43.9813798 |
| H | 44.8575784 | 41.0016614 | 44.0861038 |
| H | 45.8002695 | 38.7764658 | 45.9859698 |
| H | 45.9645788 | 40.7022977 | 46.1954432 |
| H | 49.2929445 | 41.2519879 | 44.4749358 |
| H | 48.3099007 | 41.8731057 | 45.7991439 |
| H | 47.8058970 | 41.2619675 | 41.9233366 |
| H | 46.1913675 | 41.9267154 | 42.0896965 |
| H | 44.3804859 | 38.4974386 | 44.8098996 |
| H | 48.3778263 | 37.3815553 | 44.2528048 |
| H | 46.5858771 | 37.5246262 | 44.3141091 |
| H | 47.4266887 | 37.4133821 | 42.7255112 |
| H | 48.1220472 | 43.0198044 | 43.4405625 |
| H | 46.1379461 | 39.4124883 | 42.1635746 |
| H | 48.3342417 | 39.3760357 | 45.7094344 |
| H | 48.5132199 | 39.4695926 | 43.1674339 |
| H | 46.0093650 | 32.7052420 | 50.8829295 |
| H | 49.9429702 | 37.5930117 | 48.9272465 |
| H | 44.5678978 | 40.0666368 | 50.4695536 |
| H | 45.0253433 | 44.3000171 | 44.6161855 |
| H | 38.5371728 | 42.1582572 | 46.1001485 |

**IC1$^{L290F}$**

| | | | |
|---|---|---|---|
| C | 48.9479022 | 37.1364414 | 48.2591834 |
| N | 49.0232528 | 35.9671351 | 47.5280047 |
| H | 49.8658948 | 35.4044002 | 47.3605559 |
| C | 47.8063997 | 35.6910529 | 47.0288493 |
| H | 47.5816234 | 34.8306727 | 46.4019254 |
| N | 46.9432223 | 36.6310716 | 47.4094175 |
| C | 47.6382062 | 37.5415858 | 48.1766028 |
| H | 47.1264282 | 38.3904843 | 48.6138979 |
| C | 43.5683905 | 39.6876388 | 50.1573399 |
| H | 42.9703387 | 39.5874492 | 51.0745909 |
| H | 43.0557452 | 40.4250550 | 49.5232975 |
| C | 43.5371584 | 38.3663948 | 49.3784081 |
| O | 44.6256707 | 38.0182857 | 48.7574971 |
| O | 42.4876091 | 37.7285431 | 49.3402227 |
| C | 45.6250193 | 33.5349295 | 50.5388576 |
| N | 45.4651546 | 34.6668470 | 51.3146078 |
| H | 45.5953203 | 34.7602089 | 52.3260165 |
| C | 45.0928076 | 35.6704724 | 50.5020491 |
| H | 44.9217487 | 36.6858765 | 50.8377123 |
| N | 44.9915201 | 35.2458668 | 49.2479879 |
| C | 45.3262306 | 33.9096510 | 49.2522321 |
| H | 45.3287472 | 33.3382233 | 48.3282358 |
| C | 39.4602743 | 42.6018133 | 45.7322834 |
| H | 39.3102752 | 42.7749956 | 44.6514083 |
| H | 39.6414123 | 43.5953180 | 46.1746842 |
| C | 40.6790554 | 41.7423154 | 45.9986461 |
| C | 41.8468107 | 42.3649904 | 46.4639548 |
| H | 41.8489006 | 43.4481955 | 46.6117908 |
| C | 42.9880892 | 41.6212048 | 46.7760547 |
| H | 43.8735658 | 42.1221054 | 47.1781714 |
| C | 42.9834577 | 40.2365989 | 46.6244153 |
| H | 43.8450086 | 39.6301120 | 46.8967181 |
| C | 41.8418234 | 39.6036190 | 46.1248360 |
| H | 41.8655252 | 38.5211162 | 45.9933684 |
| C | 40.7049034 | 40.3482415 | 45.8102083 |
| H | 39.8233927 | 39.8336073 | 45.4195098 |
| Fe | 44.8693867 | 36.4994508 | 47.4922879 |
| O | 44.1129440 | 37.4169772 | 46.0855536 |
| O | 43.6357455 | 35.3557054 | 44.6108379 |
| C | 44.1785274 | 34.4951620 | 45.3258002 |
| C | 44.2077881 | 33.0564602 | 44.8519339 |
| C | 42.7878961 | 32.4876476 | 44.7573188 |
| C | 42.6608725 | 31.1037685 | 44.0935211 |
| O | 43.4192247 | 30.8261712 | 43.1302401 |
| O | 41.7452218 | 30.3648759 | 44.5375088 |
| O | 44.6943081 | 34.7476425 | 46.4906284 |
| H | 44.8312073 | 32.4640145 | 45.5353415 |
| H | 44.6608117 | 33.0264846 | 43.8482678 |
| H | 42.1812640 | 33.1817399 | 44.1497235 |
| H | 42.3163101 | 32.4353191 | 45.7499744 |
| C | 45.4721192 | 40.0969795 | 44.1752294 |
| C | 46.3297134 | 40.1770234 | 45.4119613 |
| C | 47.7770335 | 40.1194879 | 45.0441385 |
| C | 48.3871906 | 41.5098969 | 44.7524237 |
| C | 47.5662727 | 42.3095208 | 43.7382005 |
| C | 47.0263299 | 41.4337103 | 42.5989051 |
| C | 46.5367143 | 40.0570486 | 43.0517827 |

C 45.8808870 44.0027955 43.9229814
C 47.5621438 37.8904147 43.8814260
N 47.7122917 39.3668819 43.7265871
O 46.4664484 42.9386125 44.4723662
O 46.1761132 44.5221666 42.8802500
O 44.6468124 38.9616089 44.1055658
H 44.8510687 41.0062948 44.0702112
H 43.8056385 36.7306472 45.4251491
H 45.9739687 40.4235256 46.4102589
H 49.4065786 41.3760247 44.3612514
H 48.4778550 42.0649380 45.6943251
H 47.8423040 41.2923633 41.8770262
H 46.2300813 41.9601693 42.0555146
H 44.4887231 38.5429056 44.9974151
H 48.4757689 37.5026229 44.3511608
H 46.6835862 37.6753152 44.4973384
H 47.4341232 37.4592511 42.8802976
H 48.1668183 43.1239014 43.3115677
H 46.2096527 39.4434893 42.2019197
H 48.4198599 39.5693428 45.7466954
H 48.5827649 39.5334624 43.1737534
H 46.0058926 32.5898077 50.9258299
H 49.7801755 37.5555656 48.8246241
H 44.5571156 40.0595804 50.4259458
H 45.0790615 44.3576232 44.5704690
H 38.5229928 42.2017282 46.1189389

**TS2$^{L290F}$**

C 48.9983072 37.1340414 48.2278144
N 49.0672266 35.9561611 47.5090145
H 49.9038097 35.3807468 47.3564026
C 47.8541980 35.6944428 46.9932380
H 47.6227955 34.8387745 46.3624245
N 47.0030441 36.6533591 47.3501975
C 47.6973342 37.5588676 48.1203030
H 47.1963482 38.4232193 48.5372168
C 43.6395627 39.7186784 50.0204064
H 42.9942352 39.6233370 50.9044126
H 43.1742571 40.4645131 49.3592152
C 43.6393456 38.3964134 49.2473565
O 44.6877215 38.1497585 48.4982064
O 42.6687541 37.6556566 49.3279898
C 45.6351869 33.6448009 50.3562918
N 45.4614118 34.7800270 51.1243892
H 45.5757279 34.8707999 52.1381703
C 45.1163995 35.7845824 50.3008723
H 44.9306565 36.7948278 50.6398365
N 45.0450551 35.3604974 49.0435546
C 45.3732548 34.0203447 49.0602822
H 45.4016150 33.4430900 48.1406578
C 39.3642698 42.6942961 45.7243873
H 39.1979218 42.8939058 44.6508924
H 39.5429749 43.6744183 46.1962402
C 40.5801681 41.8265796 45.9531734
C 41.7377243 42.3999852 46.4991057
H 41.7434058 43.4681866 46.7303670
C 42.8663310 41.6238232 46.7772417
H 43.7472981 42.0812395 47.2356225
C 42.8580751 40.2563138 46.5107408
H 43.7168805 39.6340547 46.7516994
C 41.7194881 39.6776262 45.9429405
H 41.7195956 38.6087601 45.7442739
C 40.5992365 40.4531552 45.6595373
H 39.7284662 39.9712349 45.2090101
Fe 44.9353629 36.6261542 47.2662807
O 43.3322395 37.2095322 46.2478651
O 43.2250857 35.1702837 44.7525264
C 44.0917789 34.4588722 45.2998445
C 44.1993834 33.0125559 44.8692383
C 42.8097356 32.3730988 44.8226218
C 42.6858432 31.0158432 44.1155084
O 43.4484474 30.7621647 43.1493184
O 41.7535505 30.2791169 44.5261468
O 44.8712144 34.8846737 46.2419918
H 44.8883737 32.4802349 45.5393604
H 44.6303290 32.9851968 43.8535731
H 42.1464093 33.0590472 44.2705505
H 42.3815231 32.2765908 45.8317059
C 45.6824835 39.3968653 44.7851019
C 46.6451353 39.9180508 45.8133978
C 48.0152197 40.0738177 45.2418034
C 48.2988136 41.4910777 44.7024554
C 47.2425631 41.9675670 43.6972696
C 46.6617584 40.8411711 42.8229103
C 46.5636884 39.4649349 43.4969430
C 45.6854161 43.7660138 43.9302541
C 48.1570405 37.7166554 44.3218311
N 47.9508214 39.1588668 44.0259230

| | | | |
|---|---|---|---|
| O | 46.1717220 | 42.6330360 | 44.4437170 |
| O | 46.0230136 | 44.2944947 | 42.9052072 |
| O | 45.2262754 | 38.1066504 | 45.0963752 |
| H | 44.8286116 | 40.0933172 | 44.6507761 |
| H | 43.0868891 | 36.4416862 | 45.6450116 |
| H | 46.3745512 | 40.1349001 | 46.8428833 |
| H | 49.2854426 | 41.4963774 | 44.2151183 |
| H | 48.3574145 | 42.1922287 | 45.5440825 |
| H | 47.3185275 | 40.7363087 | 41.9457503 |
| H | 45.6804878 | 41.1449735 | 42.4323906 |
| H | 43.9803538 | 37.7937752 | 45.6246713 |
| H | 49.1307417 | 37.5960260 | 44.8108633 |
| H | 47.3336429 | 37.3720447 | 44.9511561 |
| H | 48.1455482 | 37.1806339 | 43.3655559 |
| H | 47.6735960 | 42.7297049 | 43.0368169 |
| H | 46.2894871 | 38.6995316 | 42.7605144 |
| H | 48.8389581 | 39.7452936 | 45.8919001 |
| H | 48.6830577 | 39.4409763 | 43.3363813 |
| H | 45.9774480 | 32.6968813 | 50.7714702 |
| H | 49.8239827 | 37.5486713 | 48.8060956 |
| H | 44.6172977 | 40.0780888 | 50.3412267 |
| H | 44.9178012 | 44.1790849 | 44.5846292 |
| H | 38.4424643 | 42.2615820 | 46.1131036 |

**IC2$^{L290F}$**

| | | | |
|---|---|---|---|
| C | 49.0983038 | 37.1027463 | 48.3098775 |
| N | 49.1627960 | 35.9143117 | 47.6120898 |
| H | 49.9961468 | 35.3340957 | 47.4648774 |
| C | 47.9458246 | 35.6519248 | 47.0992839 |
| H | 47.7141621 | 34.7948540 | 46.4701474 |
| N | 47.1008684 | 36.6188770 | 47.4368945 |
| C | 47.7958020 | 37.5265228 | 48.1976999 |
| H | 47.3007546 | 38.3961197 | 48.6106858 |
| C | 43.7000337 | 39.7949337 | 50.1007948 |
| H | 43.1545783 | 39.5295871 | 51.0185739 |
| H | 43.1374406 | 40.6174369 | 49.6408232 |
| C | 43.6126586 | 38.6399949 | 49.1008093 |
| O | 44.6444316 | 38.3071302 | 48.4210526 |
| O | 42.4716257 | 38.1389672 | 48.9831466 |
| C | 45.6611269 | 33.7468414 | 50.2681764 |
| N | 45.5135290 | 34.8871331 | 51.0319100 |
| H | 45.6325263 | 34.9748782 | 52.0439408 |
| C | 45.1885608 | 35.8967278 | 50.1952326 |
| H | 45.0399472 | 36.9188555 | 50.5214053 |
| N | 45.1079681 | 35.4720209 | 48.9447091 |
| C | 45.4041993 | 34.1282320 | 48.9694676 |
| H | 45.4174206 | 33.5433144 | 48.0531676 |
| C | 39.3831653 | 42.5045621 | 45.6323820 |
| H | 39.1875742 | 42.7870607 | 44.5835197 |
| H | 39.6841496 | 43.4260578 | 46.1556392 |
| C | 40.5063189 | 41.4983463 | 45.7497521 |
| C | 41.6422799 | 41.8568060 | 46.4875699 |
| H | 41.6916876 | 42.8600940 | 46.9124648 |
| C | 42.6977426 | 40.9633032 | 46.6807280 |
| H | 43.5609291 | 41.2615792 | 47.2806481 |
| C | 42.6425057 | 39.6942101 | 46.1068355 |
| H | 43.4428851 | 38.9764136 | 46.2527692 |
| C | 41.5425413 | 39.3403905 | 45.3249532 |
| H | 41.5454791 | 38.3651723 | 44.8370235 |
| C | 40.4753338 | 40.2240464 | 45.1563684 |
| H | 39.6274795 | 39.9213917 | 44.5367082 |
| Fe | 45.0206371 | 36.7493127 | 47.0198011 |
| O | 42.9979202 | 36.4750380 | 47.1227585 |
| O | 43.0687402 | 35.2792184 | 44.8629376 |
| C | 44.0536592 | 34.5933718 | 45.2059322 |
| C | 44.1631022 | 33.1606280 | 44.7117209 |
| C | 42.8009069 | 32.4699095 | 44.7386424 |
| C | 42.6833565 | 31.0996676 | 44.0551104 |
| O | 43.4332453 | 30.8320405 | 43.0822285 |
| O | 41.7658669 | 30.3571212 | 44.4922054 |
| O | 44.9882778 | 35.0204568 | 45.9910039 |
| H | 44.9155210 | 32.6324595 | 45.3145567 |
| H | 44.5374400 | 33.1780790 | 43.6733378 |
| H | 42.0822451 | 33.1292502 | 44.2243263 |
| H | 42.4296087 | 32.3671450 | 45.7699635 |
| C | 45.7603371 | 39.1435212 | 45.0900386 |
| C | 46.7722078 | 39.7293439 | 46.0183823 |
| C | 48.0633907 | 39.9867717 | 45.3185093 |
| C | 48.1558320 | 41.3962580 | 44.6962380 |
| C | 46.9899752 | 41.7176641 | 43.7504163 |
| C | 46.4504615 | 40.4977061 | 42.9793282 |
| C | 46.5343489 | 39.1651878 | 43.7324297 |
| C | 45.2630958 | 43.3427441 | 43.9486545 |
| C | 48.3260002 | 37.6073109 | 44.4918846 |
| N | 47.9823692 | 39.0105983 | 44.1524019 |
| O | 45.9234398 | 42.3407979 | 44.5389183 |
| O | 45.4328996 | 43.7418697 | 42.8282121 |
| O | 45.2830820 | 37.8776447 | 45.4905213 |

| | | | |
|---|---|---|---|
| H | 44.8907034 | 39.8205010 | 44.9847573 |
| H | 42.6378022 | 36.2188941 | 46.2397912 |
| H | 46.5475011 | 39.9942270 | 47.0452007 |
| H | 49.1003631 | 41.4722686 | 44.1366541 |
| H | 48.1984259 | 42.1418208 | 45.5004483 |
| H | 47.0401071 | 40.3972180 | 42.0550614 |
| H | 45.4153469 | 40.6862107 | 42.6631599 |
| H | 42.5182413 | 37.1203571 | 47.7124720 |
| H | 49.3286788 | 37.5854718 | 44.9327999 |
| H | 47.5767478 | 37.2223650 | 45.1841571 |
| H | 48.3029442 | 37.0258824 | 43.5635233 |
| H | 47.3058148 | 42.4688347 | 43.0167140 |
| H | 46.2665289 | 38.3327186 | 43.0715650 |
| H | 48.9675442 | 39.7696100 | 45.9054536 |
| H | 48.6394432 | 39.3170020 | 43.3977522 |
| H | 45.9688123 | 32.7904139 | 50.6908386 |
| H | 49.9257161 | 37.5255154 | 48.8797216 |
| H | 44.6973354 | 40.1326252 | 50.3826119 |
| H | 44.5399543 | 43.7902176 | 44.6304910 |
| H | 38.4408762 | 42.1433485 | 46.0443082 |

**TS3$^{L290F}$**

| | | | |
|---|---|---|---|
| C | 49.1493712 | 37.1745760 | 48.2825619 |
| N | 49.2230930 | 35.9894613 | 47.5803145 |
| H | 50.0553947 | 35.4056269 | 47.4481027 |
| C | 48.0114217 | 35.7336567 | 47.0460393 |
| H | 47.7847383 | 34.8733757 | 46.4189281 |
| N | 47.1624246 | 36.6979574 | 47.3717668 |
| C | 47.8484869 | 37.6002892 | 48.1478212 |
| H | 47.3444326 | 38.4656255 | 48.5622977 |
| C | 43.6717485 | 39.8195804 | 50.1312514 |
| H | 43.1225370 | 39.5619080 | 51.0486555 |
| H | 43.1076547 | 40.6277197 | 49.6464371 |
| C | 43.6452929 | 38.6427073 | 49.1459646 |
| O | 44.6726283 | 38.4329108 | 48.4140952 |
| O | 42.5664443 | 38.0105957 | 49.0903251 |
| C | 45.6486538 | 33.7628350 | 50.2002744 |
| N | 45.5211927 | 34.9104298 | 50.9574655 |
| H | 45.6668335 | 35.0043939 | 51.9642181 |
| C | 45.1663393 | 35.9108331 | 50.1209780 |
| H | 45.0301021 | 36.9357641 | 50.4448729 |
| N | 45.0442452 | 35.4770927 | 48.8764295 |
| C | 45.3492534 | 34.1342929 | 48.9064525 |
| H | 45.3431154 | 33.5426016 | 47.9941595 |
| C | 39.3571873 | 42.6136088 | 45.5964123 |
| H | 39.1522652 | 42.8415091 | 44.5360413 |
| H | 39.5989534 | 43.5722656 | 46.0829696 |
| C | 40.5344366 | 41.6807944 | 45.7657826 |
| C | 41.6494996 | 42.1371653 | 46.4798237 |
| H | 41.6551457 | 43.1652112 | 46.8473909 |
| C | 42.7334735 | 41.2960266 | 46.7439842 |
| H | 43.5733843 | 41.6600279 | 47.3410354 |
| C | 42.7230148 | 39.9850581 | 46.2764738 |
| H | 43.5268904 | 39.2989903 | 46.5192277 |
| C | 41.6420695 | 39.5298626 | 45.5176743 |
| H | 41.6650886 | 38.5030886 | 45.1534489 |
| C | 40.5549355 | 40.3658301 | 45.2667174 |
| H | 39.7171379 | 39.9872121 | 44.6750796 |
| Fe | 45.0644655 | 36.8521206 | 46.9937387 |
| O | 43.0299753 | 36.9896521 | 46.7119987 |
| O | 42.9953925 | 35.1492629 | 44.9894654 |
| C | 44.0631673 | 34.5307017 | 45.2487102 |
| C | 44.1750940 | 33.0925383 | 44.7610548 |
| C | 42.8195605 | 32.3923991 | 44.7771063 |
| C | 42.6971856 | 31.0397436 | 44.0667814 |
| O | 43.4461848 | 30.7858019 | 43.0884365 |
| O | 41.7727792 | 30.2939076 | 44.4854967 |
| O | 45.0410852 | 35.0367335 | 45.8958671 |
| H | 44.9300395 | 32.5731635 | 45.3693209 |
| H | 44.5617018 | 33.1061265 | 43.7265604 |
| H | 42.0964567 | 33.0595492 | 44.2800822 |
| H | 42.4468226 | 32.2713245 | 45.8059665 |
| C | 45.5924329 | 39.2962294 | 44.7486775 |
| C | 46.5046020 | 39.7146844 | 45.8154813 |
| C | 47.9011744 | 39.7585250 | 45.2916392 |
| C | 48.1902938 | 41.2186692 | 44.8575958 |
| C | 47.1711388 | 41.7666051 | 43.8439469 |
| C | 46.6030533 | 40.7114266 | 42.8731534 |
| C | 46.4640927 | 39.3084093 | 43.4787184 |
| C | 45.4716501 | 43.4261112 | 43.9864166 |
| C | 48.1098150 | 37.4830381 | 44.1318040 |
| N | 47.8402940 | 38.9479335 | 44.0105871 |
| O | 46.0873228 | 42.4019391 | 44.5953114 |
| O | 45.7123859 | 43.8335945 | 42.8833368 |
| O | 45.5672869 | 38.0599394 | 45.4445728 |
| H | 44.6112614 | 39.7727370 | 44.6204402 |
| H | 42.7523451 | 36.2246869 | 46.1237332 |
| H | 46.1839523 | 40.0058265 | 46.8134769 |

| | | | |
|---|---|---|---|
| H | 49.1950830 | 41.2380584 | 44.4100469 |
| H | 48.2212378 | 41.8631053 | 45.7440389 |
| H | 47.2827594 | 40.6398138 | 42.0113361 |
| H | 45.6413992 | 41.0567256 | 42.4704475 |
| H | 42.5819887 | 37.1202039 | 47.5826777 |
| H | 49.1052689 | 37.3577501 | 44.5732006 |
| H | 47.3431791 | 37.0169615 | 44.7482050 |
| H | 48.0936941 | 37.0751439 | 43.1139396 |
| H | 47.6419250 | 42.5587331 | 43.2489759 |
| H | 46.1689286 | 38.5740188 | 42.7195065 |
| H | 48.6939003 | 39.3693056 | 45.9434576 |
| H | 48.5745169 | 39.3074871 | 43.3562620 |
| H | 45.9584188 | 32.8110262 | 50.6317517 |
| H | 49.9679615 | 37.5886595 | 48.8712146 |
| H | 44.6658852 | 40.1688494 | 50.4101253 |
| H | 44.7171496 | 43.8729228 | 44.6338388 |
| H | 38.4376936 | 42.2095029 | 46.0198470 |

**PC$^{L290F}$**

| | | | |
|---|---|---|---|
| C | 49.0022004 | 37.2046107 | 48.2428015 |
| N | 49.0882256 | 36.0486889 | 47.4915907 |
| H | 49.9288238 | 35.4817692 | 47.3351699 |
| C | 47.8725532 | 35.7878924 | 46.9695374 |
| H | 47.6487879 | 34.9390038 | 46.3255290 |
| N | 47.0059264 | 36.7209016 | 47.3485683 |
| C | 47.6915765 | 37.6104107 | 48.1444347 |
| H | 47.1722392 | 38.4490633 | 48.5950854 |
| C | 43.5651227 | 39.8160895 | 50.2173035 |
| H | 43.0337374 | 39.5958921 | 51.1559261 |
| H | 42.9933479 | 40.6094810 | 49.7171777 |
| C | 43.5163461 | 38.5989457 | 49.2795798 |
| O | 44.5801074 | 38.2388685 | 48.6889054 |
| O | 42.3827630 | 38.0770967 | 49.1266487 |
| C | 45.6632342 | 33.5992830 | 50.4905483 |
| N | 45.5106659 | 34.7338377 | 51.2638868 |
| H | 45.6376338 | 34.8244662 | 52.2754001 |
| C | 45.1596509 | 35.7425513 | 50.4425033 |
| H | 44.9998811 | 36.7650736 | 50.7673940 |
| N | 45.0661462 | 35.3186338 | 49.1908402 |
| C | 45.3846633 | 33.9804805 | 49.1994597 |
| H | 45.3994534 | 33.4063851 | 48.2753610 |
| C | 39.4376287 | 42.5964420 | 45.6243694 |
| H | 39.2418299 | 42.8288329 | 44.5626728 |
| H | 39.6720317 | 43.5560873 | 46.1129496 |
| C | 40.6278683 | 41.6763768 | 45.7934680 |
| C | 41.7817919 | 42.1794199 | 46.4098481 |
| H | 41.8009499 | 43.2260346 | 46.7217484 |
| C | 42.8877891 | 41.3624046 | 46.6624017 |
| H | 43.7516621 | 41.7712201 | 47.1936536 |
| C | 42.8661002 | 40.0214543 | 46.2841975 |
| H | 43.6891831 | 39.3488321 | 46.5159647 |
| C | 41.7422713 | 39.5210166 | 45.6203157 |
| H | 41.7406807 | 38.4689112 | 45.3362297 |
| C | 40.6350651 | 40.3329760 | 45.3771389 |
| H | 39.7688997 | 39.9099191 | 44.8617739 |
| Fe | 44.9017892 | 36.5270406 | 47.4380496 |
| O | 43.0262906 | 36.9663256 | 46.7815966 |
| O | 42.9998677 | 35.2385297 | 44.9601744 |
| C | 44.0028230 | 34.5307114 | 45.2708567 |
| C | 44.1082763 | 33.1503863 | 44.6411121 |
| C | 42.7637074 | 32.4301094 | 44.7074028 |
| C | 42.6561341 | 31.0613627 | 44.0269812 |
| O | 43.3976702 | 30.8023520 | 43.0445412 |
| O | 41.7561004 | 30.3047065 | 44.4778246 |
| O | 44.8922628 | 34.8820984 | 46.1052530 |
| H | 44.9033924 | 32.5949880 | 45.1564767 |
| H | 44.4118108 | 33.2535097 | 43.5853492 |
| H | 42.0109859 | 33.0766646 | 44.2263192 |
| H | 42.4297098 | 32.3128351 | 45.7503011 |
| C | 45.5785253 | 39.7568120 | 44.3653261 |
| C | 46.3772523 | 39.8712892 | 45.5879423 |
| C | 47.8395845 | 39.8766127 | 45.1685568 |
| C | 48.2927415 | 41.3091953 | 44.8476854 |
| C | 47.4127757 | 42.0172154 | 43.8028925 |
| C | 46.8907765 | 41.0850785 | 42.6923127 |
| C | 46.5473155 | 39.6769449 | 43.1957535 |
| C | 45.7310029 | 43.7117241 | 43.9457263 |
| C | 47.8993180 | 37.6343870 | 43.9281909 |
| N | 47.8048231 | 39.1268894 | 43.8488372 |
| O | 46.2957367 | 42.6427972 | 44.5153909 |
| O | 46.0395261 | 44.1993005 | 42.8919450 |
| O | 45.6612770 | 38.6745705 | 45.2900518 |
| H | 44.5989205 | 40.2261922 | 44.2429233 |
| H | 42.7402083 | 36.2592502 | 46.1183016 |
| H | 46.0650912 | 40.4293540 | 46.4734770 |
| H | 49.3241863 | 41.2704152 | 44.4684871 |
| H | 48.3214221 | 41.8997909 | 45.7715417 |
| H | 47.6684131 | 41.0005765 | 41.9208222 |

H 46.0238712 41.5419899 42.1947663
H 42.4603258 37.2038608 47.5555425
H 48.8744159 37.3878781 44.3670382
H 47.0885518 37.2381398 44.5397129
H 47.8367501 37.2550355 42.9010299
H 47.9773135 42.8330915 43.3338649
H 46.2524151 39.0019492 42.3815630
H 48.5347047 39.3709120 45.8513095
H 48.6271752 39.4222652 43.2760300
H 46.0195170 32.6478297 50.8853475
H 49.8284182 37.6133628 48.8244827
H 44.5680937 40.1654448 50.4623895
H 44.9400041 44.1074238 44.5827583
H 38.5098894 42.1987680 46.0357356

**$TS_{OH}^{L290F}$**

C 49.1262641 37.1706301 48.3251413
N 49.2230635 35.9922330 47.6171281
H 50.0722439 35.4402874 47.4572422
C 48.0136979 35.6909901 47.1119078
H 47.8118717 34.8207733 46.4912358
N 47.1384361 36.6222472 47.4695235
C 47.8112611 37.5538568 48.2251591
H 47.2865700 38.4074399 48.6378724
C 43.5782339 39.6773071 50.1785003
H 43.0080405 39.5284351 51.1066734
H 43.0424624 40.4415951 49.5971047
C 43.5304547 38.4000091 49.3290172
O 44.5857846 38.1346647 48.6173630
O 42.4929063 37.7440190 49.3082095
C 45.6692062 33.6202923 50.4726450
N 45.5147818 34.7476076 51.2551961
H 45.6315902 34.8282618 52.2689733
C 45.1665249 35.7625486 50.4411379
H 44.9944214 36.7746429 50.7859370
N 45.0776085 35.3532076 49.1838417
C 45.3927019 34.0130055 49.1840818
H 45.3933205 33.4449797 48.2576469
C 39.4328589 42.5702378 45.7082483
H 39.2758025 42.7556294 44.6305868
H 39.6408920 43.5542995 46.1592346
C 40.6298249 41.6769799 45.9586987
C 41.8047310 42.2607284 46.4544773
H 41.8303296 43.3402475 46.6255070
C 42.9207189 41.4828926 46.7721993
H 43.8111948 41.9527765 47.1994414
C 42.8820600 40.1018617 46.5960156
H 43.7241940 39.4752663 46.8753861
C 41.7372643 39.5102912 46.0575714
H 41.7230844 38.4292259 45.9008093
C 40.6248077 40.2875975 45.7357395
H 39.7400295 39.8028286 45.3162648
Fe 45.0634196 36.6478651 47.4050101
O 44.9397967 37.7710353 45.8316092
O 44.3903818 35.6751274 44.3138670
C 44.4714147 34.7403967 45.1384106
C 44.3405659 33.2910096 44.6823220
C 42.8968336 32.7588585 44.6622573
C 42.7253340 31.3418363 44.0506534
O 43.4470063 31.0182238 43.0734970
O 41.8221654 30.6217941 44.5481056
O 44.6818406 34.9360521 46.3978049
H 44.9458316 32.6729917 45.3625681
H 44.7547323 33.2055909 43.6675055
H 42.2740476 33.4363108 44.0497045
H 42.4616978 32.7479260 45.6726352
C 45.2795487 39.9707828 44.0659977
C 46.0386992 39.6264864 45.3319652
C 47.5009375 39.6302064 45.0063616
C 48.1209173 41.0557968 44.9617122
C 47.4683088 42.0184352 43.9573857
C 46.9621886 41.3144321 42.6861753
C 46.3817667 39.9282193 42.9684055
C 45.9040468 43.8310775 44.0947736
C 47.3550212 37.6312582 43.4998238
N 47.5099681 39.1091100 43.5875081
O 46.3744267 42.7102307 44.6487570
O 46.2974129 44.3437184 43.0818397
O 44.1735313 39.1462759 43.8161779
H 44.9132614 41.0069392 44.1504141
H 44.8494478 37.0786605 45.1102806
H 45.7174109 39.9692528 46.3118758
H 49.1788058 40.9368918 44.6843123
H 48.1053205 41.4936483 45.9674870
H 47.8080335 41.2201611 41.9916136
H 46.2223529 41.9393412 42.1691050
H 44.0644696 38.6485614 44.6571255
H 48.2372305 37.1734566 43.9659936

| | | | |
|---|---|---|---|
| H | 46.4537291 | 37.3243990 | 44.0317546 |
| H | 47.2924193 | 37.3588027 | 42.4412353 |
| H | 48.1879262 | 42.7969595 | 43.6704532 |
| H | 46.0431402 | 39.4259800 | 42.0535994 |
| H | 48.1210041 | 38.9870615 | 45.6472993 |
| H | 48.4127487 | 39.3515761 | 43.1156871 |
| H | 46.0194897 | 32.6660107 | 50.8659836 |
| H | 49.9419937 | 37.5952232 | 48.9102679 |
| H | 44.5718579 | 40.0447540 | 50.4349399 |
| H | 45.0925983 | 44.2379197 | 44.6981788 |
| H | 38.4929341 | 42.1844831 | 46.1029656 |

**$PC_{OH}^{L290F}$**

| | | | |
|---|---|---|---|
| C | 48.9888640 | 36.9631268 | 48.3430523 |
| N | 49.1054508 | 35.8320775 | 47.5632268 |
| H | 49.9712295 | 35.3319745 | 47.3388385 |
| C | 47.8758092 | 35.4934288 | 47.1207883 |
| H | 47.6882910 | 34.6453094 | 46.4657009 |
| N | 46.9716510 | 36.3462450 | 47.5842631 |
| C | 47.6500318 | 37.2651046 | 48.3495835 |
| H | 47.1203804 | 38.0734342 | 48.8359145 |
| C | 43.6182453 | 39.6559326 | 50.0371856 |
| H | 43.0038704 | 39.5275567 | 50.9386768 |
| H | 43.1244141 | 40.4161573 | 49.4126808 |
| C | 43.6290956 | 38.3517602 | 49.2326928 |
| O | 44.6014181 | 38.1765985 | 48.3773051 |
| O | 42.7112848 | 37.5483173 | 49.3722261 |
| C | 45.6704470 | 33.4679204 | 50.7836620 |
| N | 45.4824126 | 34.5693619 | 51.5915065 |
| H | 45.5841528 | 34.6338868 | 52.6066709 |
| C | 45.0796970 | 35.5861262 | 50.8023105 |
| H | 44.8520954 | 36.5779559 | 51.1778810 |
| N | 44.9984658 | 35.1984436 | 49.5408857 |
| C | 45.3711354 | 33.8743281 | 49.5091045 |
| H | 45.3863642 | 33.3244450 | 48.5724548 |
| C | 39.3986125 | 42.5511684 | 45.6537909 |
| H | 39.2178314 | 42.7785707 | 44.5884570 |
| H | 39.6453221 | 43.5098204 | 46.1380536 |
| C | 40.5698358 | 41.6112082 | 45.8393415 |
| C | 41.7326809 | 42.1049969 | 46.4467410 |
| H | 41.7706981 | 43.1554861 | 46.7443027 |
| C | 42.8247070 | 41.2720690 | 46.7057518 |
| H | 43.7014333 | 41.6739641 | 47.2211163 |
| C | 42.7743726 | 39.9274743 | 46.3431867 |

| | | | |
|---|---|---|---|
| H | 43.5979841 | 39.2585731 | 46.5664599 |
| C | 41.6455419 | 39.4318284 | 45.6869814 |
| H | 41.6396255 | 38.3878379 | 45.3692022 |
| C | 40.5516969 | 40.2612518 | 45.4397496 |
| H | 39.6819668 | 39.8510904 | 44.9209365 |
| Fe | 44.8514025 | 36.2583259 | 47.6969135 |
| O | 46.1231510 | 38.9213889 | 46.3213617 |
| O | 44.3669717 | 36.0393362 | 45.3382442 |
| C | 44.3590789 | 34.8056851 | 45.6086136 |
| C | 44.2455860 | 33.7284942 | 44.5486227 |
| C | 42.9928769 | 32.8577714 | 44.7205239 |
| C | 42.8283173 | 31.7392276 | 43.6712986 |
| O | 43.2187240 | 31.9726944 | 42.5020033 |
| O | 42.2483536 | 30.6826007 | 44.0503804 |
| O | 44.4672687 | 34.4317418 | 46.8215852 |
| H | 45.1469834 | 33.1000938 | 44.6433385 |
| H | 44.2703495 | 34.1727117 | 43.5483691 |
| H | 42.1016140 | 33.5011190 | 44.6174652 |
| H | 42.9537259 | 32.4233002 | 45.7286114 |
| C | 45.5149192 | 40.0150091 | 44.2318101 |
| C | 46.3505360 | 40.0873049 | 45.5767770 |
| C | 47.8193472 | 40.1326221 | 45.1052485 |
| C | 48.3412776 | 41.5375684 | 44.7841846 |
| C | 47.4896508 | 42.2917719 | 43.7619090 |
| C | 47.0021729 | 41.3763894 | 42.6323775 |
| C | 46.5834013 | 39.9869196 | 43.1177171 |
| C | 45.7365259 | 43.9178846 | 43.9392799 |
| C | 47.7362535 | 37.8728466 | 43.9383103 |
| N | 47.7835271 | 39.3613180 | 43.8023809 |
| O | 46.3562102 | 42.8722529 | 44.4898850 |
| O | 46.0114690 | 44.4371098 | 42.8919565 |
| O | 44.7029469 | 38.8819656 | 44.2134032 |
| H | 44.8811673 | 40.9059325 | 44.1198213 |
| H | 45.6017850 | 39.0403392 | 47.1435429 |
| H | 46.1056888 | 40.9920430 | 46.1465759 |
| H | 49.3598887 | 41.4437605 | 44.3793732 |
| H | 48.4283226 | 42.1147604 | 45.7129526 |
| H | 47.8259386 | 41.2669118 | 41.9140345 |
| H | 46.1787973 | 41.8497854 | 42.0800351 |
| H | 44.9245773 | 38.3404992 | 44.9976227 |
| H | 48.6627473 | 37.5508183 | 44.4312102 |
| H | 46.8741781 | 37.5639890 | 44.5327045 |
| H | 47.6826036 | 37.4538567 | 42.9258875 |
| H | 48.0499585 | 43.1327107 | 43.3329368 |

| | | | |
|---|---|---|---|
| H | 46.2776311 | 39.3444862 | 42.2809300 |
| H | 48.4880553 | 39.6215874 | 45.8085912 |
| H | 48.6430715 | 39.5705659 | 43.2471985 |
| H | 46.0708088 | 32.5111642 | 51.1189159 |
| H | 49.8154750 | 37.4304012 | 48.8782495 |
| H | 44.6010663 | 40.0173783 | 50.3396538 |
| H | 44.9366502 | 44.2602810 | 44.5958092 |
| H | 38.4611967 | 42.1708840 | 46.0596268 |
